# Ancestral State Reconstruction under Localized Evolutionary Change: Error Patterns and Their Control by Tree Partitioning

**DOI:** 10.64898/2026.10.12.758557

**Authors:** Yuqi Cheng, Changqiao Chen, Deng-Ke Niu

**Affiliations:** MOE Key Laboratory for Biodiversity Science and Ecological Engineering, College of Life Sciences, Beijing Normal University, Beijing, 100875, China

**Author notes:** Correspondence to be sent to: Deng-Ke Niu, College of Life Sciences, Beijing Normal University, Beijing, 100875, China.

**Keywords:** Ancestral state reconstruction, model misspecification, evolutionary jumps, tree partitioning, Brownian motion, continuous traits, macroevolution, phylogenetic comparative methods

## Abstract

Localized evolutionary changes can distort ancestral state reconstruction (ASR) because conventional continuous-trait methods generally treat branchwise changes as draws from a shared background process. A disproportionately large change confined to a single branch therefore creates localized model misspecification, but how the resulting errors vary among reconstruction paradigms and phylogenetic positions remains poorly understood. Using controlled simulations on idealized and empirical mammalian phylogenies, we characterized the error patterns produced by single-branch large changes (SBLCs) across five Brownian-motion-based ASR implementations. On intact trees, errors were highly structured and method dependent: the recursive local method propagated distortion mainly upstream, whereas global methods produced multidirectional effects involving both SBLC boundary nodes, neighboring lineages, and descendants of the affected branch. We then evaluated patching, a conditional robustness procedure that partitions the phylogeny at a candidate branch, applies the same ASR method independently to the resulting subtrees, and estimates the upstream boundary node state by branch-length-weighted interpolation. Patching reduced errors near and away from the affected branch and lowered tree-wide error across trait-increasing and trait-decreasing SBLCs, a broad range of sensitivity analyses, and configurations with two co-occurring SBLCs, while introducing little additional error when no SBLC was present. Under the reference upsizing SBLC, for instance, it cut the maximum-likelihood reconstruction error at the upstream boundary node from 56.20 to 0.03 (mean true state, 4.98), matching the stable-process benchmark. In cetacean body-length evolution, patching candidate branches identified by a Lévy-process model generally shifted conventional estimates toward stable-process estimates and nearby fossil references. Under an Ornstein–Uhlenbeck process with strong post-SBLC mean reversion, however, neither patching nor stable-process reconstruction recovered downstream ancestral states once the phenotypic signature of the SBLC had largely disappeared from descendant tip values, revealing an identifiability limit of terminal-only inference. Localized evolutionary change therefore creates structured, method-dependent reconstruction errors, and tree partitioning provides a complementary diagnostic of the robustness of ancestral inferences to candidate branch-localized changes.

---

Ancestral state reconstruction (ASR) is widely used to infer the evolutionary histories of continuous traits on phylogenetic trees (Joy et al. 2016, Revell 2025, Revell 2026). Because ancestral states are unobserved, however, their estimates depend on both the assumed model of trait evolution and the way information from observed tips is transmitted through the phylogeny. This dependence becomes especially important when most of a tree is reasonably compatible with the fitted model but a small part contains an atypical evolutionary change.

Such localized departures present a model-misspecification problem that may not be adequately described by measures of average model fit across the entire tree (Pennell et al. 2015).

For the continuous-trait methods considered here, a useful distinction concerns how information is combined across the phylogeny. Local, contrast-based procedures use recursive calculations involving neighboring branches and nodes. Felsenstein-type independent-contrast logic provides the conceptual basis for this approach (Felsenstein 1985). In recursive implementations, the estimate at each internal node becomes an input to the calculation at its parent, so information is transmitted stepwise from the tips toward the root. Global, model-based estimators instead fit an explicit stochastic model to the full tree and observed tip data. Maximum-likelihood reconstruction under Brownian motion (BM), generalized least-squares reconstruction based on a phylogenetic covariance matrix, and phylopars-type joint estimation are examples of this approach (Martins and Hansen 1997, Schluter et al. 1997, Goolsby 2017, Goolsby et al. 2017). The Bayesian implementation examined here likewise uses a tree-wide evolutionary model but estimates posterior distributions of ancestral states and integrates uncertainty in model parameters (Huelsenbeck and Bollback 2001, Pagel et al. 2004). Despite these differences, all of the implementations considered here interpret branchwise changes relative to a common background model. A localized change that departs strongly from this background may therefore affect ancestral-state estimates under any of them, although the affected nodes and the direction and extent of the resulting errors may depend on how each method combines information across the phylogeny.

Under the BM assumptions shared by the conventional implementations examined here, stochastic variation among branches is allowed, but branchwise changes are expected to arise from a shared distribution after accounting for branch length. A disproportionately large phenotypic change concentrated on one branch can therefore create localized model misspecification even when the same background model remains an adequate approximation for most of the phylogeny. Rapid evolutionary episodes, abrupt shifts, and punctuated changes are increasingly recognized as important features of trait evolution (Uyeda et al. 2011, Landis et al. 2013, Landis and Schraiber 2017, Gao and Wu 2022a, Brennan et al. 2024). Here, we use localized evolutionary change for this broader problem and define a single-branch large change (SBLC) as the operational case examined in this study: a disproportionately large phenotypic change concentrated on one phylogenetic branch, irrespective of its underlying mechanism.

Alternative evolutionary models can explicitly accommodate departures from Gaussian BM. Lévy-process models represent discrete jumps on individual branches, whereas stable-process reconstruction uses a heavy-tailed distribution of branchwise changes that allows occasional changes of large magnitude (Landis et al. 2013, Elliot and Mooers 2014, Landis and Schraiber 2017). These approaches provide principled ways to reconstruct ancestral states under evolutionary processes that include abrupt or heavy-tailed change. A complementary question remains, however: how does a candidate localized change affect estimates from a specified conventional ASR method on the intact tree, and can its influence be limited while retaining that method? This question is relevant both for assessing the robustness of conventional reconstructions and for evaluating candidate branches suggested by biological or fossil evidence or by diagnostic or process-based analyses.

We address this question through a conditional robustness analysis based on tree partitioning. Given a candidate branch, the procedure—here termed patching—partitions the phylogeny at that branch and applies the same conventional ASR method independently to the resulting tree components. Comparing the intact-tree and patched reconstructions therefore reveals how strongly an ancestral-state estimate depends on information transmitted across the candidate branch. Because the cut position is supplied rather than inferred by the patch procedure, patching is not a method for detecting localized changes and does not replace an explicit evolutionary process model. Using controlled simulations with known ancestral states, we first characterize how SBLC-induced errors on intact trees vary among five BM-based reconstruction methods and across phylogenetic positions. We then test whether patching reduces these errors under a broad range of conditions, compare the remaining errors with those from stable-process reconstruction as an external benchmark, and apply the procedure to candidate SBLCs in cetacean body-length evolution.

## Materials and Methods

### Operationalization of SBLCs and the Patch Procedure

For the simulation analyses, SBLCs were imposed on prespecified focal branches, whereas candidate SBLC-bearing branches in the empirical analysis were identified using the Lévy-process procedure described below. The patch framework therefore presupposed a designated candidate branch and was not itself used to detect SBLCs.

Given a candidate SBLC-bearing branch from node *I* to node *J*, the patch procedure removes that branch and partitions the original phylogeny into the descendant subtree rooted at *J* (*T*_1_) and the complementary subtree (*T*_2_). Here, *I* and *J* are the upstream and downstream endpoints of the affected branch, respectively; *H* is the parent of *I*; and *J*^1^ is the root of the unaffected sister lineage descending from *I* (Fig. 1). Nodes *H*, *I*, *J*, and *J*^1^ constitute the four focal positions in the primary node-wise analyses. The same ASR method is applied independently to *T*_1_ and *T*_2_.

**Figure 1.**
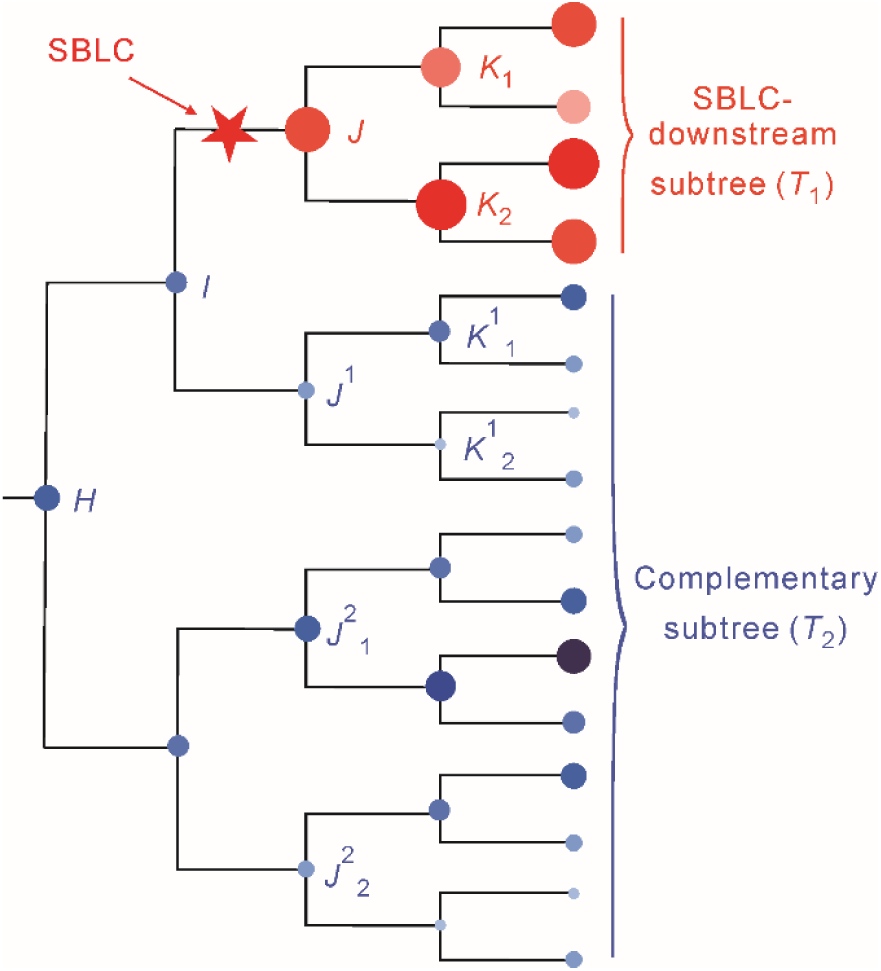
Conceptual illustration of a single-branch large change (SBLC), the patch procedure, and the SBLC-centered node-coordinate system. The star marks an SBLC on the branch from *I* to *J*, which separates the SBLC-descendant subtree (*T*1) from the complementary subtree (*T*2). Circle size schematically indicates relative trait magnitude and is not drawn to a quantitative scale. The patch procedure removes the *I*–*J* branch and applies the same ancestral state reconstruction method independently to the resulting subtrees; the upstream boundary node *I* is estimated separately by branch-length-weighted interpolation, as described in the text. The schematic does not represent the exact simulation topology. Primary simulations used a 128-tip balanced ultrametric tree with *T*1 = 8, whereas propagation analyses used a 4,096-tip balanced tree with *T*1 = 64. Node labels define positions relative to the SBLC-bearing branch: *H*, *G*, *F*, *E*, and *D* denote successive upstream positions from *I* toward the root; *K*, *L*, *M*, *N*, and *O* denote successive downstream positions from *J* within *T*1; and superscripts denote lateral positions away from the SBLC-bearing lineage. Nodes *H*, *I*, *J*, and *J*^1^ are the four focal positions in the primary analyses. This coordinate system is also used to locate the propagation-analysis nodes in Figure 5; labels therefore denote corresponding topological positions rather than fixed node identities across trees.

Removal of the *I*–*J* branch changes *I* from a bifurcating internal node to a boundary point. Consequently, a direct estimate for *I* is not available from the standard ASR applied to either resulting subtree. Its patched state is therefore estimated from the adjacent nodes *H* and *J*^1^ in *T*_2_ by branch-length-weighted interpolation:

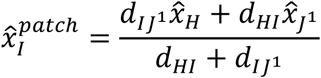

where *x̂_H_* and *x̂_J_*1 are the subtree-based estimates for nodes *H* and *J*^1^, and *d_HI_* and *d_IJ_*1 are the corresponding branch lengths. Weighting each estimate by the distance to the opposite adjacent node gives greater weight to the node closer to *I*, while excluding information from the SBLC-bearing branch.

The final patched reconstruction combines the independently obtained estimates for *T*_1_ and *T*_2_ with the interpolated estimate for *I*. Across trees and SBLC placements, corresponding nodes are identified by their topological positions in the SBLC-centered coordinate system shown in Figure 1, rather than by invariant node identities.

### Conventional ASR Methods and External Benchmark

We evaluated the patch procedure using five conventional continuous-trait ASR implementations, all based on Brownian-motion assumptions but spanning contrasting reconstruction paradigms: local contrast-based reconstruction (PIC-local; Felsenstein 1985), maximum-likelihood reconstruction (ML-BM; Schluter et al. 1997), generalized least-squares reconstruction (GLS-BM; Martins and Hansen 1997), phylopars-type joint reconstruction (Rphy-BM; Goolsby et al. 2017), and Bayesian MCMC reconstruction (Bayes-MCMC; Huelsenbeck and Bollback 2001, Pagel et al. 2004). Within each method, the same reconstruction procedure and method-specific settings were used for the intact tree and the partitioned subtrees. Thus, differences between unpatched and patched estimates reflect reconstruction on the intact phylogeny versus independently on the resulting subtrees, rather than a change in the ASR model or algorithm.

Stable-process reconstruction, which was not subjected to patching, was included separately as an external benchmark (Stable; Elliot and Mooers 2014, Castiglione et al. 2018). Its heavy-tailed evolutionary model accommodates occasional changes of large magnitude and therefore provides a process-based reference for assessing whether patching reduces the residual errors of conventional methods to a comparable scale.

The selected methods were intended to represent contrasting reconstruction paradigms rather than to exhaust all available ASR algorithms. Implementation details for the five conventional methods and the Stable benchmark, including software, method-specific options, MCMC configurations, and posterior summaries, are provided in Supplementary Methods Section S1.

### Primary Simulations on a Fixed Idealized Phylogeny

For each simulation condition, we generated 100 independent continuous-trait histories on the same fixed, balanced ultrametric phylogeny with 128 terminal taxa. Branch lengths were scaled to a unit root-to-tip height, and the prespecified focal branch from node *I* to node *J* subtended a descendant subtree (*T*_1_) containing eight terminal taxa. Holding the topology and branch-length structure constant allowed variation in reconstruction error to be evaluated across independently simulated trait histories.

Background trait evolution followed BM. For a branch of length *t_ab_* connecting parent node *a* to descendant node *b*,

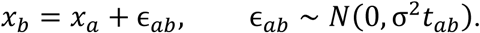

We fixed the root state at *x*_root_ = 5 and the BM rate parameter at *σ*^2^ = 1. Conditional on the fixed root, a terminal state under the background process therefore had marginal variance 1 and standard deviation 1; the root value placed the background simulations on a predominantly positive trait scale. All terminal and internal-node states were retained as the known values against which reconstructed states were evaluated.

Under the background BM process, the transition along the focal branch would be

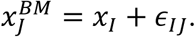

To impose an SBLC while retaining the stochastic BM increment on this branch, we instead generated

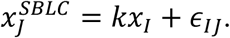

Equivalently,

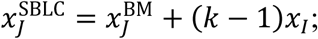

that is, the imposed SBLC adds (*k* − 1)*x_I_* to the value at *J* that would otherwise have arisen under background BM. The primary simulations examined an upsizing SBLC with *k* = 16 and a reciprocal downsizing SBLC with *k* = 1/16. After the modified transition, all descendants of *J* evolved under the same BM process, with *x*^SBLC^ as the initial state. The imposed large change was therefore localized to the *I*–*J* branch, whereas subsequent evolution within *T*1 followed the background model.

For each simulated trait history, the five conventional ASR methods were applied to the intact phylogeny and, for the patched analyses, independently to the resulting subtrees.

Stable-process reconstruction was applied separately to the intact phylogeny as an external benchmark. The resulting estimates were compared with the recorded internal-node states in the error analyses described below.

To characterize the spatial extent of SBLC-induced reconstruction error, we conducted an additional propagation analysis on a balanced ultrametric tree with 4,096 terminal taxa, with *T*_1_ containing 64 taxa. Under the same BM background process, we examined both an upsizing SBLC (*k* = 16) and its reciprocal downsizing counterpart (*k* = 1/16) across four six-position series: upstream from *I* (*I*–*D*), downstream from *J* (*J*–*O*), along a depth-controlled lateral series (*J*^1^–*J*^6^), and progressively within the adjacent sister lineage (*J*^1^–*O*^1^; Fig. 1). Trait simulation and reconstruction procedures otherwise followed those of the primary simulations. Node-coordinate labels represent topological position classes rather than unique nodes. When a class contained multiple nodes, signed errors were averaged across all nodes in that class within each replicate before being summarized across the 100 replicates; no representative node was selected. Full node-class definitions and summarization procedures are provided in Supplementary Methods Section S2.1.

### Evaluation Metrics and Error Analysis

Signed reconstruction error was used as the primary node-wise metric because it retains the direction of estimation error and therefore distinguishes systematic overestimation from underestimation. For each reconstruction method, signed error at internal node *n* in replicate *r* was calculated as

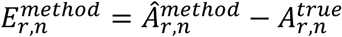

where 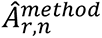 is the reconstructed ancestral state and 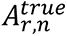 is the corresponding state recorded during simulation. Positive and negative 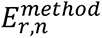 values indicate overestimation and underestimation, respectively.

For descriptive scale comparisons in the representative scenarios, we also calculated percentage relative signed error as 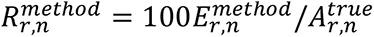. Because this ratio becomes unstable when the true state approaches zero, substantive conclusions were based on unscaled signed errors.

The primary node-wise analyses considered the four prespecified focal nodes *H*, *I*, *J*, and *J*^1^ (Fig. 1). For each method, simulation condition, and focal node, directional bias was summarized as the mean signed error across the 100 independent trait histories. Bootstrap 95% confidence intervals were obtained from 10,000 nonparametric resamples of the replicate-level errors, using the 2.5th and 97.5th percentiles of the bootstrap distribution. An interval excluding zero was interpreted as evidence of detectable directional bias. Errors were analyzed separately at each focal node; nodes within the same trait history were not treated as independent observations. The independently simulated trait histories, rather than individual nodes, constituted the sampling units.

Because the unpatched and patched estimates for each node were obtained from the same simulated trait history, their difference was evaluated on a paired, replicate-wise basis:

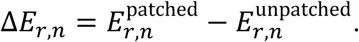

For each method, condition, and focal node, mean Δ*E_r_*_,*n*_ was calculated across the 100 replicates, and its 95% confidence interval was obtained from 10,000 bootstrap resamples of the replicate-level differences. An interval excluding zero indicated a detectable patch-induced change in signed error. Because Δ*E_r_*_,*n*_ measures a signed change rather than error magnitude, its direction was interpreted relative to the corresponding unpatched error; movement toward zero indicated correction of directional bias.

To assess reconstruction accuracy across the phylogeny, we additionally calculated mean absolute error (MAE) within each trait history for three sets of internal nodes: all 127 internal nodes, including the root; the 123 nodes remaining after exclusion of the four focal nodes; and the 113 nodes remaining after exclusion of a broader set of 14 nodes surrounding the SBLC-bearing branch. For each node set, patch performance was summarized by the paired contrast

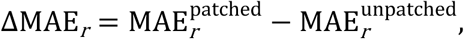

so that negative values indicated lower tree-wide reconstruction error after patching. The exclusion analyses assessed whether an overall reduction in error was confined to the immediate SBLC region and whether partitioning caused deterioration farther from the affected branch. Confidence intervals were obtained by resampling complete trait histories, thereby preserving both dependence among nodes within each history and the pairing between patched and unpatched reconstructions. Full node-set definitions and computational details are provided in Supplementary Methods Section S2.2.

### Sensitivity Analyses

To assess whether the effects of SBLCs and the performance of patching were robust beyond the primary reference setting described above, we conducted controlled sensitivity analyses on idealized phylogenies and complementary analyses on an empirical mammalian phylogeny. In the controlled analyses, one major feature was varied at a time while the remaining settings were held at their reference values. These analyses varied SBLC magnitude, total phylogenetic size, the size of the SBLC-descendant subtree (*T*_1_), tree topology, and the background evolutionary model. The magnitude analysis included *k* = 1, representing the absence of an imposed SBLC. Unless otherwise stated, the remaining analyses were conducted under both the upsizing condition (*k* = 16) and the reciprocal downsizing condition (*k* = 1/16). Because absolute node identities were not comparable across trees and SBLC placements, focal positions were defined using the SBLC-centered coordinate system shown in Figure 1.

Sensitivity to the background evolutionary process was evaluated by simulating traits under variance-normalized Pagel’s λ-transformed BM, Ornstein–Uhlenbeck (OU), and early-burst (EB) models following the covariance parameterizations of Huang et al. (2026). These models were used only as alternative data-generating processes; the ASR procedures and their method-specific settings remained unchanged. Model parameters were scaled so that terminal marginal variance matched the BM reference, allowing differences in reconstruction error to be interpreted primarily in relation to covariance structure and the temporal distribution of evolutionary variance rather than differences in overall trait scale. We excluded extreme parameter values under which phylogenetic covariance among tip states was absent or nearly absent. In such limiting cases, the tip data provide little information for reconstructing internal-node states, and the analysis would mainly reflect a general loss of phylogenetic signal rather than the effects of SBLCs or patching.

Complementary analyses were conducted on the DNA-only maximum-clade-credibility tree from the VertLife mammalian phylogeny of Upham et al. (2019), which contained 4,099 mammalian terminal taxa after removal of the non-mammalian outgroup. These analyses examined contrasting combinations of local topology within *T*_1_ and its surrounding phylogenetic context, post-SBLC evolutionary duration under strong OU attraction, and the spatial interaction of nearby and phylogenetically distant pairs of SBLCs. The paired-SBLC analyses additionally tested whether jointly partitioning the phylogeny at both affected branches retained the corrective effect observed for a single SBLC. Detailed parameter settings, variance-normalization procedures, tree processing, branch-selection criteria, and simulation implementations are provided in Supplementary Methods Section S3.

### Empirical Cetacean Analysis

For the empirical analysis of cetacean body-length evolution, we used the time-calibrated mitochondrial phylogeny of Zurano et al. (2019), inferred independently of body-length data, for extant-species ASR; the total-group cetacean metatree of Lloyd and Slater (2021) to place fossils; and mean total body lengths from Burin et al. (2023) and the publicly available Back to the Water dataset (Park et al. 2022). Body length was analyzed as mean total length in meters; when multiple directly measured records were available for a taxon, their arithmetic mean was used, and imputed records were excluded. After taxonomic matching, the molecular tree was pruned to 79 extant species with directly measured body lengths.

Topological and temporal screening of fossil placements retained 15 fossil reference points. Fossil data were not used in candidate SBLC identification or ASR, but only as external paleontological references for evaluating reconstructed internal-node states.

Using the 79-tip molecular tree and extant-species body lengths, we fitted the jump-normal Lévy-process model of Landis et al. (2013) to identify candidate branches showing evidence of localized change. Branches were ranked by the absolute value of a branch-normalized signal-to-noise ratio (|SNR|). Among the five highest-ranking branches, we retained those whose central 90% posterior credible interval for jump magnitude excluded zero. The retained branches were treated as candidate rather than confirmed SBLCs and used as cut points individually and in combination. We compared conventional ASR on the intact tree with patched reconstructions and with Stable-process reconstruction as an external benchmark; estimates were also compared with the 15 fossil references. Full procedures for taxon matching, fossil mapping, Lévy-process inference and diagnostics, candidate selection, and empirical ASR are provided in Supplementary Methods Section S4.

### Software Implementation and Reproducibility

The patch procedure was implemented in the open-source R package patchASR (version 0.1.1; Cheng 2026), which partitions a phylogeny at one or more user-specified candidate branches, applies a selected ASR implementation independently to the resulting tree components, and merges the estimates onto the original phylogeny. The archived release includes automated unit and numerical regression tests against version-controlled reference outputs and passed R CMD checks on Linux, macOS, and Windows; implementation and validation details are provided in Supplementary Methods Section S5. All materials required to reproduce the analyses and reported outputs are available in the accompanying reproducibility archive (https://doi.org/10.5061/dryad.47d7wm3wt).

## Results

### ASR Error Patterns Under Representative SBLCs

We first characterized reconstruction errors under a representative upsizing SBLC (*k* = 16) imposed on the branch from *I* to *J*. The analysis focused on four positions surrounding the affected branch: the upstream node *H*, the upstream and downstream branch endpoints *I* and *J*, and the adjacent lateral node *J*^1^ (Fig. 1).

At node *I*, all five methods showed positive mean signed errors, ranging from 4.60 to 56.20, and all 95% bootstrap confidence intervals excluded zero (Fig. 2a; Supplementary Table S1). The largest error occurred under ML-BM, for which the mean signed error was 56.20, compared with a mean true state of 4.98. Positive errors persisted at *H* but were smaller than at *I* for every method, ranging from 3.30 to 34.52, indicating attenuation along the ancestral path. Bayes-MCMC showed detectable positive errors at both upstream nodes, but they were substantially smaller than those of the other methods.

**Figure 2.**
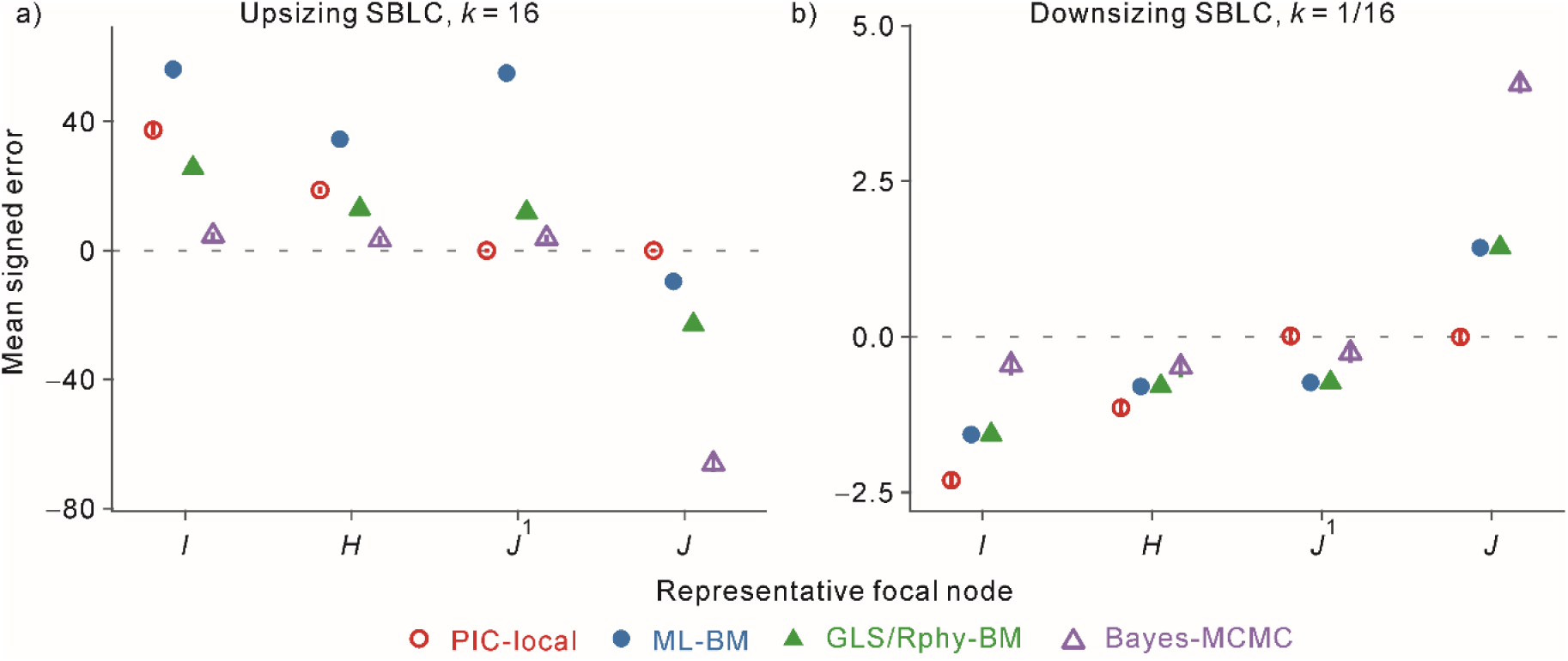
Method- and position-dependent ancestral state reconstruction (ASR) errors under representative upsizing and downsizing single-branch large changes (SBLCs). Mean signed errors, calculated as reconstructed minus true ancestral state, are shown for conventional unpatched reconstructions at four focal nodes surrounding the SBLC-bearing branch *I*–*J* (Figure 1). Panel (a) shows an upsizing SBLC with *k* = 16, and panel (b) shows the reciprocal downsizing SBLC with *k* = 1/16. Points show means across 100 independently simulated trait histories, and error bars show 95% bootstrap confidence intervals for the mean. GLS-BM and Rphy-BM produced identical mean errors under these conditions and are plotted together as GLS/Rphy-BM. Confidence intervals excluding zero indicate detectable positive or negative directional bias. Many intervals are narrower than the plotting symbols; numerical values are provided in Supplementary Table S1.

The position-dependent contrast was especially pronounced at the lateral node *J*^1^ and the downstream boundary node *J*. At *J*^1^, the PIC-local mean error was close to zero and its confidence interval included zero, whereas all global methods showed positive mean errors; the largest was 55.00 under ML-BM. At *J*, PIC-local again showed no detectable directional bias, but all global methods showed negative mean errors. Bayes-MCMC had the largest error at this node, with a mean of −66.11. Thus, PIC-local errors were confined mainly to the ancestral path above the affected branch, whereas the global methods showed positive errors at upstream and lateral positions together with error in the opposite direction at the immediate descendant-side boundary.

The reciprocal downsizing SBLC (*k* = 1/16) produced a broadly reversed pattern. Method–node combinations with positive mean errors under upsizing generally showed negative errors under downsizing, whereas the negative errors at *J* became positive and combinations with no detectable displacement remained close to zero (Fig. 2b; Supplementary Table S1). Absolute errors were substantially smaller than under upsizing. Bayes-MCMC retained its distinctive spatial profile, showing comparatively small upstream and lateral errors but the largest absolute error at *J*. The approximate reversal in error direction indicates that the observed patterns tracked the direction of the imposed SBLC, whereas their spatial extent depended on the reconstruction method.

### Error Reduction by Patching Under Representative SBLCs

We next compared conventional reconstructions on the intact phylogeny with patched reconstructions from the same simulated trait histories. Figure 3 repeats the unpatched mean errors shown in Figure 2 and pairs them with the corresponding patched results, allowing the effect of tree partitioning to be evaluated directly.

**Figure 3.**
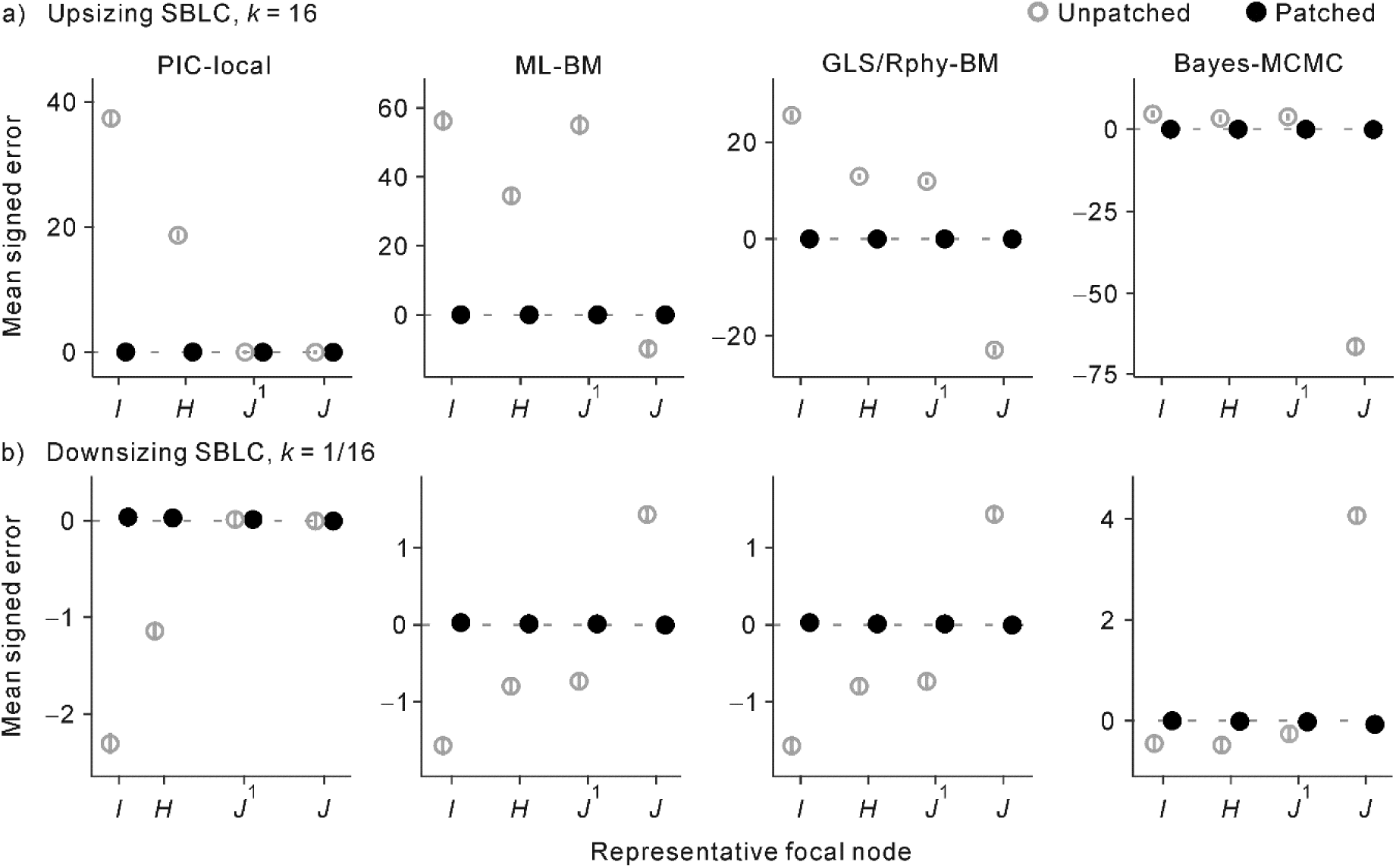
Reduction of directional ancestral state reconstruction (ASR) error by patching under representative single-branch large changes (SBLCs). Mean signed errors are shown before and after patching at four focal nodes surrounding the SBLC-bearing branch. Panel (a) shows an upsizing SBLC with *k* = 16, and panel (b) shows the reciprocal downsizing SBLC with *k* = 1/16. GLS-BM and Rphy-BM produced identical mean errors and are shown together as GLS/Rphy-BM. Open gray circles show conventional reconstructions on the intact phylogeny, and filled black circles show patched reconstructions. Points are means across 100 independently simulated trait histories, and error bars show 95% bootstrap confidence intervals for the mean. Note that y-axis scales differ among panels; many intervals for the patched estimates are narrower than the plotting symbols. Focal nodes follow Figure 1. The unpatched values are repeated from Figure 2 for direct comparison; numerical values for the unpatched and patched reconstructions are provided in Supplementary Tables S1 and S2, respectively.

Under the representative upsizing SBLC (*k* = 16), patching moved mean signed errors close to zero across methods and focal nodes (Fig. 3a; Supplementary Tables S1–S2). All 95% bootstrap confidence intervals for the patched conventional reconstructions included zero except that for Bayes-MCMC at node *J*, where a small negative mean error remained (−0.09). The downsizing SBLC (*k* = 1/16) showed the same corrective pattern despite its smaller unpatched errors (Fig. 3b). Patched mean errors again remained close to zero, with Bayes-MCMC at *J* providing the only confidence interval that excluded zero; its residual mean error was −0.07. Patching therefore reduced the directionally reversed errors generated by both increases and decreases in trait value.

Paired bootstrap comparisons showed that the positions affected by patching corresponded closely to the method-specific error patterns on the intact tree (Supplementary Table S3). For PIC-local, detectable changes were confined to the upstream nodes *I* and *H*, whereas the patched–unpatched differences at *J*^1^ and *J* were exactly zero under both SBLC directions. The global methods showed detectable changes at all four focal nodes. For Bayes-MCMC, mean changes at *I*, *H*, and *J*^1^ were comparatively modest, whereas the changes at *J* were 66.02 under upsizing and −4.13 under downsizing—the largest absolute patch-induced changes among the method–node combinations in their respective scenarios. Thus, the spatial distribution of patch-induced changes mirrored the distinct error footprints of the local and global reconstruction methods.

### Comparable Residual Errors from Patched and Stable-Process Reconstructions

To place the corrected errors in context, we compared the patched conventional reconstructions with Stable-process reconstruction. Stable served as an external benchmark because its heavy-tailed evolutionary model can accommodate occasional changes of large magnitude without requiring the affected branches to be specified in advance. This comparison was descriptive: we asked whether patching reduced the residual errors of conventional BM-based reconstructions to a scale comparable to that obtained under Stable, rather than formally testing the superiority of one approach over another.

Under the representative upsizing SBLC, the patched conventional methods and Stable all showed small residual mean signed errors (Fig. 4a; Supplementary Table S2). Across the four focal nodes, these errors ranged from −0.09 to 0.04, and the 95% bootstrap confidence intervals included zero for every estimate except Bayes-MCMC at node *J*. The reciprocal downsizing scenario produced similarly small residual errors, ranging from −0.08 to 0.08, although the pattern was not a strict mirror image of that under upsizing (Fig. 4b).

**Figure 4.**
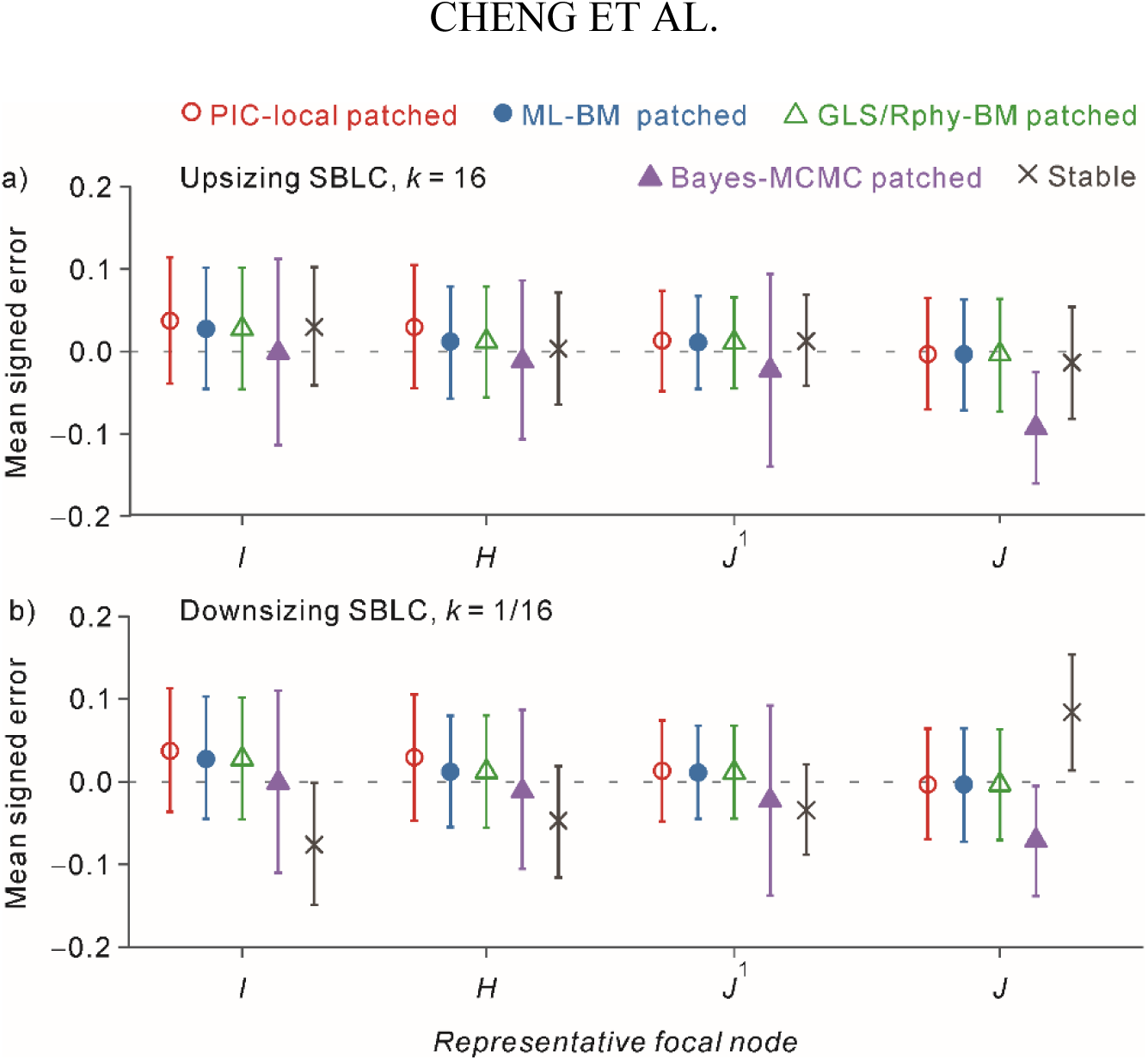
Residual signed errors after patching compared with Stable-process reconstruction under representative single-branch large-change (SBLC) scenarios. Mean signed ASR errors are shown at four focal nodes for the patched conventional methods and for Stable-process reconstruction fitted to the intact phylogeny. Panel (a) shows an upsizing SBLC with *k* = 16, and panel (b) shows a downsizing SBLC with *k* = 1/16. The patched estimates are the same as those shown in Figure 3 but are replotted over a narrower y-axis range to show the residual errors more clearly. Points represent mean signed errors, and error bars indicate 95% bootstrap confidence intervals for the means across simulation replicates. Values closer to zero indicate smaller directional reconstruction errors. Focal-node labels follow Figure 1. Numerical values are reported in Supplementary Table S2, and percentage relative signed errors are reported in Supplementary Table S4.

Confidence intervals excluded zero for Bayes-MCMC at *J* and for Stable at *I* and *J*. These residual deviations were nevertheless minor relative to the corresponding unpatched errors, placing the patched conventional reconstructions and Stable within a broadly comparable residual-error range.

The magnitude of the correction was particularly clear at node *I* under the upsizing SBLC. The mean true ancestral state was 4.98, whereas ML-BM reconstruction on the intact tree produced a mean signed error of 56.20. After patching, the corresponding error fell to 0.03, the same value to two decimal places as that obtained under Stable. Because Stable was included as a process-based external benchmark and the primary inferential comparison concerned patched versus unpatched reconstruction within each conventional method, we did not conduct additional node-wise tests to rank the patched methods against Stable.

### Spatial Propagation of SBLC-Induced Reconstruction Error

To determine how far SBLC-induced reconstruction errors extended across the phylogeny, we examined six successive position classes in four directions from the affected branch on a balanced 4,096-tip tree: upstream from *I* toward the root, downstream from *J* within *T*_1_, laterally across lineages of controlled depth, and progressively within the adjacent sister lineage. Figure 5 shows the upsizing condition (*k* = 16); the reciprocal downsizing condition (*k* = 1/16) is shown in Supplementary Figure S1.

**Figure 5.**
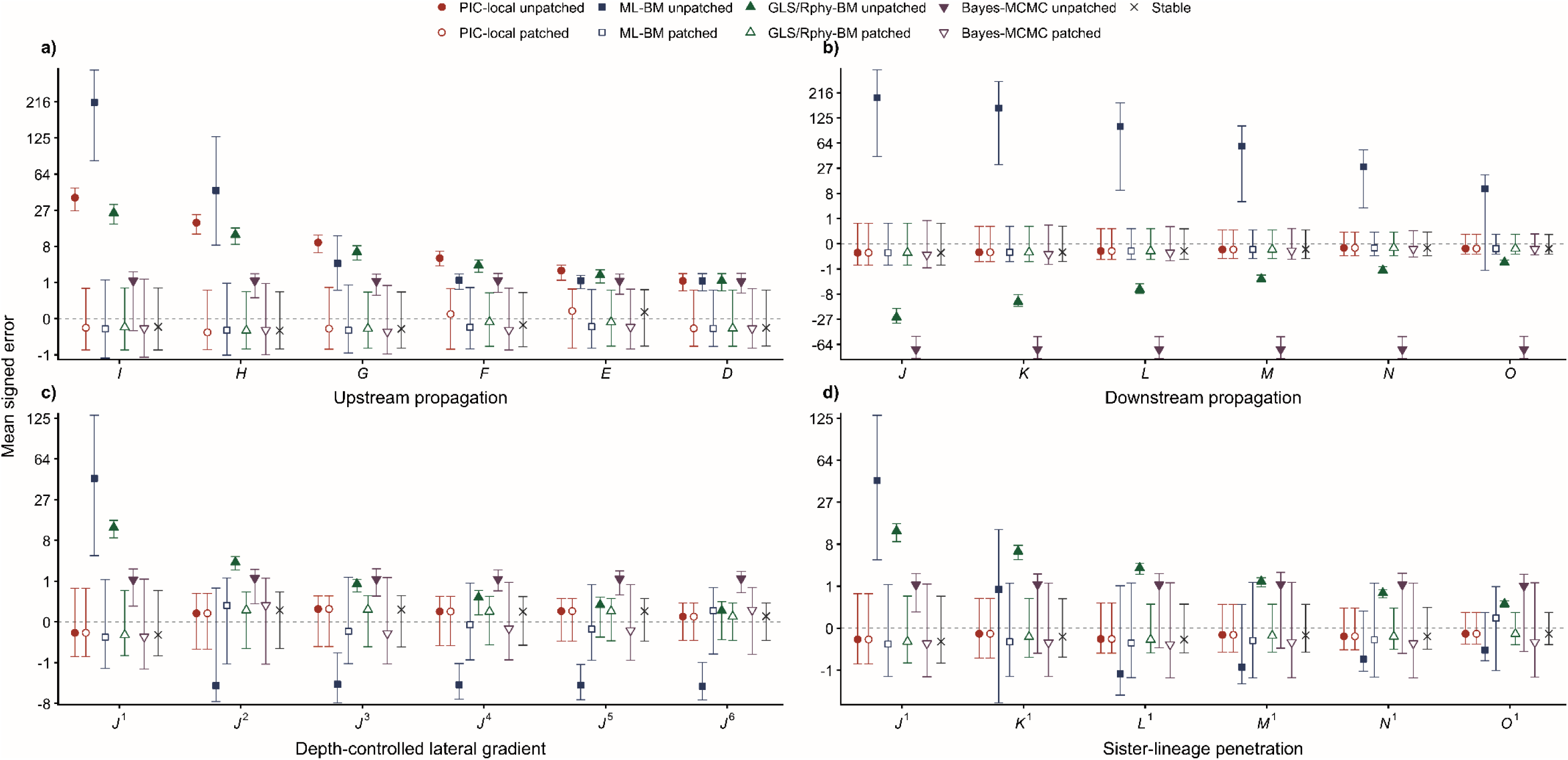
Spatial propagation of SBLC-induced reconstruction error under an upsizing SBLC. Results are shown for an upsizing SBLC (*k* = 16) on a balanced ultrametric tree with 4,096 terminal taxa and an SBLC-descendant subtree (*T*1) containing 64 taxa. Mean signed reconstruction errors were evaluated across six topological position classes in four directions relative to the SBLC-bearing *I*–*J* branch: (a) upstream from *I* toward the root (*I*–*D*); (b) downstream from *J* within *T*1 (*J*–*O*); (c) across a depth-controlled lateral series (*J*^1^–*J*^6^); and (d) progressively within the adjacent sister lineage (*J*^1^–*O*^1^). Filled and open symbols denote unpatched and patched conventional reconstructions, respectively; crosses denote Stable-process reconstruction on the intact phylogeny.

Under the upsizing SBLC, unpatched errors showed strongly method- and direction-dependent spatial patterns (Fig. 5). Upstream, the positive errors of PIC-local, ML-BM, and GLS/Rphy-BM generally attenuated with increasing topological separation from the affected branch. The decline was especially pronounced for ML-BM under the 4,096-tip tree, whose mean signed error fell from 214.36 at *I* to 1.15 at *D*, although it remained above zero at the most distant position examined. Bayes-MCMC showed much smaller upstream errors but little attenuation across the series. Downstream, PIC-local remained essentially unaffected, whereas the global methods differed markedly: the positive ML-BM error decreased from 195.42 at *J* to 10.42 at *O*, the negative GLS/Rphy-BM error declined in magnitude from −24.83 to −0.40, and the Bayes-MCMC error remained close to −74 throughout the series. Under this larger-tree configuration, the ML-BM error at *J* was positive rather than negative as in the primary 128-tip analysis, showing that even the direction of distortion can depend on phylogenetic context.

The lateral analyses revealed a related but nonmonotonic pattern. In the depth-controlled lateral series, the largest ML-BM and GLS/Rphy-BM errors occurred at *J*^1^, the position closest to the SBLC. GLS/Rphy-BM errors then declined progressively toward zero, whereas ML-BM showed a sharp reduction and reversal from positive error at *J*^1^ to smaller but persistent negative errors from *J*^2^ onward (Fig. 5c). Within the adjacent sister lineage, ML-BM likewise declined sharply and reversed direction, but subsequently approached zero with increasing penetration into the lineage; GLS/Rphy-BM errors decreased more steadily.

Bayes-MCMC retained small positive errors with little systematic distance dependence in both lateral series, whereas PIC-local remained close to zero (Fig. 5d).

The reciprocal downsizing SBLC (*k* = 1/16) produced broadly reversed but substantially smaller errors (Supplementary Fig. S1). Across both SBLC directions, spatial propagation on the intact tree was strongly method and direction dependent. PIC-local transmitted the influence primarily upstream, whereas the global methods could also show downstream and lateral distortion; attenuation with increasing topological separation was evident in some method–direction combinations but weak, nonmonotonic, or absent in others.

Patching substantially reduced these errors across all four position series. Patched estimates generally remained close to zero and showed little systematic variation with topological separation, and Stable-process estimates showed a similarly limited spatial pattern. Thus, although intact-tree reconstructions produced structured spatial error patterns, patching largely removed them across the examined topological positions.

Axis labels denote topological position classes rather than individual nodes. Within each replicate, signed errors were averaged across all nodes assigned to each class; for example, *K*^1^ and *J*^2^ each contained two nodes. Points show the means of these replicate-level class averages across the 100 simulated trait histories, and error bars indicate their 95% bootstrap confidence intervals. The vertical axis uses a sign-preserving cube-root transformation, with tick labels expressed in the original signed-error units. For PIC-local, patched and unpatched errors were identical at all downstream and lateral positions; the horizontally offset symbols therefore represent the same values. Node relationships follow Figure 1, and full node-class definitions are provided in Supplementary Methods Section S2. Numerical results are reported in Supplementary Table S5.

### Patching Reduced Tree-Wide Reconstruction Error

We next returned to the primary simulations on the fixed 128-tip balanced phylogeny to determine whether the improvements near the SBLC-bearing branch translated into lower average error across the tree and whether restricting each reconstruction to the tips within its own tree component caused a net loss of accuracy elsewhere. Across all reconstruction methods and both SBLC directions, the mean paired MAE contrasts, calculated as patched minus unpatched MAE, were negative when all 127 internal nodes were included, and their 95% bootstrap confidence intervals lay entirely below zero (Supplementary Fig. S2). When the analysis was repeated after excluding the four focal nodes and after excluding the broader set of 14 local nodes, the contrasts generally shifted toward zero but remained negative, with confidence intervals below zero for every method and SBLC direction. Thus, the tree-wide reduction in reconstruction error was not driven solely by nodes near the affected branch.

Under the reference conditions, patching also reduced mean error among the remaining, more distant nodes, with no evidence of a net loss in reconstruction accuracy outside the local neighborhood of the SBLC.

### Robustness and Limits of Patch Performance

The preceding analyses showed that SBLCs produced method- and position-dependent reconstruction errors under both upsizing (*k* = 16) and reciprocal downsizing (*k* = 1/16) conditions, and that patching reduced focal, spatially propagated, and tree-wide errors in both directions. We next tested whether this corrective performance persisted across departures from the primary reference setting: a balanced 128-tip phylogeny evolving under BM, with root-to-tip height normalized to 1 and an eight-tip SBLC-descendant subtree. Detailed results are provided in Supplementary Results Section S1.

We first varied SBLC magnitude while holding the remaining features of the reference setting fixed (Fig. 6). On intact trees, the errors of unpatched ASR methods generally increased as the upsizing multiplier increased above *k* = 1, but their spatial distributions differed among methods. PIC-local errors increased primarily at the upstream nodes, whereas the global methods also showed lateral displacement and opposing error at the descendant-side boundary node *J*. ML-BM followed this broader pattern at large SBLC magnitudes but showed several nonmonotonic responses at intermediate values. Patching substantially reduced these magnitude-dependent errors across methods and focal nodes. Reciprocal downsizing SBLCs produced broadly reversed but smaller unpatched errors, with a comparable corrective response to patching (Supplementary Fig. S3).

**Figure 6.**
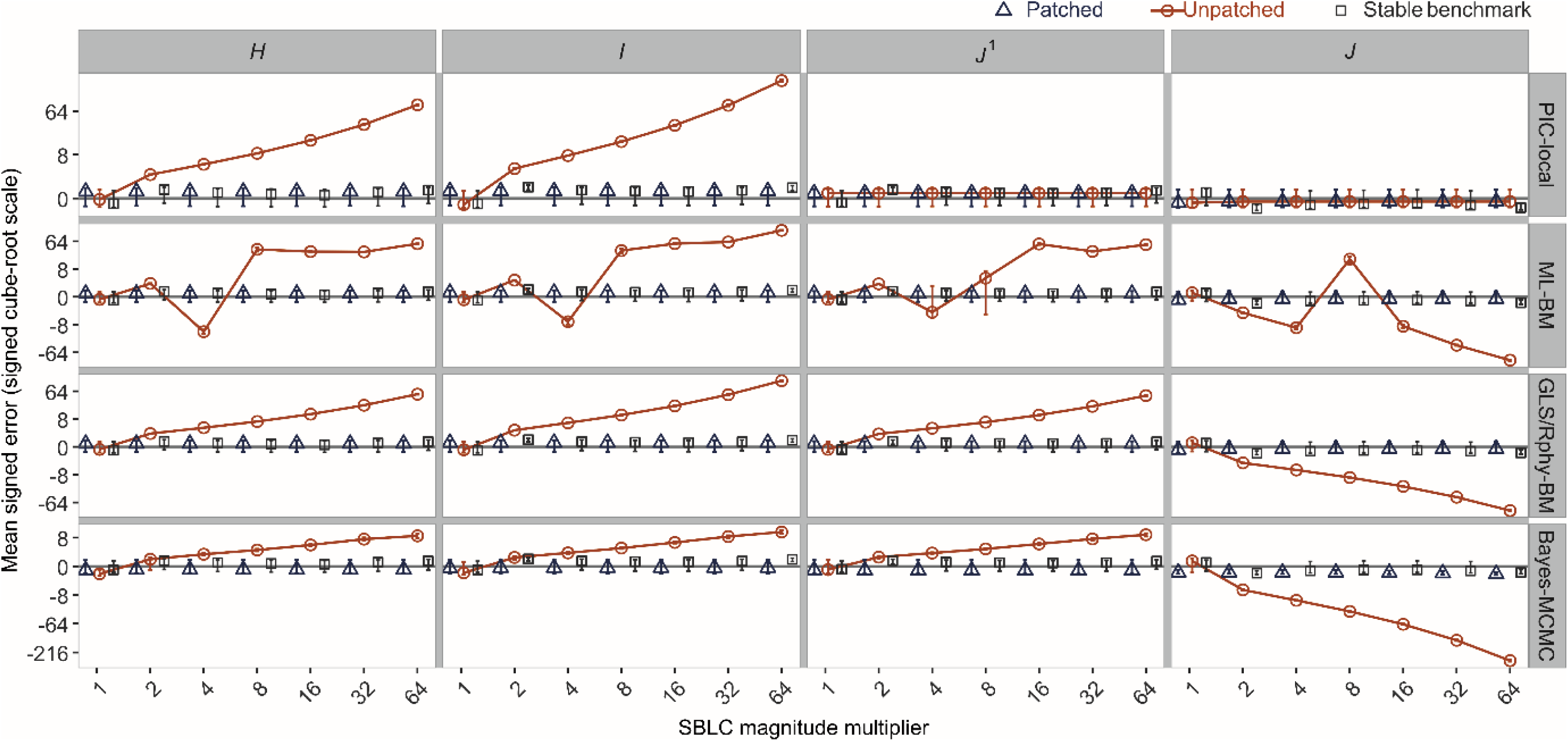
Effects of SBLC magnitude on ancestral-state reconstruction error before and after patching. Mean signed reconstruction errors are shown across upsizing single-branch large-change (SBLC) multipliers from *k* = 1, representing no imposed SBLC, to *k* = 64. Columns correspond to the focal nodes defined in Figure 1. Circles, triangles, and squares denote unpatched, patched, and Stable-process reconstructions, respectively; Stable was included as an external benchmark. Points show means across 100 independent simulation replicates, and error bars indicate 95% bootstrap confidence intervals for the means. Positive and negative values indicate overestimation and underestimation, respectively. GLS-BM and Rphy-BM produced identical mean errors and are displayed together as GLS/Rphy-BM.

Importantly, the *k* = 1 baseline provided a direct negative control for partitioning at a branch without an imposed SBLC. Patched errors closely matched the corresponding unpatched errors and generally remained near zero across methods and focal nodes; errors from Stable-process reconstruction were similarly small. Thus, under the reference conditions, unnecessary partitioning did not introduce appreciable additional directional error. This result suggests that patching is unlikely to substantially increase reconstruction error when a candidate branch represents a false-positive identification.

Because the complete upsizing and downsizing magnitude gradients established broadly reciprocal responses, the remaining sensitivity results presented below and in Supplementary Results Section S1 use *k* = 16 as a common strong-perturbation reference. All corresponding analyses were also conducted under *k* = 1/16 and produced generally smaller, directionally reversed unpatched errors with a comparable response to patching. Their complete graphical and tabulated results are retained in Repository Appendix A of the archived reproducibility materials.

Patch performance remained robust to changes in the amount and arrangement of phylogenetic information. Increasing total tree size from 64 to 1,024 terminal taxa, varying the size of the SBLC-descendant subtree (*T*_1_) from 1 to 16 taxa, and replacing the balanced tree with random Yule topologies altered some method- and node-specific errors without changing the principal spatial patterns (Supplementary Figs. S4–S6). ML-BM was the most sensitive to realized topology. Nevertheless, patching consistently reduced errors across these variations, with patched estimates generally remaining close to zero and similar in scale to the Stable-process benchmark.

Changing the background evolutionary process revealed a more consequential distinction. Under variance-normalized *λ*-BM and early-burst models, changes in the distribution and sharing of evolutionary variance had only modest effects on the principal error patterns, and patching remained effective across the examined gradients (Supplementary Figs. S7–S8). Under Ornstein–Uhlenbeck evolution, however, stronger mean reversion progressively erased the phenotypic signature of the immediate post-SBLC state from the descendant tips (Supplementary Figs. S9–S10). This loss of signal reduced unpatched distortion at upstream and lateral nodes but made the state at *J*, and other ancestral states within *T*_1_, increasingly difficult to reconstruct. Patching continued to restrict the remaining SBLC influence across the cut boundary, but it could not recover ancestral information that was no longer retained in the tip observations. Under the strongest mean reversion, substantial error at *J* therefore persisted under both patched conventional reconstruction and Stable-process reconstruction.

The robustness to topology also extended to BM simulations on the fixed 4,099-species empirical mammalian phylogeny. Across the four combinations of balanced-like and ladder-like topology in *T*_1_ and its surrounding phylogenetic context, realized topology altered some method- and node-specific errors, particularly under ML-BM, but did not change the principal corrective response to patching (Supplementary Figs. S11–S12).

Beyond these topological contrasts, the empirical mammalian phylogeny allowed us to test whether the OU-related information-loss mechanism persisted across differences in post-SBLC evolutionary duration. Across placements representing short, intermediate, and long durations, mean signed error at *J* became progressively more negative (Fig. 7; Supplementary Fig. S13). After patching, the global methods converged toward the duration-dependent error scale shared by PIC-local and Stable-process reconstruction, with mean errors of approximately −5, −43, and −60 from the short- to long-duration placements. A similar duration-dependent loss of accuracy occurred at internal nodes within *T*_1_ (Supplementary Table S6). Outside *T*_1_, however, the relationship with duration was reversed: shorter post-SBLC durations preserved a stronger anomalous signal in the descendant tips and consequently produced greater upstream distortion in several unpatched methods, whereas patching kept upstream errors close to zero.

**Figure 7.**
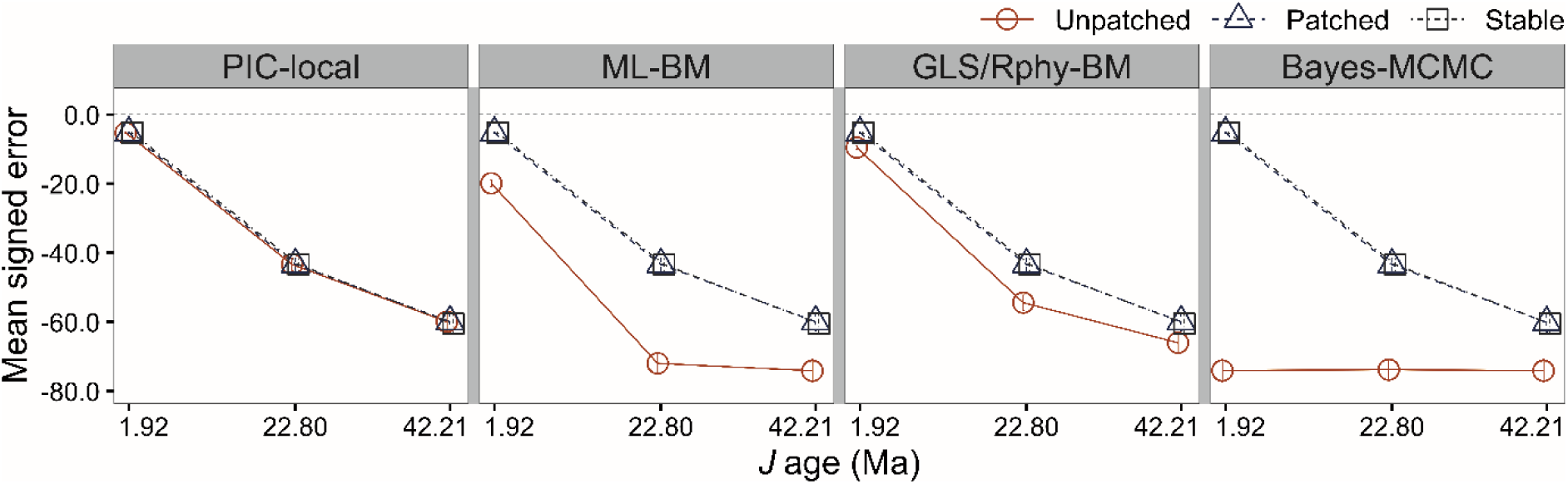
Effect of post-SBLC evolutionary duration on reconstruction error at the descendant-side boundary node under Ornstein–Uhlenbeck evolution. Mean signed errors at node *J* are shown for three SBLC placements on the empirical mammalian phylogeny, with *J* ages of 1.92, 22.80, and 42.21 Ma representing short, intermediate, and long periods of post-SBLC evolution, respectively. Each placement had an eight-tip descendant subtree, and the upsizing SBLC used *k* = 16. Circles, triangles, and squares denote unpatched, patched, and Stable-process reconstructions, respectively. Points show means across 100 simulation replicates, and error bars indicate 95% bootstrap confidence intervals for the means. The corresponding phylogenetic placements and local tree configurations are shown in Supplementary Figure S13.

Finally, using the same empirical-topology simulation framework, we examined configurations containing two SBLCs. Three placements with similar descendant-subtree sizes and topologies were selected to form one nearby and two phylogenetically distant pairs. Adding a second SBLC altered unpatched error in several method-, node-, and placement-specific comparisons. The nearby pair produced greater cumulative distortion than the corresponding single-SBLC scenarios in some cases, but this pattern was not uniform across methods or nodes (Supplementary Figs. S14–S15). Phylogenetic proximity could therefore strengthen the combined influence of two SBLCs, but did not determine it by itself. When both affected branches were jointly patched, the principal upstream and descendant-side errors were reduced toward the Stable-process benchmark in both nearby and distant configurations. At the adjacent lateral node evaluated for the two distant configurations, patched errors were likewise generally comparable to the benchmark. Joint patching therefore retained the corrective effect observed for a single SBLC across both nearby and distant configurations.

Taken together, the sensitivity analyses distinguish robustness from recoverability.

Patching consistently limited distortion transmitted across candidate branches, but under an Ornstein–Uhlenbeck process with strong post-SBLC mean reversion, none of the examined approaches recovered ancestral states within the SBLC-descendant subtree once information about those states had largely disappeared from its tip values.

### Cetacean ASR Responses to Candidate-Branch Patching

Following branch ranking by |SNR| and application of the prespecified posterior-support criterion, three branches were retained as candidate SBLCs. In decreasing order of |SNR|, they were designated SBLC-*φ*, SBLC-*ψ*, and SBLC-*ω*. All three represented inferred increases in body length, and their phylogenetic locations are shown in Supplementary Figure S16. Their posterior jump estimates and other candidate characteristics are reported in Supplementary Results Section S2.1.

Consistent with the upsizing simulations, patching generally reduced ancestral-state estimates upstream of the candidate branches under PIC-local, ML-BM, GLS-BM, and Rphy-BM, frequently moving them toward the corresponding Stable-process estimates and, where available, nearby fossil references. For example, at node 149, which lies upstream of both SBLC-*φ* and SBLC-*ω*, the ML/GLS/Rphy-BM estimate decreased from 11.38 m in the unpatched reconstruction to 9.66 m after joint *φ* + *ω* patching, toward the Stable-process estimate of 8.97 m and fossil reference FR14 of 7.41 m. At the evaluated lateral and descendant positions, PIC-local estimates were unchanged by patching, whereas the corresponding ML-BM, GLS-BM, and Rphy-BM estimates shifted from their unpatched values, consistent with the contrast between local and global reconstruction observed in the simulations.

The responses to individual and joint candidate-branch patching were spatially structured. SBLC-*φ* and SBLC-*ω* occurred near each other in the phylogeny. When only one branch was patched, the other remained connected to their shared upstream nodes and continued to influence the ancestral-state estimates there. Joint *φ* + *ω* patching therefore produced larger reductions in those estimates than patching either branch alone. By contrast, the more distant SBLC-*ψ* had little influence on estimates around *φ* and *ω*, and adding *ψ* to the joint *φ* + *ω* configuration produced virtually no further change in that region. Bayes-MCMC responses were less regular and showed no comparable directional or distance-dependent pattern.

Thus, the preceding simulation sensitivity analyses and the empirical candidate-branch analyses addressed complementary aspects of multiple SBLCs: the simulations showed that the effect of adding a second unpatched SBLC varied among methods, nodes, and branch pairs; in the empirical analysis, when only one candidate branch was patched, a nearby untreated candidate SBLC exerted greater residual influence on ancestral-state estimates than a phylogenetically distant one. Detailed node-wise results are provided in Supplementary Results Sections S2.2–S2.3 and Table S7.

## Discussion

Ancestral-state reconstruction depends on both phylogenetic signal and the assumed evolutionary process, and a model that describes average trait variation adequately can still be locally misspecified when evolution includes pulsed or heavy-tailed change (Litsios and Salamin 2012, Elliot and Mooers 2014, Gao and Wu 2022b). Our results show that such localized misspecification can produce a structured error footprint extending beyond the affected branch. On intact trees, the direction, magnitude, and phylogenetic extent of SBLC-induced error depended on both reconstruction method and node position. Patching substantially reduced these errors at nodes adjacent to and farther from the affected branch and lowered average error across the phylogeny. Tree partitioning can therefore limit an important consequence of localized evolutionary heterogeneity while retaining the selected ASR method and applying it to tree components expected to be more internally compatible with its background-model assumptions.

The spatial structure of the errors reflected how each reconstruction method combined information across the phylogeny. Under the specific recursive contrast-based local implementation considered here, ASR errors are propagated sequentially upstream, aligning with the operational bottom-up traversal conceptualized by Felsenstein (1985); SBLC-induced distortions were consequently concentrated along the ancestral path above the affected branch. The global methods instead use a phylogeny-wide covariance structure, allowing a localized change to influence upstream, lateral, and descendant-side estimates (Martins and Hansen 1997, Schluter et al. 1997, Huelsenbeck and Bollback 2001). These effects did not always decline monotonically with topological separation: some attenuated rapidly, whereas others persisted or changed direction, and the direction of ML-BM error at the descendant-side boundary differed between the 128-tip and 4,096-tip configurations. The effect of an SBLC therefore cannot be inferred from its magnitude or phylogenetic distance alone. Patching reduced the distortion by interrupting the transmission of information across the candidate branch while leaving the reconstruction procedure within each component unchanged.

We found no evidence that this local correction was obtained at the cost of net deterioration elsewhere. Tree-wide mean absolute error remained lower after patching even when the four focal nodes, or a broader set of 14 nearby nodes, were excluded. The principal corrective response also persisted across changes in tree size, descendant-subtree size, topology, and the distribution of evolutionary variance under λ-transformed BM and early-burst processes. When two SBLCs occurred together, the effect of adding the second unpatched change varied among methods, nodes, and branch pairs. Phylogenetic proximity sometimes strengthened the cumulative distortion but did not determine it by itself. Joint patching nevertheless reduced the principal errors in both nearby and distant configurations, showing that the procedure can be extended to more than one candidate boundary.

Because patching is conditional on designated cut positions, candidate selection is central to its application and interpretation. Candidate branches may be nominated by independent biological or fossil evidence or by process-based jump models (Landis et al. 2013); unusually large phylogenetically independent contrasts may provide a simpler preliminary clue for further examination (Chen et al. 2026). These sources provide graded rather than definitive evidence. Process-based methods assign posterior support to possible jump histories, whereas contrast-based screening depends on a chosen threshold for identifying observations that warrant further examination. Candidate branches should therefore be regarded as model- and criterion-dependent hypotheses rather than as verified historical events. Nevertheless, at the *k* = 1 baseline, partitioning a branch without an imposed SBLC produced little additional directional error. Under the conditions examined, this negative control suggests that patching is relatively robust to false-positive candidates. Branch selection should still be biologically or statistically motivated, with plausible alternatives compared when the assignment to a particular branch is uncertain.

In our simulations, residual errors after patching were generally similar in scale to those from Stable-process reconstruction, although this descriptive comparison should not be interpreted as a formal test of equivalence or superiority. The generally small Stable-process errors under the SBLC conditions are consistent with the capacity of heavy-tailed models to accommodate occasional large evolutionary changes. Stable-process errors were also generally close to zero at k = 1, indicating no appreciable directional-error penalty when no SBLC was imposed under the conditions examined. The MCMC implementation used here can be computationally demanding, especially on large phylogenies (Gao and Wu 2022b), but this is an implementation-specific consideration rather than an inherent limitation of heavy-tailed evolutionary models, and more computationally tractable approaches are available (Bastide and Didier 2023). Computational cost should therefore not be the primary basis for choosing between the two approaches.

Instead, the choice should be guided by the inferential question. Stable-process reconstruction and patching should be viewed as complementary approaches to non-gradual trait evolution rather than as competing methods intended to answer the same question.

Patching is conditional on a candidate SBLC-bearing branch and asks how strongly conventional ancestral-state estimates depend on information transmitted across that branch. By restricting this influence while retaining the original reconstruction method, it may be useful when independent biological evidence, fossil information, or a separate shift-detection analysis supports the inference of an SBLC on a particular branch, or when comparability with conventional ASR results is important. Stable, by contrast, provides a coherent process-based framework in which occasional changes of large magnitude arise from a heavy-tailed distribution, without requiring affected branches to be specified in advance (Elliot and Mooers 2014). When the primary objective is to infer the underlying evolutionary process without prespecifying candidate branches, Stable or another model of pulsed or heavy-tailed evolution may therefore be more appropriate. The two approaches can also be used together, with Stable providing a process-based reconstruction and patching evaluating the branch-specific robustness of conventional estimates.

The cetacean analysis illustrates patching primarily as an empirical sensitivity analysis rather than as a means of selecting a uniquely correct reconstruction. Movement of conventional estimates toward Stable-process estimates and nearby fossil references provides useful external agreement, but cannot validate either the reconstructed states or the inferred SBLC locations because the fossils are not observations of those ancestors. Differences between individual and joint patching further show that empirical conclusions may depend on which nearby candidate branches remain connected to the analyzed region. Empirical applications should therefore compare plausible patch configurations and emphasize conclusions that remain robust to uncertainty in candidate-branch assignment.

The OU analyses identify a general information limit. Under an OU process, the influence of a realized ancestral state on its descendants decays as trait values are drawn toward the optimum, with more rapid loss under stronger mean reversion (Hansen 1997, Hansen et al. 2008). Strong post-SBLC mean reversion therefore progressively removed information about ancestral states within the affected subtree from the descendant tip values. Patching continued to limit upstream and lateral distortion produced by the remaining anomalous signal, but none of the examined approaches—including patched conventional methods and Stable-process reconstruction—accurately recovered ancestral states within the SBLC-descendant subtree once information about those states had largely disappeared from descendant tip values. This represents a general information limit for all ASR methods that rely exclusively on terminal trait data: once subsequent evolution has largely erased the signal of a realized ancestral state, the observed tips provide little basis for recovering it accurately. Patching should therefore be understood as controlling the transmission of available information across a candidate boundary, not as restoring historical information that is no longer retained in the observed data.

Although simple to implement, patching provides a systematic way to test the robustness of ASR to localized model misspecification. Our analyses clarify when tree partitioning can reduce reconstruction error and when loss of historical information prevents recovery. More broadly, the general usefulness of a background model across most of a phylogeny does not guarantee robustness to an atypical change on a single branch. Patching does not determine whether such a change occurred. Instead, it shows how strongly a conventional reconstruction depends on information crossing a candidate branch. Used alongside explicit evolutionary models and independent biological or fossil evidence, patching provides a practical way to assess whether conclusions from conventional ASR remain robust after the influence of a candidate branch is restricted.

## Acknowledgments

We thank Zheng-Lin Chen for his earlier observation, made during his work on robust phylogenetic independent contrasts (Chen et al. 2026), that a single abrupt shift on a branch inflates not only the associated phylogenetically independent contrast but also the contrasts at upstream nodes. This observation helped motivate the present study. General-purpose large language models (ChatGPT 5, OpenAI; Gemini 3, Google; DeepSeek V3, DeepSeek; ChatGLM-4.7, Zhipu AI; Kimi K3, Moonshot AI) were each used for one or more of the following tasks: drafting and refining text; checking the consistency of numerical values in the text, tables, and figures; and proofreading mathematical expressions for consistency in symbols and typography. The authors reviewed and edited all model-assisted output and take full responsibility for the content of this article.

## Supplementary Materials

Supplementary material, including Supplementary Methods Sections S1–S5, Supplementary Results Sections S1–S2, Figures S1–S16, and Tables S1–S7, is available at SYSBIO online submission system.

## Data availability

The taxonomically matched empirical datasets and pruned phylogenies derived from previously published sources, the fossil-reference data, simulation inputs, machine-readable results underlying all figures and tables, complete reproduction code, computational-environment files, workflows, documentation, and Repository Appendix A are available in Dryad at https://doi.org/10.5061/dryad.47d7wm3wt; the original data sources are cited in Materials and Methods. The versioned patchASR software is archived in Zenodo at https://doi.org/10.5281/zenodo.23319830, with the development version available at https://github.com/Dreabder/patchASR.

