## Supplementary materials for "Ancestral State Reconstruction under Localized Evolutionary Change: Error Patterns and Their Control by Tree Partitioning"

for

### Supplementary Methods

#### Section S1. Implementation Details for Conventional ASR Methods and the Stable-Process Benchmark

This section describes the software implementations and method-specific settings used for the five conventional ASR methods and the stable-process benchmark summarized in the main text. Unless otherwise stated, functions were called with their default settings, and identical settings were used for the unpatched and patched analyses within each conventional method.

The first conventional ASR method was a local, contrast-based reconstruction implemented with ace(..., method = "pic") in ape (1, 2). This approach is conceptually related to Felsenstein's method of phylogenetically independent contrasts and estimates ancestral values through recursive calculations along neighboring branches and nodes. It was included as a representative local reconstruction procedure in which the influence of an SBLC may be transmitted stepwise through the tree.

The second method was maximum-likelihood reconstruction under a Brownian motion (BM) model, implemented with ace(..., method = "ML", model = "BM") in ape (2-4). This method estimates ancestral states under an explicit stochastic model fitted to the phylogeny and observed tip values, and was included as a standard likelihood-based ASR method for continuous traits.

The third method was generalized least-squares reconstruction under a BM phylogenetic covariance structure, implemented with corBrownian() and ace(..., method = "GLS") in ape (2-4). This method was included because it represents an explicitly covariance-based formulation of ancestral reconstruction, in which tip values are combined according to the expected covariance structure induced by the phylogeny.

The fourth method was a phylopars-type global reconstruction implemented with phylopars(...) in Rphylopars (5). Although Rphylopars was developed primarily for phylogenetic analyses of datasets with missing trait values and within-species variation, the same framework can be used to obtain estimates of unobserved internal-node values under a phylogenetic covariance model. In our implementation, we used a single-trait BM model with phenotypic error disabled and REML-based parameter estimation. This method was included because it provides a widely used framework for joint inference of trait evolution parameters and ancestral values while remaining within the broad class of conventional BM model-based ASR methods.

The fifth method was Bayesian ancestral reconstruction implemented with anc.Bayes(...) in phytools (6-8). Unlike the point-estimate methods, this approach estimates posterior distributions of ancestral states through MCMC sampling. For comparability with the other methods, posterior means were used as point estimates when calculating reconstruction errors and patch performance. For each simulation replicate, the MCMC was run for 10,000 generations, with samples saved every 100 generations. The first 20% of stored samples were discarded as burn-in, leaving 80 posterior samples for summarizing ancestral states at each internal node. Other anc.Bayes control settings were left at their defaults. The implementation checked that burn-in removal did not eliminate all stored samples and that posterior samples could be extracted for all internal nodes. No additional convergence diagnostics, such as effective sample sizes, trace plots, or multiple-chain comparisons, were implemented in this batch analysis.

Stable-process ancestral reconstruction, analyzed separately as the external benchmark, was implemented with StableTraitsR() in RRphylo (9, 10). This approach differs from the conventional

BM-based reconstructions above by modeling continuous trait evolution with a stable model, which allows heavy-tailed evolutionary changes and can therefore accommodate large branch-localized changes without assuming gradual BM increments. The RRphylo function StableTraitsR() provides an R interface to the external programs StableTraits and StableTraitsSum, rather than reimplementing the stable model entirely within R. In our implementation, the phylogenetic tree and observed tip trait values were supplied to StableTraitsR(), with four MCMC chains and 200,000 iterations per chain. For each internal node, the posterior median stored in ST_fit$aces$Median was used as the stable-process ancestral state estimate.

For clarity in figures and tables, we refer to the five conventional methods as PIC-local, ML-BM, GLS-BM, Rphy-BM, and Bayes-MCMC, respectively, and to stable-process reconstruction as Stable. The purpose of including multiple conventional methods was not to exhaust all available ASR algorithms, but to test whether SBLC-induced distortion and patch performance were consistent across different reconstruction procedures; Stable was evaluated separately as an external benchmark.

#### Section S2. Tree-Wide and Propagation Error Summaries

##### S2.1 Propagation analysis and node-class summarization

The propagation-coordinate labels identify topological position classes and do not necessarily correspond to unique nodes. A position class could therefore contain multiple nodes within a replicate; for example, the $K^1$ and $J^2$ classes each contained two nodes. For every reconstruction method and analysis condition, signed error was first calculated separately at each node as the reconstructed state minus the corresponding simulated true state.

Within each replicate, these errors were then averaged arithmetically across all nodes assigned to the same position class. The resulting replicate-level class averages, rather than the individual node-level errors, constituted the 100 independent observations used for subsequent summarization. For each position class, the plotted point was the mean of these 100 replicate-level class averages. A 95% confidence interval for the mean was obtained from 10,000 nonparametric bootstrap resamples of the 100 replicate-level class averages, using the 2.5th and 97.5th percentiles of the bootstrap distribution.

##### S2.2 Tree-wide evaluation of reconstruction error

To complement the focal-node analysis described in the main text, we evaluated whether patching altered reconstruction error across the internal nodes of the primary simulated phylogeny. This analysis used the same 100 independently generated trait realizations on the fixed 128-tip balanced tree. For each conventional ASR method and each SBLC scenario, patched and unpatched ancestral-state estimates were available at all 127 ancestral non-tip nodes, including the root. Stable-process reconstruction was not included in these paired comparisons because it was used only as an external benchmark and was not subjected to the patch procedure.

Errors at different nodes within the same trait realization are generally correlated because they share the same simulated evolutionary history and phylogeny. Nodes were therefore not treated as independent sampling units, and the 12,700 node-by-replicate observations were not pooled as independent data. Instead, the internal nodes defined the domain over which reconstruction error was summarized within each realization, whereas the 100 independently generated trait realizations constituted the sampling units used to estimate uncertainty. This use of realization-level summaries follows the general principle of cluster-level analysis, in which correlated lower-level observations are summarized within an independent higher-level unit and inference is based on the resulting higher-level summaries (11). The analogy is limited to the statistical structure; phylogenetic nodes and trait realizations are not literal participants or clusters.

Three nested internal-node sets were evaluated. The all-node set contained all 127 internal nodes, including the root. The focal-excluded set contained the 123 nodes remaining after exclusion of the four prespecified focal nodes shown in Fig. 1 of the main text: $H$, $I$, $J$, and $J^1$. For the broader local-exclusion analysis, we removed the two endpoints of the SBLC-bearing branch and every internal node within two branching steps of either endpoint. On the fixed 128-tip balanced phylogeny, this topology-based rule excluded 14 local nodes, including the four focal nodes, and retained 113 internal nodes for analysis.

For each trait realization, ASR method, and node set, absolute reconstruction error was calculated at every retained node as the absolute difference between the reconstructed and true ancestral states. These node-level absolute errors were then averaged within each realization to obtain separate mean absolute error (MAE) values for the patched and unpatched reconstructions. Because the two reconstructions were obtained from the same simulated trait realization, the effect of patching within realization $r$ was summarized as

$$\Delta\mathrm{MAE}_r = \mathrm{MAE}_{\mathrm{patched}} - \mathrm{MAE}_{\mathrm{unpatched}}.$$

The reported contrast, denoted $\overline{\Delta MAE}$, was the mean of the 100 realization-specific paired differences, $\Delta\mathrm{MAE}_r$. Negative values indicate lower average reconstruction error after patching, whereas positive values indicate higher error after patching.

The all-node contrast quantified the overall tree-wide change in reconstruction error. The two exclusion analyses addressed whether an apparent overall improvement was driven primarily by nodes near the SBLC-bearing branch and whether partitioning the tree caused collateral deterioration at more distant nodes by reducing the number of tips and the phylogenetic information available to each subtree-based reconstruction. Progressively excluding the focal and broader local node sets provided increasingly stringent assessments of reconstruction performance away from the affected branch.

For each combination of ASR method, SBLC scenario, and node set, uncertainty in $\overline{\Delta\mathrm{MAE}}$ was quantified by resampling the 100 independent trait realizations with replacement. For each bootstrap iteration, 100 realization indices were sampled with replacement. Whenever a realization was selected, its complete patched–unpatched pair and its complete vector of internal-node errors were retained; nodes were not resampled individually. The same sampled realization indices were used for all three node sets within each bootstrap iteration. This procedure was repeated 10,000 times, and the 2.5th and 97.5th percentiles of the resulting bootstrap distribution were used as the limits of the 95% confidence interval.

All MAE contrasts were calculated separately for each ASR method, SBLC direction, and simulation condition. A confidence interval entirely below zero was interpreted as evidence of lower expected MAE after patching, whereas an interval entirely above zero indicated higher expected MAE. Intervals overlapping zero provided no clear evidence of a mean difference.

#### Section S3. Sensitivity Analyses

##### Overview

The primary simulations evaluated how single-branch large changes (SBLCs) affect ancestral state reconstruction (ASR) and whether the patch procedure reduces the resulting reconstruction error. To assess robustness beyond the primary reference setting, we conducted two complementary sets of analyses: controlled one-factor-at-a-time sensitivity analyses on idealized phylogenies and extensions on an empirical mammalian phylogeny.

The controlled analyses were centered on a balanced ultrametric tree with 128 terminal taxa, unit root-to-tip height, an SBLC-descendant subtree ($T_1$) containing eight taxa, a BM background process, and upsizing and reciprocal downsizing SBLCs with $k$ = 16 and $k$ = 1/16, respectively. One major feature was varied at a time while the remaining settings were held at their reference values. These analyses varied SBLC magnitude, total phylogenetic size, $T_1$ size, tree topology, and the background evolutionary model. The magnitude series included $k$ = 1 as a no-SBLC baseline; unless otherwise stated, the remaining analyses were conducted under both $k$ = 16 and $k$ = 1/16.

Across all simulations, SBLCs were treated as operational branch-localized perturbations superimposed on the specified background evolutionary process. True ancestral states were retained as reference values for evaluating reconstruction error, and the conventional ASR methods and patch procedure described in the main text were applied consistently. The empirical-topology extensions examined contrasting local and surrounding tree shapes, post-SBLC evolutionary duration under strong OU attraction, and spatial interactions between nearby and phylogenetically distant pairs of SBLCs, including the performance of joint patching. These extensions tested the framework under realistic combinations of branch lengths and topology.

#### S3.1 General simulation framework

All sensitivity analyses followed the same general structure. First, a phylogenetic tree was specified or generated. Second, continuous trait values were simulated along the tree under a background evolutionary model. Third, one or more candidate SBLC-bearing branches were selected. Fourth, branch-localized perturbations were introduced on the selected branches. Fifth, conventional ASR methods were applied to the original full tree, and patched reconstructions were obtained by cutting the SBLC-bearing branch or branches and reconstructing ancestral states independently on the resulting tree components. Finally, estimated ancestral states were compared with the true ancestral values recorded during simulation.

For each simulation replicate, trait values at all terminal and internal nodes were retained. These recorded values represented the known evolutionary history for that replicate and were used as the reference values in the node-wise error analysis. Reconstruction error was evaluated primarily using signed error,

$$E_{r,n}^{method} = \hat{A}_{r,n}^{method} - A_{r,n}^{true},$$

where $r$ denotes the simulation replicate, $n$ denotes the internal node being evaluated, $\hat{A}_{r,n}^{method}$ is the ancestral state estimated by a given ASR method, and $A_{r,n}^{true}$ is the true ancestral state recorded during simulation for the same replicate and node. Positive values of $E_{r,n}^{method}$ indicate overestimation, whereas negative values indicate underestimation.

All comparisons between unpatched and patched reconstructions were paired within replicate, because both reconstructions were applied to the same simulated trait dataset. Node-wise errors were not pooled across internal nodes as if nodes were independent observations. Instead, errors were evaluated separately at predefined focal nodes or focal-node categories defined relative to the SBLC-bearing branch.

#### S3.2 SBLC magnitude

The primary analysis used multiplicative SBLCs with magnitudes $k$ = 16 and $k$ = 1/16. This setting was chosen as a strong localized perturbation that generated clearly detectable SBLC-induced distortion while retaining variation among reconstruction methods. To assess whether the observed patterns depended on this particular magnitude, we repeated the simulations across a gradient of SBLC strengths.

- For upsizing SBLCs, the multiplicative factor was varied as $k$ = 1, 2, 4, 8, 16, 32, and 64.
- For downsizing SBLCs, the reciprocal factors were used, $k$ = 1, 1/2, 1/4, 1/8, 1/16, 1/32, and 1/64.

The value $k = 1$ represents the no-SBLC baseline, under which the focal branch follows the background evolutionary process without an imposed branch-localized perturbation. Including $k = 1$ allowed us to verify whether patching induces an artificial effect when no large localized change is present.

For a candidate SBLC branch from upstream node $I$ to descendant node $J$, the background transition under BM would be

$$x_J^{BM} = x_I + \epsilon_{IJ},$$

where $x_I$ is the true state at node $I$, and $\epsilon_{IJ}$ is the branch-specific evolutionary increment. To introduce a multiplicative SBLC, the true value of node $I$ itself was retained, but the starting value for the transition into node $J$ was shifted to $kx_I$. The value at node $J$ was therefore generated as

$$x_J^{SBLC} = kx_I + \epsilon_{IJ}.$$

After this transition, all descendants of $J$ evolved under the same background evolutionary process, using $x_J^{SBLC}$ as the root value of the descendant subtree. This implementation localizes the imposed large change to the *I–J branch* while allowing subsequent evolution in the descendant subtree to proceed under the same model used elsewhere in the tree.

The downsizing simulations used the same procedure with $0 < k < 1$. These simulations were included to verify that the patch procedure responded to the direction of the SBLC-induced distortion rather than to a method-specific tendency toward overestimation. Under an upsizing SBLC, affected upstream or neighboring nodes were expected to show positive signed error if the localized large value influenced their reconstruction. Under a downsizing SBLC, the corresponding distortion was expected to be negative.

The full magnitude gradient was used to determine whether reconstruction error and patch performance changed smoothly with SBLC intensity or whether strong changes produced threshold-like behavior.

##### S3.3 Phylogenetic size

To determine whether patch performance depended on the number of terminal taxa, we repeated the simulations on balanced ultrametric trees of different sizes. In addition to the primary 128-tip tree, we considered trees with $N$ = 64, 128, 256, 512, and 1,024 terminal taxa.

To isolate the effect of total tree size, the local SBLC configuration was kept as close as possible to that of the primary analysis. In particular, the descendant subtree rooted at the SBLC downstream node $J$, denoted $T_1$, was kept at 8 terminal taxa. The primary SBLC magnitudes, $k = 16$ for upsizing shifts and $k = 1/16$ for downsizing shifts, were used, and the background evolutionary model was BM.

All trees were scaled to a total root-to-tip height of 1. This scaling ensured that the marginal evolutionary variance accumulated from the root to any terminal taxon was comparable across trees of different sizes under the same BM rate parameter. As in the primary analysis, the root trait value was set to $x_{\text{root}} = 5$, and the BM rate parameter was set to $\sigma^2 = 1$.

Because $T_1$ was fixed across tree sizes, the relative depth of the SBLC-bearing branch necessarily varied with total tree size in balanced trees. We therefore interpret this analysis as testing sensitivity to total phylogenetic background size under a fixed local SBLC subtree size, rather than as a test of SBLC depth. Sensitivity to SBLC depth and descendant-subtree size was evaluated separately.

This analysis was intended to test whether the qualitative behavior observed in the primary 128-tip tree remained stable when the same local SBLC configuration was embedded within progressively larger phylogenetic backgrounds.

##### S3.4 Descendant-subtree size and local phylogenetic context

The effectiveness of the patch procedure may depend on the amount of phylogenetic information separated by the SBLC-bearing branch. In particular, the size of the descendant subtree rooted at the downstream node $J$ determines the amount of trait information available for reconstructing the descendant side after the SBLC-bearing branch is removed. To evaluate whether patch performance depended on this local phylogenetic context, we varied the size of the descendant subtree ($T_1$) while keeping other simulation settings consistent with the primary analysis.

Candidate SBLC-bearing branches were selected to generate descendant subtrees ($T_1$) of different sizes, ranging from terminal-lineage events ($T_1 = 1$) to larger descendant clades. For each selected $T_1$ size, the total tree size, SBLC magnitude, and background evolutionary model were kept constant unless otherwise stated.

The descendant-subtree size was defined as the number of terminal taxa descended from node $J$, where the SBLC-bearing branch connects upstream node $I$ to downstream node $J$. Candidate SBLC-bearing branches were selected from the 128-species balanced tree used in the primary analysis to generate descendant subtrees of different sizes, with $T_1 = 1, 2, 4, 8$, and 16 terminal taxa. When $T_1 = 1$, the SBLC corresponds to a lineage-specific change occurring directly on a terminal branch. Although such cases do not contain internal nodes within the descendant subtree for downstream ancestral reconstruction, they represent an important limiting case of highly localized evolutionary change and were retained for evaluating upstream distortion and patch correction.

##### S3.5 Random ultrametric tree topology

The primary analysis used a fixed balanced tree, which provides a controlled and interpretable setting but does not capture the full variation in empirical tree shape. To evaluate whether the results were sensitive to topology, we repeated simulations on randomly generated ultrametric trees.

Random ultrametric trees with 128 terminal taxa were generated using a pure-birth (Yule) process implemented with the pbtree function in the R package phytools. Each tree was scaled to unit root-to-tip height before trait simulation. For each random tree, an SBLC-bearing branch was selected such that the descendant subtree rooted at $J$ contained 8 terminal taxa, matching the descendant-subtree size used in the primary balanced-tree analysis. The upstream side of the SBLC-bearing branch was also required to contain sufficient ancestral structure for node-wise evaluation, including at least three ancestral nodes along the path from the upstream boundary node $I$ toward the root.

The same BM background model and the same paired SBLC magnitudes were then applied to each random tree. Upsizing SBLCs used $k = 16$, and downsizing SBLCs used $k = 1/16$. Thus, this analysis isolated the effect of replacing the balanced tree with randomly generated ultrametric topologies, rather than simultaneously changing tree size, SBLC magnitude, descendant-subtree size, or the background evolutionary model.

Because node identities are not comparable across independently generated trees, focal nodes were defined using the SBLC-centered structural coordinate system. Under this system, nodes were classified by their topological position relative to the SBLC-bearing branch rather than by absolute labels. These categories included the immediate upstream boundary node $I$, upstream ancestral nodes along the path from $I$ toward the root, the descendant-side boundary node $J$, nodes within the descendant subtree rooted at $J$, and lateral or sister-lineage nodes outside the SBLC descendant subtree. For each random tree, focal-node categories were redefined relative to the locally selected SBLC-bearing branch. This ensured that, for example, the “upstream boundary node” always referred to the node immediately upstream of the SBLC-bearing branch, even though its absolute identity differed among random trees.

This random-tree analysis tested whether the main conclusions depended on the symmetry and regular branching structure of the balanced tree used in the primary simulation. It also allowed evaluation of SBLC-induced distortion and patch performance across more heterogeneous local branching patterns while preserving the primary simulation settings in all other respects.

#### S3.6 Alternative background evolutionary models

The primary simulations used Brownian motion (BM) as the background evolutionary process. To determine whether SBLC-induced ancestral-state distortion and the corrective performance of patching depended on this assumption, we repeated the analyses under three alternative classes of continuous-trait models: Pagel’s $\lambda$-transformed BM models, Ornstein–Uhlenbeck (OU) models, and early-burst (EB) models. The covariance parameterizations followed Huang et al. (12), in which λ-BM, OU, and EB were expressed as modifications of a common BM baseline. In the present study, these models were used as alternative background processes on which the same controlled SBLC perturbation and reconstruction workflow were superimposed.

Changing an evolutionary-model parameter can affect two analytically distinct properties of the simulated traits. First, it can change the *relative covariance structure*: which taxa share more or less evolutionary information, how rapidly phylogenetic memory decays, or whether evolutionary variance is concentrated on deeper versus more recent branches. Second, if the diffusion-rate parameter is held numerically constant, the same parameter change can also alter the *overall marginal variance scale* of the trait. These two effects must be distinguished because the reconstruction errors used in this study are expressed in the original trait units. If one model setting generates a background trait distribution with a larger standard deviation than another, its raw signed errors and their confidence intervals may also be larger even when the reconstruction method has the same performance relative to the background variation. A difference in raw error magnitude would then reflect both covariance structure and trait scale, rather than covariance structure alone.

Extremely weak-dependence settings that provide little biologically meaningful information for ASR were not considered. Specifically, we excluded $\lambda = 0$, which removes off-diagonal phylogenetic covariance by definition, and the weakest-dependence OU and EB settings, $h/H = 0.05$ and $\rho = 8$, identified by Huang et al. (12). The retained settings span substantial differences in the strength and temporal distribution of phylogenetic dependence while avoiding limiting cases in which internal-node reconstruction has little phylogenetic information to use.

##### S3.6.1 BM-referenced normalization of marginal tip variance

To facilitate comparisons among evolutionary models, we used a BM-referenced normalization of marginal tip variance, under which the parameters of each alternative model were calibrated so that the variance accumulated from the fixed root to any terminal taxon matched the corresponding BM reference value.

All trees in the controlled model-sensitivity analysis were ultrametric and scaled to a common root-to-tip height $H = 1$. The root state was fixed at $x_{\mathrm{root}} = 5$. Under the BM reference process, the diffusion-rate parameter was $\sigma^2_{\mathrm{BM}} = 1$, so the marginal background variance accumulated from the fixed root to any terminal taxon was

$$v_{\mathrm{ref}} = \sigma^2_{\mathrm{BM}} H = 1.$$

We used this root-to-tip marginal variance as the common variance scale for the $\lambda$-BM, OU, and EB comparisons. Let $\mathbf{V}_\psi$ denote the theoretical covariance matrix generated by a model with parameter setting $\psi$ before variance normalization, and let $v_\psi$ denote its terminal marginal variance. Because the trees were ultrametric, all diagonal entries of $\mathbf{V}_\psi$ were equal to $v_\psi$. The normalized covariance matrix was

$$\mathbf{V}^*_\psi = \frac{v_{\mathrm{ref}}}{v_\psi} \mathbf{V}_\psi.$$

This multiplication changes only the overall variance scale. It does not change the correlation matrix implied by a parameter setting:

$$\mathrm{Cor}\left(\mathbf{V}^*_\psi\right) = \mathrm{Cor}\left(\mathbf{V}_\psi\right).$$

Consequently, OU attraction can still alter the rate at which phylogenetic memory decays, and EB rate heterogeneity can still redistribute variance between deeper and more recent branches, but every parameter setting is expressed on the same terminal background-variance scale. Differences in raw signed errors among normalized settings can therefore be interpreted more directly in relation to differences in covariance structure and temporal variance allocation, rather than to a simple expansion or contraction of the entire trait scale.

Normalization was applied to the theoretical covariance or, equivalently, to the model diffusion rate or transformed branch lengths *before* trait simulation. It was not performed by standardizing each realized simulated dataset. Replicate-specific z-score standardization would condition the transformation on random sample variation, would remove biologically relevant differences among replicates, and could change the realized size of the imposed SBLC relative to the background process. It would therefore represent a different simulation design rather than a clean control of theoretical background variance.

After background traits had been simulated, the SBLC was imposed using the same operational procedure and multipliers as in the primary BM analysis. For a background model with parameter setting ψ, let $x_J^{\mathrm{bg},\psi}$ denote the state at node *J* that would have arisen without an imposed SBLC, and define the realized background change on the focal branch as

$$\delta_{IJ}^{(\psi)} = x_J^{\mathrm{bg},\psi} - x_I.$$

This quantity represents the complete realized background transition under the corresponding model. For example, under an OU model, it includes both the deterministic mean-reversion component and the stochastic residual. The perturbed state at node *J* was constructed as

$$x_J^{\mathrm{SBLC},\psi} = k x_I + \delta_{IJ}^{(\psi)} = x_J^{\mathrm{bg},\psi} + (k-1) x_I.$$

Thus, the additional displacement attributable to the imposed SBLC was

$$x_J^{\mathrm{SBLC},\psi} - x_J^{\mathrm{bg},\psi} = (k-1) x_I,$$

whereas the total realized change from node (I) to node (J) was

$$x_J^{\mathrm{SBLC},\psi} - x_I = (k-1)x_I + \delta_{IJ}^{(\psi)}.$$

When $k = 1$, the imposed displacement is zero and the unperturbed background transition is recovered exactly.

The primary upsizing and downsizing settings were $k = 16$ and $k = 1/16$, respectively, and these settings were used for all background models.

For each model parameter setting, the model-specific variance treatment was applied consistently throughout the data-generating process, including whole-tree background simulation, the realized background transition on the *I*–*J* branch, and resimulation of the descendant subtree below node *J*. The ancestral-state reconstruction method-specific settings were otherwise kept identical to those used in the primary BM analyses. True ancestral values were recorded after simulation, and signed and absolute errors were calculated in the common original trait units. Because $v_{\mathrm{ref}} = 1$, no additional rescaling of the reported errors was required.

##### S3.6.2 Pagel's λ-transformed Brownian models

Pagel's λ-transformed BM models were used to vary the strength of shared phylogenetic covariance while retaining the same tree topology and marginal variance scale. Starting from the BM covariance matrix $\mathbf{V}_{\mathrm{BM}}$, λ rescales the off-diagonal entries but leaves the diagonal entries unchanged:

$$V_{ij}^{(\lambda)} = \begin{cases} V_{ii}^{\mathrm{BM}}, & i = j, \\ \lambda V_{ij}^{\mathrm{BM}}, & i \neq j. \end{cases}$$

Thus, $\lambda = 1$ recovers the standard BM covariance structure, whereas smaller positive values reduce covariance among lineages without changing the marginal variance of any terminal taxon. We considered $\lambda$ = 0.25, 0.50, 0.75, and 1.00.

Because Pagel's λ leaves the diagonal of the covariance matrix unchanged, these settings were already matched to the BM marginal variance and required no additional variance normalization. This analysis therefore provides the most direct comparison of changing off-diagonal phylogenetic covariance at a fixed trait scale. It tested whether the propagation of a localized SBLC into upstream or neighboring ancestral-state estimates depends on the amount of shared covariance available to transmit its influence across the tree.

##### S3.6.3 Ornstein-Uhlenbeck models

OU models were used to represent constrained evolution toward a fixed adaptive optimum (13, 14). Along each branch, trait evolution followed

$$dX_t = \alpha(\theta - X_t)dt + \sigma dB_t,$$

where $\theta$ is the optimum, $\alpha > 0$ is the attraction parameter, $\sigma^2$ is the diffusion variance parameter, and $B_t$ is standard BM. The root state and optimum were fixed at $X_0 = \theta = x_{\mathrm{root}} = 5$.

The OU process was parameterized using the phylogenetic half-life ratio $h/H$, where $h$ is the time required for the expected deviation from the optimum to decay to one-half of its initial magnitude and $H$ is the total root-to-tip tree height. Because

$$\mathrm{E}(X_t - \theta \mid X_0) = (X_0 - \theta)e^{-\alpha t},$$

when $t = h$,

$$e^{-\alpha h} = e^{-\alpha t} = \tfrac{1}{2}.$$

Therefore, the attraction parameter was

$$h = \frac{\ln 2}{\alpha}, \quad \alpha H = \frac{\ln 2}{h/H}.$$

We considered $h/H$ = 0.10, 0.25, 0.50, and 1.00. Smaller $h/H$ values correspond to stronger attraction toward the optimum and faster decay of ancestral influence, whereas larger values correspond to weaker attraction and more persistent phylogenetic dependence. Because $h/H$ determines the dimensionless quantity $\alpha H$, these settings specify the extent to which ancestral deviations are expected to persist over a complete root-to-tip interval.

In addition to controlling the decay of the expected ancestral influence, $\alpha$ also determines how strongly stochastic changes arising during evolution are retained. Under the fixed-root implementation used here, the root state contributes no random variance. All variation among simulated trait histories therefore arises from stochastic perturbations accumulated along branches after the root. A perturbation arising early on a branch is progressively attenuated by attraction toward the optimum before reaching the terminal taxon, whereas a perturbation arising near the tip undergoes less attenuation. Integrating the contributions of these continuously arising and differentially attenuated perturbations over a root-to-tip interval of length $H$ gives the terminal marginal variance

$$v_{\mathrm{OU}}(H \mid \alpha, \sigma^2) = \frac{\sigma^2}{2\alpha}\left(1 - e^{-2\alpha H}\right).$$

Here, $\sigma^2$ controls the rate at which stochastic variation is introduced, whereas $\alpha$ controls how rapidly the effects of earlier perturbations are lost. Consequently, terminal variance increases with $\sigma^2$ but is increasingly constrained as attraction becomes stronger.

Covariance between two terminal taxa arises only from stochastic perturbations that occurred during their shared evolutionary history. Taxa $i$ and $j$ experience the same stochastic trajectory from the root to their most recent common ancestor (MRCA), but after divergence the stochastic changes occurring along their separate descendant branches are independent. The variance accumulated along the shared root-to-MRCA path therefore provides the initial shared component, which is then attenuated separately as it propagates from the MRCA to each terminal taxon. Accordingly, the covariance between terminal taxa $i$ and $j$ is

$$\mathrm{Cov}\left(X_i, X_j \mid X_0\right) = \frac{\sigma^2}{2\alpha}\left(1 - e^{-2\alpha t_{ij}}\right)e^{-\alpha\left(t_i + t_j - 2t_{ij}\right)},$$

where $t_i$ and $t_j$ are the root-to-tip distances and $t_{ij}$ is the length of the shared root-to-MRCA path. The term

$$\frac{\sigma^2}{2\alpha}\left(1 - e^{-2\alpha t_{ij}}\right)$$

represents the stochastic variance accumulated along the shared ancestral path, whereas

$$e^{-\alpha\left(t_i + t_j - 2t_{ij}\right)}$$

represents the combined attenuation of that shared component along the two post-MRCA lineages. For the ultrametric trees used here, $t_i = t_j = H$, so $t_i - t_{ij}$ and $t_j - t_{ij}$ are the respective MRCA-to-tip distances. Thus, covariance is greater when taxa share a longer ancestral path, when their MRCA is closer to the tips, or when attraction toward the optimum is weaker.

The parameter $\alpha$ therefore determines both the persistence of stochastic variation within individual lineages and the relative decay of shared phylogenetic covariance among lineages. By contrast, $\sigma^2$ multiplies all variances and covariances by the same factor and consequently controls the overall scale of the covariance matrix.

If $\sigma^2$ were held at 1 for every OU setting, changing $h/H$ would change both covariance decay and marginal variance. If no variance correction were applied, then for $H = 1$ and $\sigma^2 = 1$, the unnormalized terminal variances would be approximately 0.072, 0.180, 0.338, and 0.541 for $h/H = 0.10, 0.25, 0.50,$ and 1.00, respectively, compared with 1 under the BM reference. Thus, stronger attraction would generate both weaker long-range dependence and a narrower overall trait distribution. Raw error confidence intervals could become narrower partly because the simulated trait itself varies less, not necessarily because reconstruction or patching is intrinsically more precise.

To separate covariance decay from this overall scale effect, we selected an OU diffusion rate for each value of $\alpha$ such that the terminal marginal variance equaled $v_{\text{ref}}$:

$$\sigma^2_{\text{OU}}(\alpha) = v_{\text{ref}} \frac{2\alpha}{1 - e^{-2\alpha H}}.$$

With $H = 1$ and $v_{\text{ref}} = 1$, the required diffusion rates ($\sigma^2$) are approximately 13.863, 5.567, 2.957, and 1.848 for $h/H = 0.10, 0.25, 0.50,$ and 1.00, respectively. These rate adjustments do not reverse or weaken the intended differences in phylogenetic memory; they simply place all OU settings on the same terminal variance scale.

These formulas apply to the fixed-root OU implementation used in this study. If a simulation routine instead draws the root from the stationary OU distribution, the diagonal variance and the required scaling factor differ. The implementation was therefore checked to ensure that the root was fixed at $x_{\text{root}} = 5$ and that stationary-root variance was not added implicitly.

The same branch-localized SBLC was then superimposed on each variance-normalized OU background using the construction defined in Section S3.6.1. Specifically, for an OU parameter setting ψ,

$$x_J^{\text{SBLC},\psi} = x_J^{\text{bg},\psi} + (k-1)x_I,$$

where $x_J^{\text{bg},\psi}$ is the state that would have arisen at node $J$ under the OU background process without an imposed SBLC. Thus, the SBLC added $(k-1)x_I$ to the realized OU background state without replacing the model-specific background transition on the $I$–$J$ branch. After this modification, evolution within the descendant subtree proceeded from the modified state at node $J$ under the same OU process as elsewhere in the tree. This analysis tested whether patching remained effective across faster versus slower decay of phylogenetic memory after removing the otherwise confounding change in overall terminal trait variance.

###### S3.6.4 Early-burst models

EB branch-length transformations were used to represent temporal heterogeneity in the rate of trait evolution. Under the parameterization used here, negative values of the rate parameter correspond to the conventional early-burst pattern, in which evolutionary rates decline toward the present, whereas positive values represent temporal acceleration toward recent and terminal branches. Let $t$ denote elapsed time from the root, with the root at $t = 0$ and all terminal taxa at $t = H$, where $H$ is the height of the ultrametric tree. Time was first expressed as the normalized root-to-tip position

$$\mu = \frac{t}{H},$$

so that $\mu = 0$ at the root and $\mu = 1$ at the tips. The instantaneous relative evolutionary rate at normalized time $\mu$ was then defined as

$$r(\mu) = e^{\rho\mu},$$

where $\rho$ controls the direction and strength of temporal rate heterogeneity. We considered $\rho$ = −4, −2, 0, and 4.

The setting $\rho = 0$ recovers the BM baseline because $r(\mu) = 1$ throughout the tree. When $\rho < 0$, the relative rate is highest near the root and decreases toward the present. Consequently, a larger proportion of evolutionary variance is accumulated on deeper branches. Because evolutionary changes occurring on deeper branches are inherited by broader groups of descendants, these changes contribute more strongly to shared components of trait variation. When $\rho > 0$, the relative rate increases toward the present, shifting a larger proportion of evolutionary variance toward recent and terminal branches. This increases lineage-specific variation and reduces the relative contribution of deeply shared evolutionary history.

Following the EB time-transformation formulation described by Huang et al. (12), the transformed length of a branch extending from parent time $t_p$ to child time $t_c$ was obtained by integrating the instantaneous relative rate over that branch,

$$L_\rho(t_p, t_c) = \int_{t_p}^{t_c} r\left(\frac{t}{H}\right) dt$$

$$= \int_{t_p}^{t_c} e^{\rho t/H} dt$$

$$= \begin{cases} \dfrac{H}{\rho}\left[e^{\frac{\rho t_c}{H}} - e^{\frac{\rho t_p}{H}}\right], & \rho \neq 0, \\ t_c - t_p, & \rho = 0. \end{cases}$$

Thus, $L_\rho(t_p, t_c)$ represents the amount of evolutionary time accumulated on the branch after accounting for the temporally varying relative rate. For a diffusion coefficient $\sigma^2$, the corresponding contribution of this branch to evolutionary variance is

$$\sigma^2 L_\rho(t_p, t_c).$$

The expression for $\rho = 0$ is also the continuous limit of the first expression as $\rho \to 0$. Therefore, when $\rho = 0$, the transformed branch length reduces exactly to the original branch duration $t_c - t_p$.

More generally, the cumulative transformed length from the root to any time $t$ can be written as

$$B_\rho(t) = L_\rho(0, t) = \begin{cases} \dfrac{H}{\rho}\left(e^{\rho t/H} - 1\right), & \rho \neq 0, \\ t, & \rho = 0. \end{cases}$$

Because the tree is ultrametric, every terminal taxon occurs at $t = H$. The total transformed length from the root to any terminal taxon is therefore

$$B_\rho(H) = \begin{cases} \dfrac{H}{\rho}(e^\rho - 1), & \rho \neq 0, \\ H, & \rho = 0. \end{cases}$$

If the same diffusion rate were used for all EB settings, the terminal marginal variance conditional on the root state would be

$$\mathrm{Var}\left(X_{tip} \mid X_0\right) = \sigma_{BM}^2 B_\rho(H)$$

Under such an uncorrected parameterization, $B_\rho(H)$ would determine not only where evolutionary variance accumulates along the tree but also how much total variance accumulates between the root and the tips. For $H$ = 1 and $\sigma_{BM}^2$ = 1, the root-to-tip variance factors before variance normalization would be approximately 0.245, 0.432, 1.000, and 13.400 for $\rho = -4, -2, 0$, and 4, respectively. For example, without variance correction, the terminal marginal standard deviation under $\rho = 4$ would be $\sqrt{13.4} = 3.66$ times the BM standard deviation.

These uncorrected values are presented here only to demonstrate why variance normalization is required. They do not represent an additional set of non-normalized sensitivity analyses, and no unnormalized EB results were retained in the reported analyses. All EB simulations analyzed below were normalized so that differences among $\rho$ settings reflected the temporal placement of evolutionary variance rather than differences in the total root-to-tip variance.

To achieve this normalization, the transformed branch lengths were rescaled so that the total transformed length along every complete root-to-tip path equaled the original tree height $H$. Define the setting-specific normalization factor as

$$c_\rho = \frac{H}{B_\rho(H)}$$

Each transformed branch length was then rescaled according to

$$L_\rho^*(t_p, t_c) = c_\rho L_\rho(t_p, t_c) = L_\rho(t_p, t_c)\frac{H}{B_\rho(H)}.$$

Because the unscaled transformed branch lengths along a complete root-to-tip path sum to $B_\rho(H)$, the normalized transformed lengths along any complete root-to-tip path consequently satisfy

$$\sum_{\text{root-to-tip branches}} L_\rho^*(t_p, t_c) = c_\rho \sum_{\text{root-to-tip branches}} L_\rho(t_p, t_c)$$

$$= c_\rho B_\rho(H)$$

$$= \frac{H}{B_\rho(H)} B_\rho(H)$$

$$= H.$$

Thus, the normalization restores the total transformed root-to-tip length to the original tree height while preserving the relative allocation of transformed length among branches. When the common BM diffusion rate $\sigma_{BM}^2$ is applied to these normalized branch lengths, the terminal marginal variance becomes

$$\mathrm{Var}(X_{tip} \mid X_0) = \sigma_{BM}^2 \sum_{\text{root-to-tip branches}} L_\rho^*(t_p, t_c)$$

$$= \sigma_{BM}^2 H,$$

which is the same terminal marginal variance as under the BM baseline for every value of $\rho$.

The same variance normalization could alternatively be achieved by adjusting the EB diffusion coefficient while retaining the transformed branch lengths on their original scale. For clarity, however, the normalization used here was expressed entirely through the branch-length rescaling defined above, with the common BM diffusion coefficient retained across all EB settings. No additional diffusion-rate correction was applied.

For the four EB settings examined here, the setting-specific branch-length normalization factors defined above were approximately

$$c_\rho = \begin{cases} 4.075, & \rho = -4 \\ 2.313, & \rho = -2 \\ 1.000, & \rho = 0 \\ 0.0746, & \rho = 4 \end{cases}.$$

Accordingly, the normalized length of every EB-transformed branch was calculated directly as

$$L_\rho^*(t_p, t_c) = L_\rho(t_p, t_c) \times \begin{cases} 4.075, & \rho = -4 \\ 2.313, & \rho = -2 \\ 1.000, & \rho = 0 \\ 0.0746, & \rho = 4 \end{cases}$$

Thus, for $\rho = -4$ and $\rho = -2$, every EB-transformed branch length was multiplied by approximately 4.075 and 2.313, respectively. These factors compensate for the fact that the total transformed root-to-tip lengths under these settings are shorter than the original tree height. For $\rho = 4$, every transformed branch length was multiplied by approximately 0.0746, compensating for the much larger transformed root-to-tip length generated by this setting. No rescaling was required for $\rho = 0$, because this setting recovers the BM branch lengths exactly.

Because the same setting-specific factor $c_\rho$ was applied uniformly to every transformed branch within a given EB setting, the normalization did not alter the relative allocation of evolutionary variance among branches. For any two branches *a* and *b*,

$$\frac{L_\rho^*(a)}{L_\rho^*(b)} = \frac{c_\rho L_\rho(a)}{c_\rho L_\rho(b)} = \frac{L_\rho(a)}{L_\rho(b)}.$$

Thus, the normalization changed the overall root-to-tip variance scale without removing the temporal rate heterogeneity represented by $\rho$. Negative values of $\rho$ continued to allocate a greater proportion of the matched total evolutionary variance to deeper branches, whereas positive values continued to allocate a greater proportion to recent and terminal branches. The normalization removed only the differences in total root-to-tip variance that would otherwise accompany these alternative temporal patterns.

More generally, let $t_{ij}$ denote the elapsed time from the root to the most recent common ancestor of terminal taxa *i* and *j*. Their normalized shared transformed length is

$$c_\rho B_\rho(t_{ij}),$$

and their covariance conditional on the root state is therefore

$$\mathrm{Cov}(X_i, X_j \mid X_0) = \sigma_{BM}^2 c_\rho B_\rho(t_{ij}).$$

When $i = j$, $t_{ij} = H$, and this expression reduces to the common terminal marginal variance $\sigma_{BM}^2 H$ derived above. Accordingly, all EB settings shared the same terminal marginal variance as the BM baseline, while their covariance structures continued to differ according to the temporal placement of evolutionary variance.

The same SBLC protocol was subsequently applied to each normalized EB background. The performance of patched ASR methods could therefore be compared across deeper-branch-weighted and terminal-branch-weighted patterns of evolutionary variance accumulation while holding the terminal marginal variance constant across parameter settings. Differences in reconstruction error or patch performance among EB settings could consequently be attributed more directly to the temporal distribution of evolutionary variance rather than to differences in the overall scale of background trait variation.

##### S3.7 Empirical-topology sensitivity analyses

To evaluate whether conclusions obtained from idealized simulations remained robust under a biologically realistic phylogenetic structure, we extended the sensitivity analyses to an empirical mammalian phylogeny. The empirical tree was treated as a fixed structural framework independent of the simulated trait values, thereby preserving realistic heterogeneity in branching patterns and relative branch lengths. Unless otherwise stated, empirical-topology simulations used the baseline BM process with an upsizing SBLC $k = 16$ and the corresponding downsizing SBLC $k = 1/16$. Conventional and patched ASR were evaluated at the predefined SBLC-centered focal nodes using the same signed-error framework as in the primary simulations.

###### S3.7.1 Empirical mammalian phylogeny and tree processing

Mammalian phylogenetic relationships were obtained from the VertLife DNA-only node-dated phylogeny of Upham et al. (15). We used the DNA-only maximum-clade-credibility tree distributed in the associated Dryad repository. The downloaded NEXUS file contained 4,100 terminal taxa; after removal of the non-mammalian outgroup, 4,099 mammalian species were retained. Although Upham et al. (15) reported 4,098 DNA-sampled mammalian species, all 4,099 mammalian labels present in the downloaded tree that could be matched to the accompanying completed taxonomy were retained for the present analyses.

The downloaded tree showed only negligible deviations from ultrametricity attributable to numerical rounding in the Newick representation. To remove these discrepancies, shorter terminal branches were extended to the maximum root-to-tip distance, without altering topology or internal branch lengths. The largest adjustment was $8.65 \times 10^{-5}$ Ma. The processed tree was fully bifurcating, contained 4,099 terminal taxa and 4,098 internal nodes, and had a root-to-tip height of 179.21 Ma. For the simulation analyses described below, branch lengths were subsequently rescaled to a root-to-tip height of 1 while preserving their relative values and the empirical topology.

###### S3.7.2 Balanced and imbalanced local topological configurations

To evaluate whether local tree shape influenced SBLC-induced reconstruction error and patch performance, we screened internal branches on the empirical mammalian phylogeny for contrasting topological configurations at two spatial scales. For each candidate SBLC-bearing branch $I \rightarrow J$, the complete descendant subtree rooted at $J$ was defined as $T_1$. Candidate branches were restricted to those for which $T_1$ contained 6–10 terminal taxa and for which the ancestral chain $G \rightarrow H \rightarrow I \rightarrow J$ could be identified unambiguously.

Topological balance within $T_1$ was quantified using the normalized Colless index (16, 17). For a rooted binary subtree with $n$ tips, the raw Colless index was calculated as

$$C = \sum_v \left| n_{L,v} - n_{R,v} \right|,$$

where $n_{L,v}$ and $n_{R,v}$ are the numbers of descendant tips in the two daughter clades of internal node $v$. This value was normalized by its theoretical maximum,

$$C_{max}(n) = \frac{(n-1)(n-2)}{2}$$

to obtain $C_{T_1,\text{norm}}$. Candidate $T_1$ subtrees with $C_{T_1,\text{norm}} \leq 0.25$ were classified as balanced-like, whereas those with $C_{T_1,\text{norm}} \geq 0.75$ were classified as ladder-like; intermediate cases were excluded from the strict topology classes.

To characterize the broader phylogenetic environment independently of the internal topology of $T_1$, we defined a surrounding clade rooted at *G* and calculated a weighted $T_1$-collapsed Colless index. In this calculation, $T_1$ was treated as an indivisible terminal unit while retaining its actual descendant-tip count ($N_{T_1}$) as its weight. Thus, branching within $T_1$ did not contribute to the surrounding-topology metric, whereas the size of $T_1$ continued to contribute to imbalance in the surrounding nodes. If the *G*-rooted clade contained *n* tips and $T_1$ contained *m* tips, the weighted surrounding Colless index was normalized by

$$C_{max,\text{weighted}} = \frac{(n-m)(n+m-3)}{2}.$$

Surrounding clades with $C_{\text{sur,norm}} \leq 0.25$ were classified as balanced-like and those with $C_{\text{sur,norm}} \geq 0.75$ as ladder-like; intermediate cases were excluded.

Combining the two scales produced four focal configurations: ($\alpha$) balanced-like surrounding topology with a balanced-like $T_1$; ($\beta$) balanced-like surrounding topology with a ladder-like $T_1$; ($\gamma$) ladder-like surrounding topology with a balanced-like $T_1$; and ($\delta$) ladder-like surrounding topology with a ladder-like $T_1$. Representative placements were selected to maximize the expression of the defining topology of each class while keeping $T_1$ as comparable as possible among configurations. Within each class, candidate placements were ranked by their Euclidean distance in the two-dimensional normalized-Colless space $\left(C_{\text{sur,norm}}, C_{T_1,\text{norm}}\right)$ from the corresponding class-specific ideal point: $\alpha = (0,0), \beta = (0,1), \gamma = (1,0),$ and $\delta = (1,1)$. The two dimensions were weighted equally, and smaller distances indicated closer agreement with the idealized topology of the corresponding class. When multiple $T_1 = 8$ candidates were available within a class, the highest-ranked $T_1 = 8$ placement was selected; otherwise, the highest-ranked candidate across the allowed $T_1 = 6{\sim}10$ range was retained.

##### S3.7.3 Post-SBLC evolutionary duration under OU evolution

The empirical phylogeny allowed us to examine whether the recoverability of the immediate post-SBLC state depended on the duration of subsequent evolution under OU mean reversion. We selected three internal SBLC placements representing short, intermediate, and long post-SBLC evolutionary durations, quantified by the age of node *J*, which is equivalent to the *J*-to-tip distance in the ultrametric tree.

To minimize differences other than post-SBLC duration, candidate placements were restricted to $T_1$ = 8. All internal *J* nodes subtending exactly eight extant tips were enumerated, yielding 113 eligible candidates, and all possible three-placement combinations were evaluated. To ensure adequate temporal separation among the three placements, each triplet was assigned an age-separation score,

$$S_{\text{age}} = \frac{A_{\text{span}} + 2\min(g_1, g_2)}{2R_{\text{all}}},$$

where $A_{\text{span}}$ is the age difference between the youngest and oldest placements in the triplet, $g_1$ and $g_2$ are the two adjacent age differences, and $R_{\text{all}}$ is the age difference between the youngest and oldest *J* nodes among all eligible candidates. Higher $S_{\text{age}}$ values indicate greater overall temporal separation while also favoring an intermediate placement that is well separated from both extremes. Triplets attaining at least 90% of the maximum observed $S_{\text{age}}$ were retained for the subsequent matching step.

Among these temporally well-separated triplets, the final combination was selected by minimizing

$$S_{\text{mismatch}} = \Delta C_{\text{norm}} + \log\left(\frac{L_{IJ,max}}{L_{IJ,min}}\right),$$

where $\Delta C_{\mathrm{norm}}$ is the difference between the maximum and minimum normalized Colless indices among the three $T_1$ subtrees, and $L_{IJ,max}$ and $L_{IJ,min}$ are the longest and shortest $I \rightarrow J$ branch lengths among the three placements. Thus, adequate separation in post-SBLC duration was enforced first, after which differences in $T_1$ topology and SBLC-bearing branch length were minimized. The realized placements and their corresponding $J$-ages, $I \rightarrow J$ branch lengths, and Colless indices are reported in Supplementary Results Section S1.6.2.

For this analysis, traits were simulated under strong OU attraction $h/H = 0.1$ with both an upsizing SBLC ($k = 16$) and the corresponding downsizing SBLC ($k = 1/16$). Reconstruction error was evaluated primarily at the descendant-side boundary node $J$ and at successive downstream nodes within $T_1$, where OU mean reversion was expected to progressively erode the SBLC-induced signal. Upstream and lateral nodes were retained as secondary reference positions to assess whether weakening of the descendant signal also reduced SBLC-induced distortion outside $T_1$. This analysis tested whether the loss of recoverability observed at $J$ and along its downstream lineage under strong OU attraction on idealized trees also occurred on the empirical topology and became more pronounced with increasing post-SBLC evolutionary duration.

##### S3.7.4 Spatial interaction between paired SBLCs

To determine whether the effects of two co-occurring SBLCs depended on their phylogenetic proximity, and whether joint patching could reduce the reconstruction distortion associated with both changes, we compared nearby and distant paired-SBLC configurations on the empirical mammalian phylogeny. Candidate SBLC placements were restricted to branches with a complete $G$–$H$–$I$–$J$ ancestral chain, a descendant subtree $T_1$ containing 6–10 terminal taxa, and a balanced-like $T_1$ topology, following the definitions in Section S3.7.2.

Three candidate placements, denoted $\alpha$, $\beta$, and $\gamma$, were selected to construct one nearby and two distant pairs while controlling local subtree size and topology. To identify the nearby pair, we considered pairs of candidate placements with identical $T_1$ tip counts and identical normalized Colless indices. Phylogenetic proximity was quantified by the length of the shared root-to-MRCA path of their $J$ nodes, and the pair sharing the longest such path was designated $\alpha + \beta$. A third placement, $\gamma$, was then selected from candidates having the same $T_1$ tip count and normalized Colless index as $\alpha$ and $\beta$. Among these candidates, the placement with the shortest root-to-MRCA path shared with the $\alpha$–$\beta$ region was selected. The nearby configuration therefore comprised $\alpha + \beta$, whereas $\alpha + \gamma$ and $\beta + \gamma$ constituted the two distant configurations. Exact ties at either selection step were resolved deterministically according to branch order in the input tree.

The three selected placements were used to define six distinct simulation scenarios. In three single-SBLC scenarios, an SBLC was introduced separately at $\alpha$, $\beta$, or $\gamma$. In three paired-SBLC scenarios, two SBLCs were introduced within the same simulated trait realization at $\alpha + \beta$, $\alpha + \gamma$, or $\beta + \gamma$. Thus, each single placement and each pair was evaluated using its own set of simulated trait realizations; no simulation scenario contained SBLCs at all three candidate positions. The single-SBLC scenarios provided the corresponding reference distributions against which the effects of adding a second SBLC were evaluated.

For each single-SBLC scenario, two reconstruction strategies were compared: conventional ASR on the intact phylogeny and patched ASR performed separately on the two subtrees obtained by partitioning the phylogeny at the SBLC-bearing branch. For each paired-SBLC scenario, conventional ASR on the intact phylogeny was compared with joint patching, in which the phylogeny was partitioned at both SBLC-bearing branches and ASR was performed separately on the resulting three subtrees. Comparisons between the paired- and corresponding single-SBLC scenarios were used to assess whether the presence of a second SBLC increased reconstruction distortion and whether this effect depended on phylogenetic proximity. Within each paired-SBLC scenario, the comparison between conventional and jointly patched reconstructions evaluated whether joint patching reduced distortion associated with the two co-occurring changes.

#### Section S4. Empirical Cetacean Data, Candidate SBLC Identification, and ASR Analysis

##### S4.1 Cetacean body-length data and extant-species phylogeny

We used total body length (TL) as the continuous trait for the empirical cetacean analysis. Body-length records for extant and fossil cetaceans were compiled primarily from Burin et al. (18) and the associated Back to the Water dataset (19). When multiple TL records were available for the same terminal taxon, we used their arithmetic mean. All analyses were conducted using mean total length expressed in meters.

To avoid introducing additional uncertainty through trait imputation, we retained only taxa with directly available TL measurements (has_tl = TRUE and nonmissing tl_m). Values inferred through phylogenetic or morphological imputation in the source dataset were not used. Taxon names were standardized to the corresponding phylogenetic tip labels, which were used as the unique identifiers linking trait and phylogenetic data. After filtering and matching to the cetacean SAFE tree described below, 258 terminals had directly available body-length data, comprising 86 extant cetacean species and 172 fossil terminals. Because some fossil records represent incompletely identified or specimen-level operational taxonomic units rather than formally recognized species, we refer to these collectively as fossil terminals.

An independently derived molecular phylogeny was used for ancestral-state reconstruction and Lévy-process analysis of the extant taxa. We used the time-calibrated Cetartiodactyla phylogeny of Zurano et al. (20), which was inferred from mitochondrial sequence data using a Bayesian framework. After removal of the three outgroup taxa, the source tree contained 318 Cetartiodactyla terminals. The tree was pruned to the intersection between its cetacean taxa and the 86 extant species with directly measured TL data, yielding 79 extant cetacean species with unique matches between phylogeny and trait data. Pruning removed only unmatched terminals; the topology and branch lengths connecting the retained species were otherwise unchanged. This 79-tip time-calibrated molecular tree was used for all empirical ASR and Lévy-process analyses.

##### S4.2 Fossil phylogenetic information and mapping to the extant molecular tree

Fossil phylogenetic placement was obtained independently from the total-group cetacean metatree of Lloyd and Slater (21). Their framework synthesized phylogenetic information from previously published morphological matrices and molecular information for extant cetaceans and subsequently generated time-scaled trees incorporating both living and extinct taxa. We used the authors' taxonomically conservative SAFE reconstruction and its representative median time tree (Cetacea_Safe_Median.tre). The tree contained 494 extant and extinct terminal taxa in the downloaded dataset used here.

The SAFE tree was used only to provide fossil topology and temporal information and was not used for empirical ASR of the extant trait data. We first retained the 86 extant species and 172 fossil terminals having directly available TL data, producing a 258-tip trait-matched auxiliary tree. We then restricted its extant component to the same 79 species represented in the molecular analysis tree while retaining all 172 fossil terminals, yielding a 251-tip fossil-reference tree. Thus, fossil body lengths were never entered as terminal observations in the 79-species ASR analyses.

To transfer fossil placement from the SAFE tree to the 79-tip molecular tree, we used a topology-based anchoring procedure based exclusively on shared extant taxa. For each fossil terminal, we traversed the path from the fossil toward the root of the SAFE tree. The first nonempty side lineage encountered that contained one or more of the 79 extant reference species was designated the proximal anchor (A), and the next such side lineage encountered toward the root was designated the external anchor (B). When anchor A contained two or more extant species, their most recent common ancestor (MRCA) on the 79-tip molecular tree was identified using ape::getMRCA, providing a candidate molecular-tree branch for the fossil.

We retained only fossil mappings for which the local topology was unambiguous across the two phylogenies. Specifically, the extant taxa in anchor A were required to correspond to the complete descendant set of the candidate MRCA on the molecular tree, anchor B had to occur entirely on the sister side of that clade, and the proximal relationship had to be identifiable from the fossil's immediate extant sister lineage in the SAFE tree. These criteria were intended to retain only fossil placements that could be transferred between the total-group and molecular trees without requiring an arbitrary resolution of topological disagreement.

##### S4.3 Temporal consistency and fossil proxy nodes

Topologically accepted fossils were subsequently screened for temporal consistency with their candidate branches on the 79-tip molecular tree. Node ages and fossil ages were expressed in millions of years before present (Ma). For each fossil, we used the arithmetic midpoint of its reported age range as a representative age for branch localization. This midpoint was used only as an operational temporal coordinate and was not interpreted as a precise estimate of the fossil's geological age.

A fossil was retained only when its representative age fell within the temporal interval bounded by the parent and child nodes of its mapped molecular-tree branch. Fossils falling outside this interval were considered temporally incompatible with the proposed mapping and were excluded from node-specific comparisons.

For each temporally compatible fossil, we calculated its absolute temporal distance to the parent and child nodes bounding the branch and assigned the fossil to the nearer endpoint. This node was designated the fossil-associated node for subsequent ASR comparisons. Cases in which the fossil was exactly equidistant from the two endpoints were excluded because the available temporal information did not provide a unique node assignment. After topological screening, temporal consistency filtering, and unique proxy-node assignment, 15 fossil terminals were retained as external paleontological reference points.

Importantly, these fossils were not treated as observations of the ancestral states themselves. Fossil TL values were used only as independent paleontological references for evaluating the biological plausibility of reconstructions near the corresponding internal nodes. They therefore contributed neither to estimation of evolutionary-model parameters nor to ASR on the 79-species molecular tree.

##### S4.4 Lévy-process identification of candidate branch-localized changes

Candidate SBLC-bearing branches were identified using the Lévy-process framework of Landis et al. (22). This approach was selected because its evolutionary formulation closely matches the phenomenon considered here: continuous background evolution combined with discrete evolutionary jumps localized to individual branches. It therefore provides a more direct operational correspondence to an SBLC than methods designed primarily to identify shifts in evolutionary rate, long-term mean, or OU optimum, whose phenotypic consequences may extend across multiple descendant branches.

The analysis used the 79-tip time-calibrated molecular tree and extant-species TL values in meters. We fitted the jump-normal (JN) Lévy model (modelType = 3) while holding the phylogenetic topology and branch lengths fixed (fixBranches = True). Four independent Markov chain Monte Carlo (MCMC) chains were run for 2,000,000 cycles each. The first 100,000 cycles of each chain were discarded as burn-in, parameter and branch-specific jump states were sampled every 1,000 cycles, and standard run information was written every 10,000 cycles. The jump proposal scale was set to sigmaJumpProposal = 1.0, and the BM tuning parameter to tuningBM = 0.9995.

After burn-in, posterior samples from the four chains were aligned by sampling cycle and combined. For each branch, we summarized the posterior distribution of jump magnitude by its mean, median, standard deviation, and 2.5%, 5%, 95%, and 97.5% quantiles, together with the posterior probabilities of positive and negative jump magnitudes.

Branches were initially ranked using a branch-length-normalized signal-to-noise statistic calculated from the posterior distribution of jump magnitude. For branch $b$, let $\bar{J_b}$ and $s_{J,b}$ denote the posterior mean and posterior standard deviation of jump magnitude, respectively, and let $\mathrm{l}_b$ denote branch length. Following Landis et al. (22), the statistic was calculated as

$$\mathrm{SNR_b} = \frac{\bar{J_b}/s_{J,b}}{\mathrm{l}_b} = \frac{\bar{J_b}}{s_{J,b}\mathrm{l}_b}.$$

We used the absolute value of this statistic, $|\mathrm{SNR_b}|$, only as a ranking criterion for comparing candidate branches, rather than as a hypothesis-test statistic or a predefined threshold for evidence of a jump. The sign of $\mathrm{SNR_b}$ retained the inferred direction of the change.

MCMC performance was evaluated for model parameters and the principal candidate branches using rank-normalized split-$\hat{R}$, bulk effective sample size (ESS), tail ESS, and the Monte Carlo standard error (MCSE) of the posterior mean. We used $\hat{R} \leq 1.01$, bulk ESS $\geq$ 400, tail ESS $\geq$ 400, and MCSE/posterior SD $\leq$ 0.05 as quality-control criteria. We additionally verified that all four chains completed successfully, that parameter and jump-state samples were aligned after burn-in, that the topology and branch lengths represented in the posterior output matched the input 79-tip tree, and that branch-specific SNR values could be reproduced independently from the posterior samples.

##### S4.5 Selection and interpretation of candidate SBLCs

After MCMC quality control, all branches were ranked by decreasing |SNR|, and the five highest-ranking branches were examined for posterior support for jump direction. For a branch with a positive posterior mean jump, the relevant one-sided posterior tail probability was $P(J \leq 0)$; for a branch with a negative posterior mean, it was $P(J \geq 0)$. A candidate was retained when this opposite-direction posterior probability was < 0.05. Operationally, this was equivalent to requiring the central 90% posterior credible interval for jump magnitude to lie entirely on one side of zero: the 5th posterior percentile had to exceed zero for a positive jump, or the 95th percentile had to be below zero for a negative jump.

Branches satisfying this criterion were retained as candidate SBLCs. The retained branches were ranked by decreasing |SNR| and labeled alphabetically in that order. They were subsequently used as candidate cut points in the empirical patch analyses, both individually and in combination. The number, direction, estimated magnitudes, and phylogenetic positions of the retained candidates are reported in Supplementary Results Section S2.1.

We emphasize that these candidate branches were not interpreted as independently verified historical SBLCs. Localization of an unobserved phenotypic change from terminal observations necessarily depends on the assumed phylogeny, evolutionary model, and inferred latent evolutionary history. The Lévy analysis was therefore used as a reproducible procedure for identifying biologically plausible candidate branches rather than as a source of empirical ground truth. Candidate branches were also examined in their local phylogenetic and phenotypic context; patterns more consistent with progressive, directionally concordant change across successive branches were not regarded as clear examples of a single branch-localized event.

This distinction is particularly important for evaluating the patch procedure. The empirical analysis asks whether reconstruction changes in a biologically interpretable manner when branches with strong evidence for localized trait change are treated as cut points; it does not require the jump detector to identify the true historical location of every large evolutionary change. The $k = 1$ simulation, in which patching was applied in the absence of an imposed SBLC, showed that an unnecessary cut introduced little additional reconstruction error under the conditions examined (Fig. 6 of the main text).

##### S4.6 Empirical ancestral-state reconstruction and fossil comparison

The unpatched, patched, and stable-process analyses were all performed on the same 79-tip molecular tree and the same extant TL dataset. Conventional ASR was first performed on the intact tree. For patched reconstructions, the candidate SBLC branches identified above were used as cut points, and ancestral states were reconstructed on the resulting tree components using the same method-specific procedures. Candidate branches were examined both individually and, where applicable, jointly in the multiple-SBLC analysis. Stable-process reconstruction was performed separately as an external model-based benchmark.

Finally, the 15 fossil reference points were linked to the corresponding fossil-associated nodes for comparison with unpatched, patched, and stable-process ancestral estimates. Fossil body lengths were treated as external paleontological references rather than as known ancestral-state values.

#### Section S5. Software Implementation and Reproducibility

The patch procedure was implemented in the open-source R package patchASR (version 0.1.0; https://doi.org/10.5281/zenodo.23319830). The package accepts a phylogeny, continuous-trait data, and one or more user-specified candidate branches; partitions the phylogeny at those branches; maintains a mapping between component-tree and original-tree node identities; applies a selected ASR implementation independently to the resulting tree components; and merges the estimates into the node system of the original phylogeny. The archived release provides interfaces to the five conventional ASR implementations evaluated in this study. The partitioning layer does not modify the reconstruction algorithm or its model assumptions, parameter settings, priors, or MCMC configuration. Consequently, the patched and corresponding unpatched analyses differ only in whether the ASR implementation is applied to the intact phylogeny or independently to the partitioned components.

Software behavior was evaluated at several levels. Unit tests cover input validation, tree partitioning, preservation of node identities, boundary-node handling, result merging, invariance to the ordering of multiple candidate branches, and integration with the supported ASR implementations. Automated numerical regression tests compare package outputs with version-controlled reference outputs generated using the original research implementation and, where applicable, direct calls to the underlying ASR software. The provenance of each reference output and the method-appropriate numerical acceptance tolerances are documented with the test data. Seed-controlled tests are used for stochastic calculations. The archived release passed automated R CMD checks under R development, release, and previous-release versions on Linux and under the release version of R on macOS and Windows.

Study-specific code and materials are maintained separately from the general-purpose patchASR package. The accompanying reproducibility archive (https://doi.org/10.5061/dryad.47d7wm3wt) contains the scripts, required input and derived data, random-number seeds, software-version information, machine-readable numerical outputs, and instructions needed to reproduce the simulations, empirical analyses, tables, and figures reported in the article and Supplementary Materials. It also contains the complete outputs summarized in Repository Appendix A. The archive provides an ordered workflow, identifies the scripts and source files associated with each reported output, and notes analyses with substantial computational requirements. The archived package release and reproducibility materials correspond to the versions used for this study; subsequent package development is available at [final GitHub URL].

### Supplementary Results

#### Section S1. Sensitivity Analysis of Patch Performance

##### S1.1 Sensitivity to the magnitude of downsizing SBLCs

Reciprocal downsizing SBLCs broadly reversed the directional distortions observed under upsizing SBLCs (Fig. S3). Before patching, PIC-local produced increasingly negative errors at the upstream nodes $I$ and $H$ as the multiplier decreased below $k = 1$, while errors at the lateral node $J^1$ and the descendant-side boundary node $J$ remained comparatively small. ML-BM and GLS/Rphy-BM showed more broadly distributed underestimation across $I$, $H$, and $J^1$, together with overestimation at $J$. Bayes-MCMC showed relatively modest negative errors at $I$, $H$, and $J^1$ but increasingly pronounced overestimation at $J$. The absolute errors were smaller than those produced by the corresponding upsizing SBLCs, reflecting the inherent asymmetry of the multiplicative perturbation: downsizing approaches a lower bound as $k$ approaches zero, whereas upsizing is not similarly bounded. After patching, signed errors remained close to zero across the examined multipliers, methods, and focal nodes and were generally similar in scale to those obtained with the Stable-process benchmark.

The upsizing and reciprocal downsizing magnitude gradients showed broadly reversed error directions and a comparable corrective response to patching. Accordingly, the remaining sensitivity results are presented below for $k = 16$ as a common strong-perturbation reference. All corresponding analyses were also conducted under $k = 1/16$. Complete graphical and tabulated results for these reciprocal analyses are provided in Repository Appendix A of the archived reproducibility materials (https://doi.org/10.5061/dryad.47d7wm3wt).

##### S1.2 Sensitivity to phylogenetic tree size

Varying total phylogenetic size from 64 to 1,024 terminal taxa altered the magnitude of reconstruction error in some method–node combinations but did not change the principal spatial patterns produced by an upsizing SBLC of $k = 16$ (Fig. S4). Before patching, PIC-local consistently showed pronounced overestimation at the upstream nodes $I$ and $H$, while errors at $J^1$ and $J$ remained comparatively small. ML-BM showed greater variation in error magnitude among tree sizes, including nonmonotonic responses at several focal nodes, but retained the broader pattern of overestimation at $I$, $H$, and $J^1$ together with underestimation at $J$. GLS/Rphy-BM likewise showed persistent overestimation at $I$, $H$, and $J^1$ and pronounced underestimation at $J$. For Bayes-MCMC, increasing the number of terminal taxa clearly attenuated overestimation at $I$, $H$, and $J^1$, whereas the strong underestimation at $J$ persisted rather than declining with tree size. Thus, the effect of increasing phylogenetic size was both method- and node-dependent: it reduced some components of SBLC-induced distortion, particularly the more broadly distributed errors of Bayes-MCMC, but did not consistently eliminate the principal biases in the unpatched reconstructions. After patching, signed errors were substantially reduced and remained close to zero across the examined tree sizes, with no systematic deterioration as the number of terminal taxa increased. Their residual scale remained broadly comparable to that of the Stable-process benchmark. Patch performance therefore remained qualitatively consistent beyond the 128-tip tree used in the primary simulation.

##### S1.3 Sensitivity to descendant-subtree size and local phylogenetic context

Varying the size of the SBLC-affected descendant subtree from 1 to 16 terminal taxa altered the magnitude of several reconstruction errors but did not change the principal spatial pattern produced by an upsizing SBLC of $k = 16$ (Fig. S5). Before patching, PIC-local consistently showed pronounced positive errors at the upstream nodes $I$ and $H$, whereas errors at the lateral node $J^1$ and the descendant-side boundary node *J* remained comparatively small. These errors changed little across descendant-subtree sizes. The global methods retained their characteristic combination of positive displacement at *I*, *H*, and $J^1$ and negative displacement at *J*, although their responses to subtree size were method- and node-dependent. ML-BM showed moderate nonmonotonic variation, whereas the corresponding changes for GLS/Rphy-BM were generally more gradual. Bayes-MCMC also showed changes in error magnitude across subtree sizes. Thus, changing the amount of descendant information and the local phylogenetic context of the SBLC-bearing branch affected the magnitude, but not the basic direction or spatial distribution, of unpatched reconstruction error.

After patching, signed errors remained close to zero across the examined descendant-subtree sizes, methods, and focal nodes, with no systematic deterioration as $T_1$ increased. Residual errors were also generally comparable in scale to those obtained with the Stable-process benchmark. Patch performance therefore remained qualitatively consistent across the examined range of descendant-subtree sizes, from terminal-lineage events to larger affected clades, and did not depend strongly on the eight-tip descendant subtree used in the primary analysis.

##### S1.4 Sensitivity to random ultrametric tree topology

Replacing the fixed balanced tree with randomly generated Yule trees did not qualitatively alter the principal patterns of SBLC-induced reconstruction error (Fig. S6). Under the upsizing SBLC ($k = 16$), unpatched PIC-local reconstructions retained pronounced positive errors at the upstream nodes *I* and *H*, whereas errors at $J^1$ and *J* remained comparatively small. The global methods—ML-BM, GLS/Rphy-BM, and Bayes-MCMC—continued to show broader positive displacement at *I*, *H*, and $J^1$ together with pronounced negative errors at the descendant-side boundary node *J*. The largest topology-dependent changes were observed for ML-BM. Relative to the balanced tree, random topologies reduced its positive errors at most upstream and lateral focal nodes but substantially increased the negative error at *J*. Thus, tree topology affected the magnitude and spatial distribution of ML-BM error while preserving its characteristic combination of positive displacement at *I*, *H*, and $J^1$ and negative displacement at *J*. Other topology-dependent differences were generally smaller, and occasional changes in error direction were largely confined to cases in which the errors themselves were already small.

Patching substantially reduced signed errors under both balanced and random topologies. Under the random Yule topologies, all bootstrap confidence intervals for patched reconstructions included zero. The few confidence intervals excluding zero occurred only on the balanced tree and represented small residual deviations relative to the corresponding unpatched errors. Stable-process reconstructions likewise remained close to zero under both topology conditions. Overall, patch performance therefore remained qualitatively robust to variation in ultrametric tree topology.

##### S1.5 Sensitivity to alternative background evolutionary models

The principal pattern of SBLC-induced reconstruction error was broadly retained when the BM background process was replaced by variance-normalized Pagel's $\lambda$-BM, OU, or EB models, although the strength of the response differed among model classes (Figs. S7–S9). The $\lambda$-BM and EB gradients produced relatively modest changes in error, whereas OU evolution generated a pronounced and mechanistically distinct response because mean reversion progressively reduced the terminal signature of the SBLC.

Across the **$\lambda$-BM gradient**, unpatched reconstructions retained the directional and spatial patterns observed under standard BM (Fig. S7). Under the upsizing SBLC, PIC-local consistently produced positive errors along the upstream path from *I* and *H*, while the global methods showed more broadly distributed positive errors outside the descendant subtree and negative errors at $J$. Changing $\lambda$ from 0.25 to 1.00 altered the magnitude of some method–node combinations, particularly for the global methods, but did not produce a systematic change in the overall distortion pattern. After patching, signed errors were close to zero across $\lambda$ settings, methods, and focal nodes and were generally comparable in scale to those of the stable-process benchmark.

Results under the **EB backgrounds** were similarly consistent across the examined rate gradients (Fig. S8). Whether evolutionary variance was concentrated relatively early in the tree ($\rho < 0$), distributed according to BM ($\rho = 0$), or shifted toward more recent branches ($\rho > 0$), the principal directions and spatial distributions of unpatched error remained recognizable. PIC-local continued to show distortion primarily along the upstream path, whereas ML-BM, GLS/Rphy-BM, and Bayes-MCMC showed broader effects across upstream and lateral-lineage nodes together with opposing error at $J$. Most changes across $\rho$ were small relative to the underlying SBLC-induced errors. Patching reduced errors toward zero throughout the EB gradient, with no evidence of systematic deterioration under either early- or late-concentrated evolutionary variance.

**OU evolution** produced a substantially stronger model gradient than either $\lambda$-BM or EB evolution (Fig. S9). This pattern can be understood most directly by first considering the descendant-side boundary node $J$. After the SBLC displaced the state at $J$ away from the OU optimum, mean reversion within the descendant subtree progressively attenuated this displacement as it propagated toward the terminal taxa. As attraction toward the optimum became stronger—that is, as $h/H$ decreased—the terminal states therefore retained progressively less information about the immediate post-SBLC state at $J$. At $h/H = 0.10$, the expected proportion of the initial displacement retained at the terminal depth of $T_1$ was only approximately 5.1%. This loss of signal is illustrated directly in Figure S10: despite comparable immediate post-SBLC states at $J$ under BM and strong OU evolution, the post-SBLC deviation remained evident throughout $T_1$ under BM but was rapidly attenuated toward the optimum under OU. Consequently, $J$ became increasingly difficult to reconstruct, producing pronounced underestimation under the upsizing scenario.

The same loss of phylogenetic memory had the opposite consequence for nodes outside the affected descendant subtree. Because the terminal observations carried a weaker signature of the SBLC under stronger OU attraction, this localized change exerted less influence on the reconstruction of upstream and lateral-lineage nodes, and their unpatched errors generally decreased as $h/H$ declined. Conversely, as $h/H$ increased and the SBLC-induced displacement persisted more strongly to the terminal taxa, $J$ became more accurately recoverable, but the stronger anomalous descendant signal produced greater distortion at upstream and lateral-lineage nodes, progressively approaching the pattern observed under BM. ML-BM exhibited an isolated nonmonotonic response at $h/H = 0.25$ under the upsizing SBLC, similar to its method-specific irregularity at one setting in the descendant-subtree analysis. Because this deviation was not shared by the other methods and disappeared after patching, it did not alter the general model-sensitivity pattern.

Patching effectively reduced OU-associated errors at upstream and lateral-lineage nodes but provided little improvement at $J$ under strong OU attraction. Patched Bayes-MCMC and Stable-process reconstruction showed modestly smaller errors at $J$ at the weaker OU settings ($h/H = 0.25$, 0.50, and 1.00), but this improvement disappeared under the strongest attraction ($h/H = 0.10$), where substantial residual error persisted across all approaches. This shared failure follows directly from the purpose and information requirements of the procedure: patching can prevent the anomalous descendant signal from distorting reconstruction outside the affected subtree, but it cannot recover the immediate post-SBLC state at $J$ once its signature has been largely erased from the terminal data by mean reversion during subsequent evolution. Figure S10 illustrates that this information loss arises during the evolutionary process itself, before any reconstruction method is applied. Thus, the residual error at $J$ under strong OU attraction reflected a general lack of recoverable descendant information—and hence a fundamental identifiability constraint for any ancestral-state reconstruction based solely on terminal-node data—rather than a limitation specific to the reconstruction approaches examined here.

Overall, patching remained effective across $\lambda$-BM, EB, and OU backgrounds in preventing the localized change from distorting reconstruction of the upstream and lateral portions of the phylogeny. However, the OU analysis identified an important boundary of the procedure: patching can remove downstream contamination of upstream reconstruction, but it cannot recover a transient post-SBLC ancestral state when strong mean reversion during subsequent evolution has erased its signal.

#### S1.6 Empirical-topology sensitivity analyses

##### S1.6.1 Balanced and imbalanced local topological configurations

Screening identified 92, 49, 3, and 2 candidate placements in classes $\alpha$–$\delta$, respectively. These four classes represented the four combinations of surrounding topology and $T_1$ topology: class $\alpha$, balanced-like surrounding topology with a balanced-like $T_1$; class $\beta$, balanced-like surrounding topology with a ladder-like $T_1$; class $\gamma$, ladder-like surrounding topology with a balanced-like $T_1$; and class δ, ladder-like surrounding topology with a ladder-like $T_1$. Representative placements were selected to emphasize the defining topology contrast while keeping $T_1$ as similar as possible across classes. The selected $\alpha$ and $\beta$ placements each had $T_1 = 8$ and strongly balanced-like surrounding topologies, whereas the selected $\gamma$ and $\delta$ placements each had $T_1 = 6$ and the most strongly ladder-like surrounding topologies available within their respective classes (Fig. S11a–d).

Across the four empirical topology classes, the principal method-specific patterns of SBLC-induced reconstruction error remained broadly consistent with those observed in the idealized-tree analyses, although error magnitude varied among placements (Fig. S12). Under the upsizing SBLC ($k = 16$), PIC-local continued to show its strongest distortion at the upstream nodes, whereas the global methods generally showed broader displacement outside $T_1$ together with opposing error at $J$. Among the methods, ML-BM was the most sensitive to the realized empirical topology, showing marked class-to-class variation and, in some focal-node comparisons, changes in error direction. Thus, local topological configuration affected the magnitude and, occasionally, the local direction of unpatched error without altering the overall SBLC-centered distortion pattern.

Patching substantially reduced these topology-dependent differences. Patched errors remained close to zero across the four classes and were broadly comparable in scale to the Stable-process benchmark, with no systematic deterioration from balanced-like to ladder-like configurations. Patch performance therefore remained robust across strongly contrasting local topologies on the empirical mammalian phylogeny.

##### S1.6.2 Post-SBLC evolutionary duration under OU evolution

The selected short-, intermediate-, and long-duration placements had $J$-ages of 1.92, 22.80, and 42.21 Ma, respectively. All three SBLC placements had $T_1 = 8$, while their $I \rightarrow J$ branch lengths were 4.84, 6.31, and 5.03 Ma and their normalized $T_1$ Colless indices were 0.286, 0.286, and 0.238, respectively. Thus, the placements spanned a large difference in the duration of post-SBLC evolution while retaining broadly similar SBLC-bearing branch lengths and descendant-subtree topologies (Fig. S13). Under OU evolution, deviations from the trait optimum decay progressively through time, so a longer interval between $J$ and the terminal taxa provides more opportunity for the SBLC-induced signal to be attenuated by mean reversion.

At the descendant-side boundary node *J*, the empirical-topology results reproduced the information-loss pattern observed in the idealized OU simulations (Fig. 7 of the main text and Table S6). As post-SBLC duration increased, OU mean reversion progressively weakened the descendant signal of the localized change, increasing reconstruction error at *J* across both unpatched and patched analyses. Under the upsizing SBLC ($k$ = 16), this shared duration effect was evident most directly in PIC-local, whose estimate at *J* is essentially unaffected by patching: mean signed error changed from approximately -5 in the short-duration placement to -43 and -60 in the intermediate- and long-duration placements, respectively. The global unpatched methods were additionally affected by intact-tree reconstruction across the SBLC boundary, so their errors could depart further from this underlying OU-driven error scale. These results indicate that increasing post-SBLC OU duration progressively reduced the information available in the terminal observations about the immediate post-SBLC state at *J*, irrespective of whether patching was applied.

Patching therefore did not eliminate the OU-induced error at *J*; rather, for the global methods, it largely removed the additional distortion associated with reconstructing across the intact SBLC-bearing branch. After patching, ML-BM, GLS/Rphy-BM, and Bayes-MCMC generally converged toward the PIC-local and Stable-process error scale. Under the upsizing SBLC, patched errors at *J* were approximately -60, -43, and -5 for the long-, intermediate-, and short-duration placements, closely matching Stable-process errors of -60.49, -43.43, and -5.30. Bayes-MCMC provided a particularly clear example of the distinction: under upsizing, its unpatched error remained near -74 across durations, whereas patching reduced this additional intact-tree distortion and revealed the underlying duration-dependent error scale of approximately -60, -43, and -5. Thus, substantial residual error after patching reflects loss of recoverable descendant information rather than a failure specific to the patch procedure.

The downstream internal nodes within $T_1$ broadly supported the same interpretation (Table S6). Shorter post-SBLC duration generally produced smaller residual errors after patching, and patched global reconstructions were usually similar in scale to PIC-local and Stable-process estimates. The duration effect became less uniform farther downstream, however, with the relative ordering of the long- and intermediate-duration placements varying among nodes because of their nonidentical empirical branch configurations. Thus, the signal-loss pattern evident at *J* persisted within $T_1$, but its expression became more heterogeneous with downstream position.

Outside $T_1$, the effect of post-SBLC duration was reversed (Table S6). When the duration was short, the descendant tips retained a stronger SBLC signal, and this signal produced greater distortion in some unpatched upstream reconstructions. This pattern was especially pronounced for PIC-local and GLS/Rphy-BM. For example, under the upsizing SBLC, PIC-local error at *I* increased from 9.18 in the long-duration placement to 14.47 in the intermediate placement and 38.56 in the short-duration placement, while GLS/Rphy-BM increased from 6.72 to 10.75 and 20.00. The response was more method-dependent for ML-BM and Bayes-MCMC, and lateral-node effects were generally smaller and less regular than those at the upstream nodes. After patching, upstream errors at *H* and *I* remained small across post-SBLC durations.

##### S1.6.3 Spatial interaction and joint patching of co-occurring SBLCs

The single- and paired-SBLC scenarios produced broadly similar location- and method-dependent patterns of reconstruction error (Figs. S14–S15). In these figures, patched results represent partitioning at the single affected branch in the single-SBLC scenarios and joint partitioning at both affected branches in the paired-SBLC scenarios. The nearby pair $\alpha + \beta$ and the two distant pairs $\alpha + \gamma$ and $\beta + \gamma$ were evaluated alongside the corresponding single-SBLC scenarios at $\alpha$, $\beta$, and $\gamma$. Adding a second SBLC did not produce a uniform increase in reconstruction error. Differences between the single- and paired-SBLC scenarios were generally smaller and less consistent than the effects of focal placement, SBLC direction, and reconstruction method.

At the upstream nodes *H* and *I*, unpatched PIC-local, ML-BM, and GLS/Rphy-BM reconstructions generally showed positive signed errors under the upsizing SBLC (Fig. S14). Thus, the direction of upstream bias largely followed the direction of the imposed change. Its magnitude was strongly dependent on SBLC placement: errors associated with $\alpha$ and $\beta$ were generally larger than those associated with $\gamma$, and these placement-specific patterns were largely retained when the focal SBLC occurred as part of a pair. The nearby $\alpha + \beta$ configuration produced larger unpatched errors than the corresponding single-SBLC scenarios in some node–method combinations, but this increase was not consistent across methods or between *H* and *I*. The distant configurations generally resembled their corresponding single-SBLC results. Across these methods, patching reduced errors at *H* and *I* toward the stable-process benchmark. Bayes-MCMC showed comparatively weak or variable upstream bias, and its patched and unpatched errors differed little. Joint patching reproduced the same method-specific pattern across all three paired configurations.

At the descendant-side boundary node *J*, the direction and magnitude of unpatched error were more strongly method dependent (Fig. S15). PIC-local estimates remained comparatively close to the stable-process benchmark and changed little after patching. Under upsizing, the global reconstruction methods showed negative signed errors at *J*, with the most pronounced underestimation observed for Bayes-MCMC. Joint patching moved the ML-BM, GLS/Rphy-BM, and Bayes-MCMC estimates substantially toward their corresponding stable-process benchmarks across all three paired configurations. This correction was especially pronounced for Bayes-MCMC, for which the large unpatched errors at *J* were almost completely removed after partitioning.

Results at $J^1$, shown for $\gamma$ alone and for the $\alpha + \gamma$ and $\beta + \gamma$ configurations, likewise provided no clear evidence that adding $\alpha$ or $\beta$ caused systematic additional distortion at this adjacent lateral node. Under upsizing, unpatched bias was most apparent for GLS/Rphy-BM. Across all ASR methods, the single- and paired-SBLC results were broadly similar, and patching generally moved the reconstructed values toward the stable-process benchmark.

Overall, the unpatched paired-SBLC scenarios retained the strong method- and location-dependent distortions observed in the corresponding single-SBLC scenarios. The presence of a second SBLC increased error in some configurations, although the magnitude of this additional distortion was not uniform and was not consistently predicted by phylogenetic proximity. More importantly, the corrective performance of joint patching was not compromised by the presence of two co-occurring SBLCs. At the upstream nodes *H* and *I*, joint patching reduced the pronounced errors of PIC-local, ML-BM, and GLS/Rphy-BM toward the stable-process benchmark, whereas the comparatively small or variable Bayes-MCMC responses remained largely unchanged and showed no evidence of deterioration after joint partitioning. At *J*, joint patching largely removed the pronounced errors of the global methods, while PIC-local remained close to the stable-process benchmark with or without patching. Patched errors at the adjacent lateral node $J^1$ were likewise generally comparable to the benchmark. Thus, the principal result was not that multiple SBLCs invariably produced additive error in unpatched ASR, but that simultaneously partitioning the phylogeny at both affected branches effectively controlled their reconstruction effects under both upsizing and downsizing conditions and for both nearby and distant configurations.

#### Section S2. Empirical Cetacean Results

##### S2.1 Identification of candidate sblcs

Among the five branches with the highest absolute signal-to-noise ratios, three satisfied the prespecified posterior-support criterion and were retained as candidate SBLCs (Fig. S16). In decreasing order of |SNR|, these branches were designated SBLC-$\varphi$, SBLC-$\psi$, and SBLC-$\omega$. All three had positive posterior mean jump magnitudes, corresponding to inferred increases in body length: 11.03 m, 4.01 m, and 7.13 m, respectively (|SNR| = 3.81, 2.63, and 1.91, in the same order). Because the SNR statistic incorporates posterior uncertainty and branch length, this ordering need not follow the ordering of the jump magnitudes themselves. SBLC-$\varphi$ and SBLC-$\psi$ occurred on terminal branches, whereas SBLC-$\omega$ occurred on an internal branch. These three branches were subsequently evaluated as candidate patching positions, individually and in combinations.

##### S2.2 ASR responses to individual candidate-branch patching

Patching generally reduced reconstructed ancestral states at nodes upstream of the three candidate branches under PIC-local and the global BM-based methods (ML-BM, GLS-BM, and Rphy-BM; Table S7). At node 149, which lies upstream of both SBLC-$\varphi$ and SBLC-$\omega$, the ML/GLS/Rphy-BM estimate decreased from 11.38 m in the unpatched reconstruction to 10.88 m after patching at $\varphi$ and 10.44 m after patching at $\omega$. Both patched estimates moved toward the Stable-process estimate of 8.97 m and the nearby fossil reference FR14 of 7.41 m. Around SBLC-$\psi$, the PIC-local estimate at node 108 decreased from 2.80 to 2.48 m after patching, approaching the Stable-process estimate of 2.40 m and fossil reference FR6 of 2.23 m. Similar reductions occurred at the more immediate upstream nodes 110 and 111 for SBLC-$\psi$, nodes 152 and 153 for SBLC-$\varphi$, and nodes 152 and 156 for SBLC-$\omega$.

Responses at lateral and descendant nodes depended on the reconstruction framework. Because SBLC-$\varphi$ and SBLC-$\psi$ occurred on terminal branches, neither had an internal descendant node available for ASR; their nearest lateral nodes were 154 and 112, respectively. SBLC-$\omega$ had an internal descendant node, node 157, but no directly corresponding lateral node in the focal set. Patching did not alter PIC-local estimates at these positions: estimates at nodes 112, 154, and 157 remained 1.93, 12.04, and 23.84 m, respectively, across the relevant patched and unpatched reconstructions. The global BM-based methods responded more clearly to partitioning. At node 154, patching at SBLC-$\varphi$ reduced the ML/GLS/Rphy-BM estimate from 14.83 to 13.17 m; at node 112, patching at SBLC-$\psi$ reduced it from 2.34 to 2.11 m; and at the descendant node 157, patching at SBLC-$\omega$ increased the estimate from 19.94 to 23.84 m. Thus, the empirical analysis reproduced the contrast observed in the simulations: local PIC-based estimates at neighboring lateral or descendant positions were comparatively insensitive to partitioning, whereas global reconstructions responded to removal of the candidate branch from the shared covariance structure.

##### S2.3 Spatial and joint effects of candidate-branch patching

The influence of candidate-branch patching under the global BM-based methods was spatially localized rather than phylogeny-wide. Because SBLC-$\varphi$ and SBLC-$\omega$ occurred in nearby regions of the tree, patching at $\varphi$ altered estimates at nodes 156 and 157 near SBLC-$\omega$, whereas patching at $\omega$ altered estimates at nodes 153 and 154 near SBLC-$\varphi$. For example, the ML/GLS/Rphy-BM estimate at node 153 decreased from 15.95 m in the unpatched analysis to 14.45 m after patching at $\omega$, while patching at $\varphi$ changed the estimate at node 157 from 19.94 to 19.36 m. In contrast, patching at $\varphi$ or $\omega$ produced little or no change at nodes 108–112 around the more distant SBLC-$\psi$, and patching at $\psi$ had similarly little effect around SBLC-$\varphi$ and SBLC-$\omega$.

Joint patching produced larger changes where the effects of nearby candidate SBLCs converged on shared upstream regions. At node 149, the ML/GLS/Rphy-BM estimate was 10.88 m after patching only at $\varphi$ and 10.44 m after patching only at $\omega$, but decreased to 9.66 m when both branches were patched. At node 152, the corresponding estimates were 14.36 m after patching at $\varphi$, 13.16 m after patching at $\omega$, and 11.02 m after joint $\varphi + \omega$ patching. Adding the phylogenetically more distant SBLC-$\psi$ as a third cut point produced virtually no further change in the region upstream of $\varphi$ and $\omega$, with a difference of only 0.01 m at a single node. These results are consistent with partially additive effects of nearby candidate SBLCs on shared upstream ancestral estimates and with attenuation of their influence over greater phylogenetic separation.

Bayes-MCMC showed a less regular empirical response. Changes in posterior point estimates varied in both magnitude and direction among nodes and patching configurations, without a consistent upstream, lateral, or distance-dependent pattern comparable to those observed for PIC-local and the global BM-based methods. The Bayesian results therefore indicate greater method-specific variability rather than a common directional response to candidate-branch patching.

### Supplementary Figures

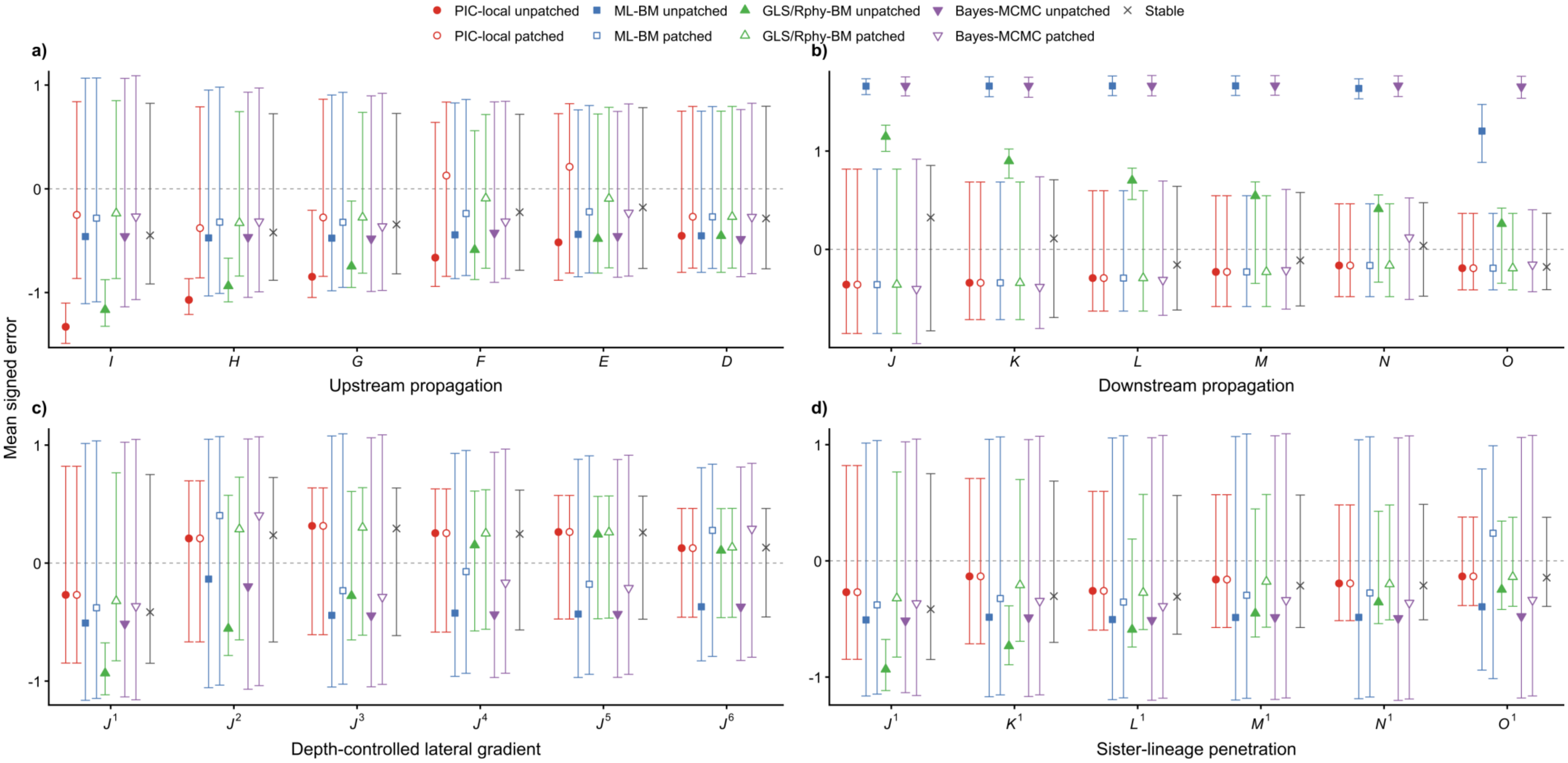

**Figure S1. Propagation of SBLC-induced reconstruction errors across phylogenetic distances.**

Results are shown for a downsizing SBLC ($k$ = 1/16) on a balanced ultrametric tree with 4,096 terminal taxa and a descendant subtree ($T_1$) of 64 taxa. Reconstruction errors were examined across six successive node positions in four topological directions relative to the SBLC-bearing *I*–*J* branch: (a) upstream propagation from *I* toward the root (*I*–*D*); (b) downstream propagation from *J* within $T_1$ (*J*–*O*); (c) a depth-controlled lateral series ($J^1$–$J^6$); and (d) progressive penetration into the sister lineage ($J^1$–$O^1$). Node nomenclature and topological relationships follow the scheme illustrated in Figure 1 in the main text. Points show mean signed reconstruction errors, with error bars indicating 95% bootstrap confidence intervals. Numerical values are reported in Supplementary Table S5.

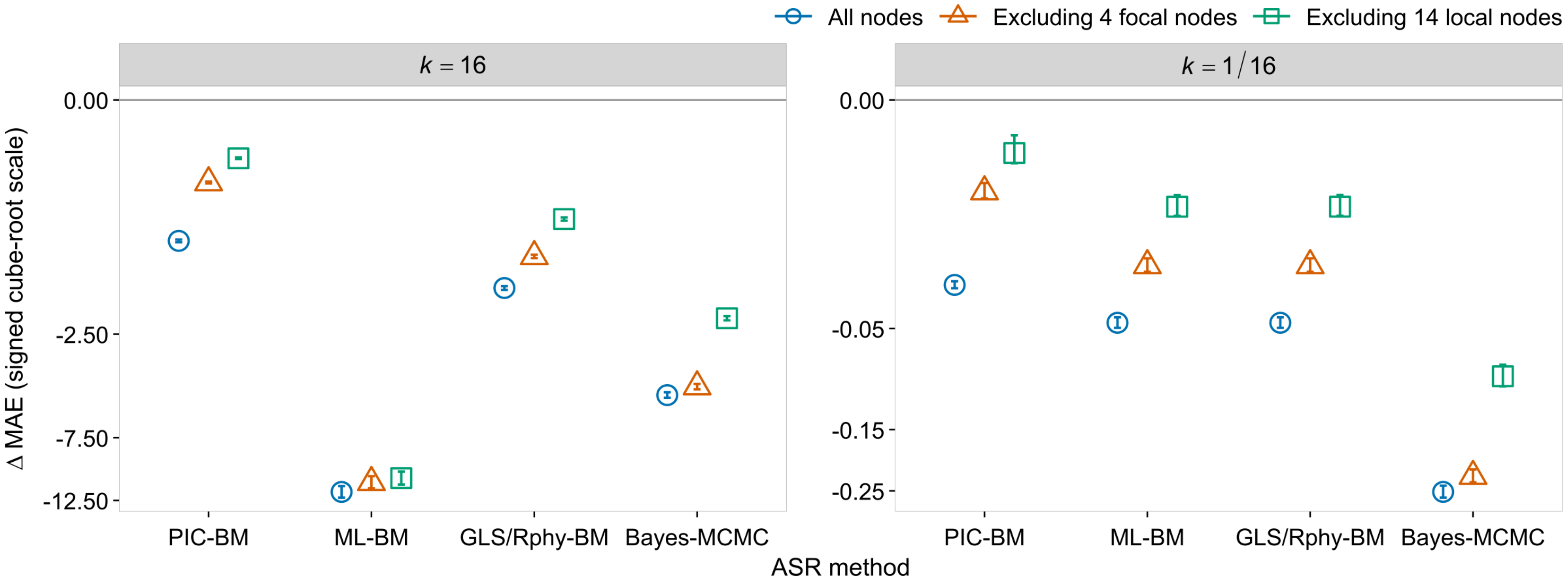

**Figure S2. Tree-wide changes in mean absolute ancestral-state reconstruction (ASR) error after patching.**

Points show $\overline{\Delta\text{MAE}}$, the mean across 100 independent trait realizations of the within-realization difference between patched and unpatched mean absolute error (patched minus unpatched). Results are shown for an upsizing SBLC with $k$ = 16 (left) and a downsizing SBLC with $k$ = 1/16 (right). MAE was calculated across three internal-node sets: all 127 internal nodes, including the root; the 123 nodes remaining after exclusion of the four focal nodes ($H$, $I$, $J$, and $J^1$); and the 113 nodes remaining after exclusion of a broader set of 14 local nodes. In the broader local-exclusion analysis, the two endpoints of the SBLC-bearing branch and every internal node within two branching steps of either endpoint were excluded, removing 14 nodes in total and leaving 113 internal nodes for analysis. Error bars indicate 95% confidence intervals obtained from 10,000 nonparametric bootstrap resamples of the complete trait realizations. Negative values indicate lower MAE after patching, and the horizontal line at zero indicates no mean difference. The vertical axis uses a signed cube-root transformation, with tick labels shown on the original $\overline{\Delta\text{MAE}}$ scale. GLS-BM and Rphy-BM produced identical results and are shown together.

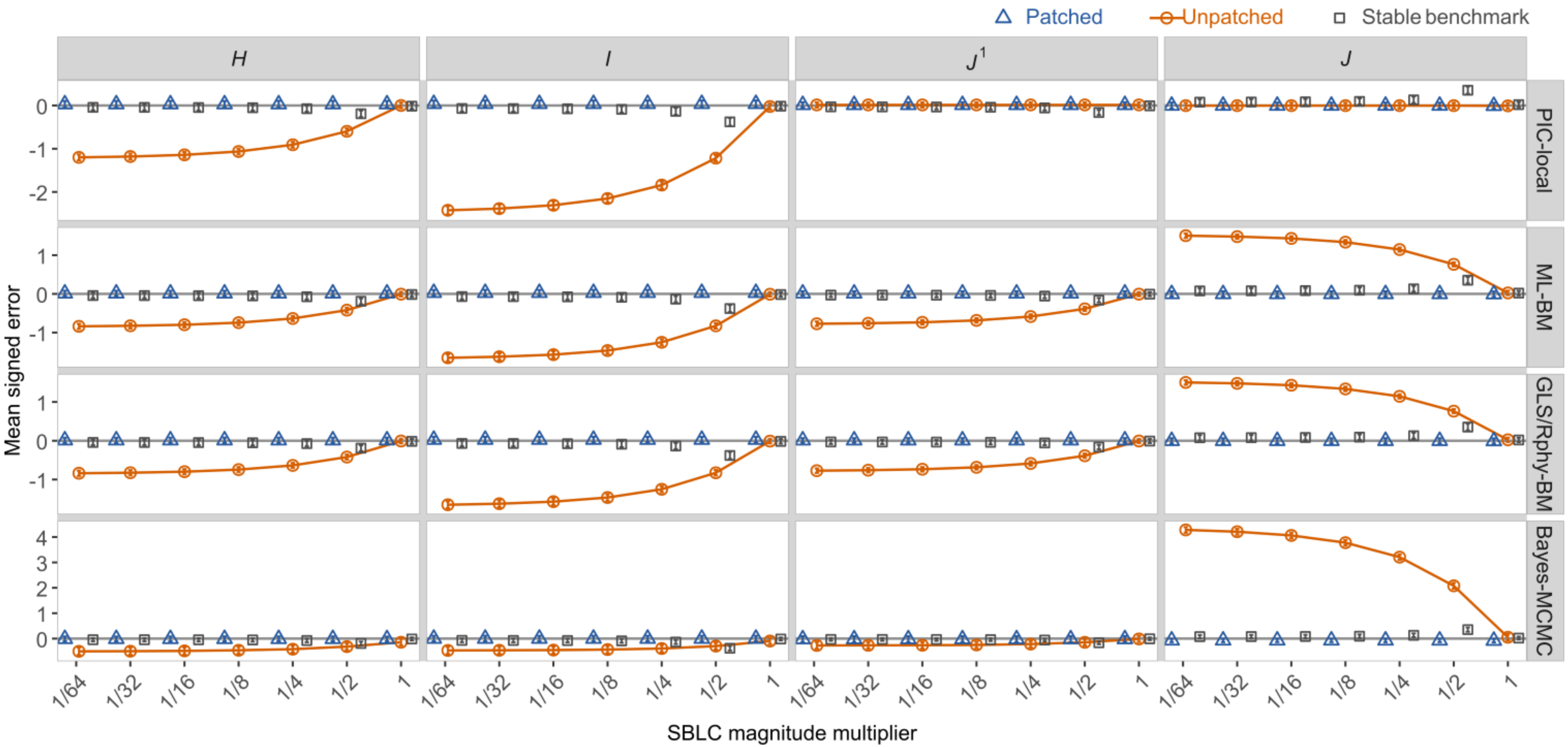

**Figure S3. Effects of downsizing SBLC magnitude on ASR error before and after patching.**

Mean signed reconstruction errors are shown across reciprocal SBLC multipliers ranging from $k = 1$, representing no imposed SBLC, to $k = 1/64$, representing the strongest downsizing change examined. Smaller values of $k$ therefore indicate increasingly strong downsizing SBLCs. Columns correspond to focal nodes defined relative to the SBLC-bearing branch. From top to bottom, rows show PIC-local, ML-BM, GLS/Rphy-BM, and Bayes-MCMC reconstructions. Circles connected by solid lines show unpatched reconstructions, whereas triangles and squares show patched and stable-process reconstructions, respectively. Stable-process reconstruction was included as an external benchmark and was not subjected to patching. Points represent mean signed errors across 100 independent simulation replicates, and error bars indicate bootstrap 95% confidence intervals. Positive and negative values indicate overestimation and underestimation, respectively. GLS-BM and Rphy-BM produced identical mean signed errors and are therefore displayed together as GLS/Rphy-BM.

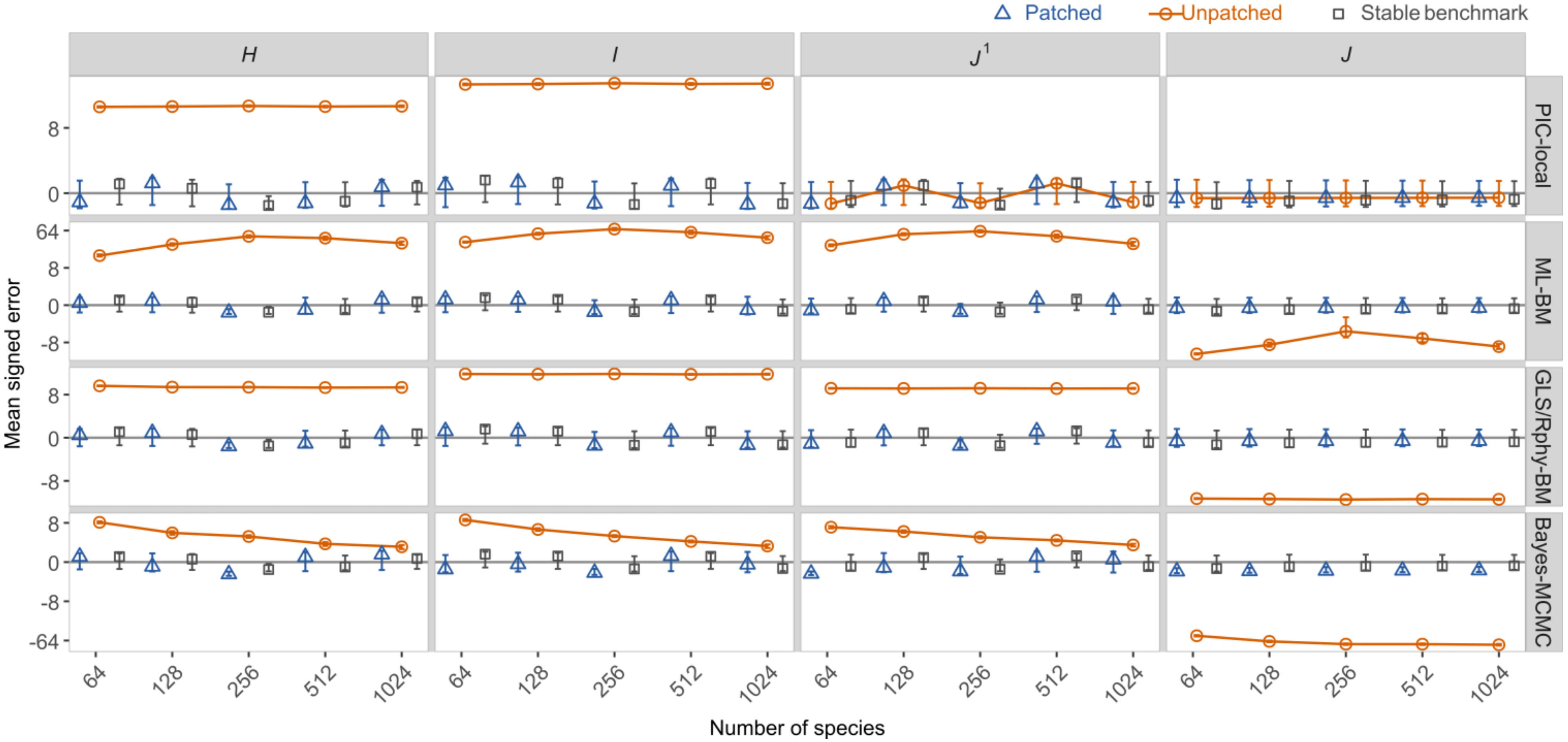

**Figure S4. Sensitivity of ASR error to phylogenetic tree size under an upsizing SBLC.**

Mean signed reconstruction errors are shown for balanced ultrametric trees containing 64, 128, 256, 512, or 1,024 terminal taxa under a fixed upsizing SBLC multiplier of $k = 16$. All trees were scaled to unit root-to-tip height, and the descendant subtree rooted at *J* contained eight terminal taxa. Columns correspond to focal nodes defined relative to the SBLC-bearing branch. From top to bottom, rows show PIC-local, ML-BM, GLS/Rphy-BM, and Bayes-MCMC reconstructions. Circles connected by solid lines show unpatched reconstructions, whereas triangles and squares show patched and stable-process reconstructions, respectively. Stable-process reconstruction was included as an external benchmark and was not subjected to patching. Points represent mean signed errors across 100 independent simulation replicates, and error bars indicate bootstrap 95% confidence intervals. Positive and negative values indicate overestimation and underestimation, respectively. The y-axis is displayed on a signed cube-root scale to accommodate differences in error magnitude while preserving direction; tick labels report signed errors on the original scale. GLS-BM and Rphy-BM produced identical mean signed errors and are therefore displayed together as GLS/Rphy-BM.

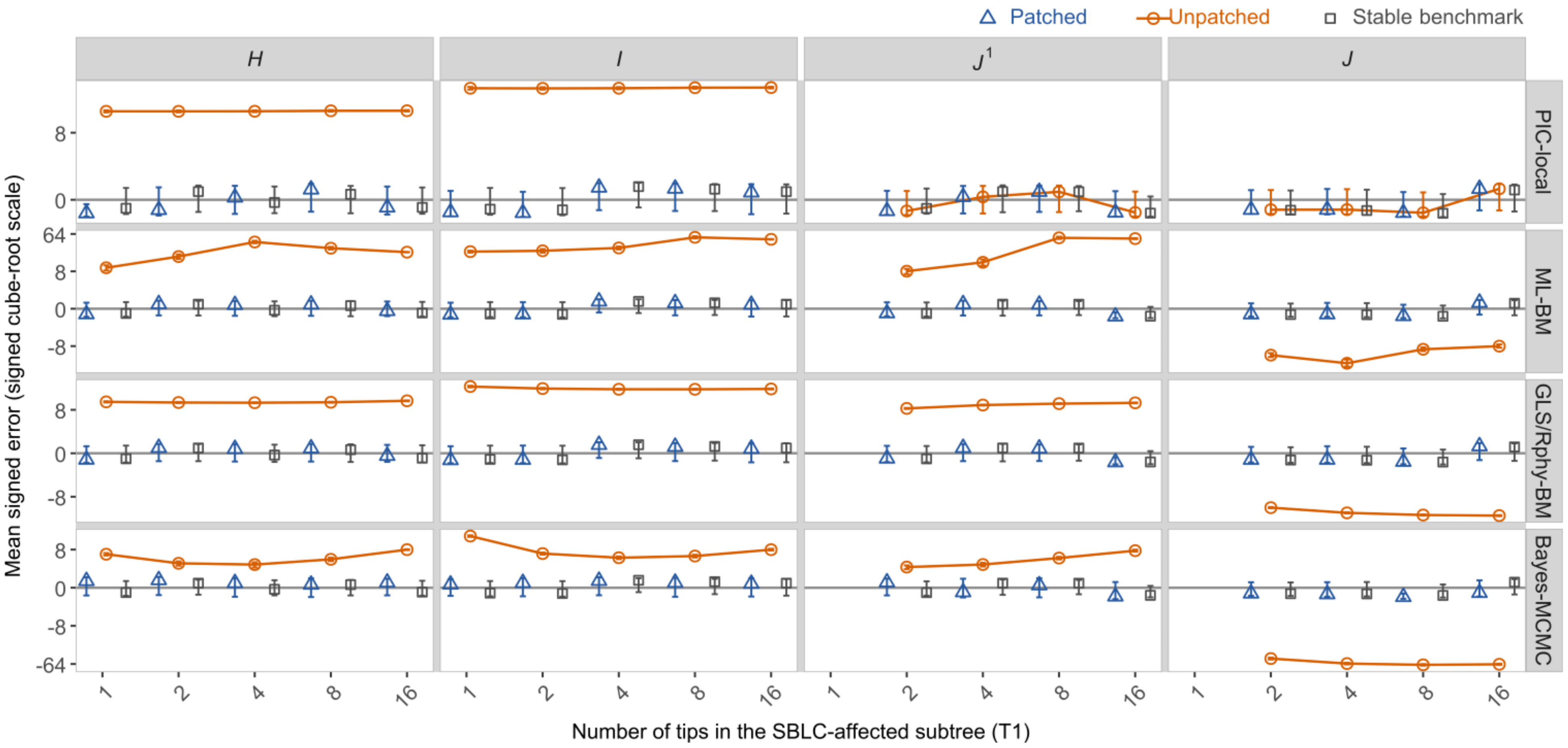

**Figure S5. Sensitivity of ASR error to descendant-subtree size under an upsizing SBLC.**
Signed reconstruction errors are shown for an upsizing SBLC of $k = 16$ as the size of the affected descendant subtree ($T_1$) varied from 1 to 16 terminal taxa. Results are presented for the four focal nodes defined relative to the SBLC-bearing branch: the upstream boundary node *I*, its successive ancestor *H*, the descendant-side boundary node *J*, and the lateral-lineage node $J^1$. Circles represent unpatched reconstructions, triangles represent patched reconstructions, and squares represent the stable-process benchmark. Points and error bars indicate the mean signed error and its 95% confidence interval, respectively. The horizontal reference line denotes zero reconstruction error. Positive and negative values indicate overestimation and underestimation of the true ancestral state, respectively.

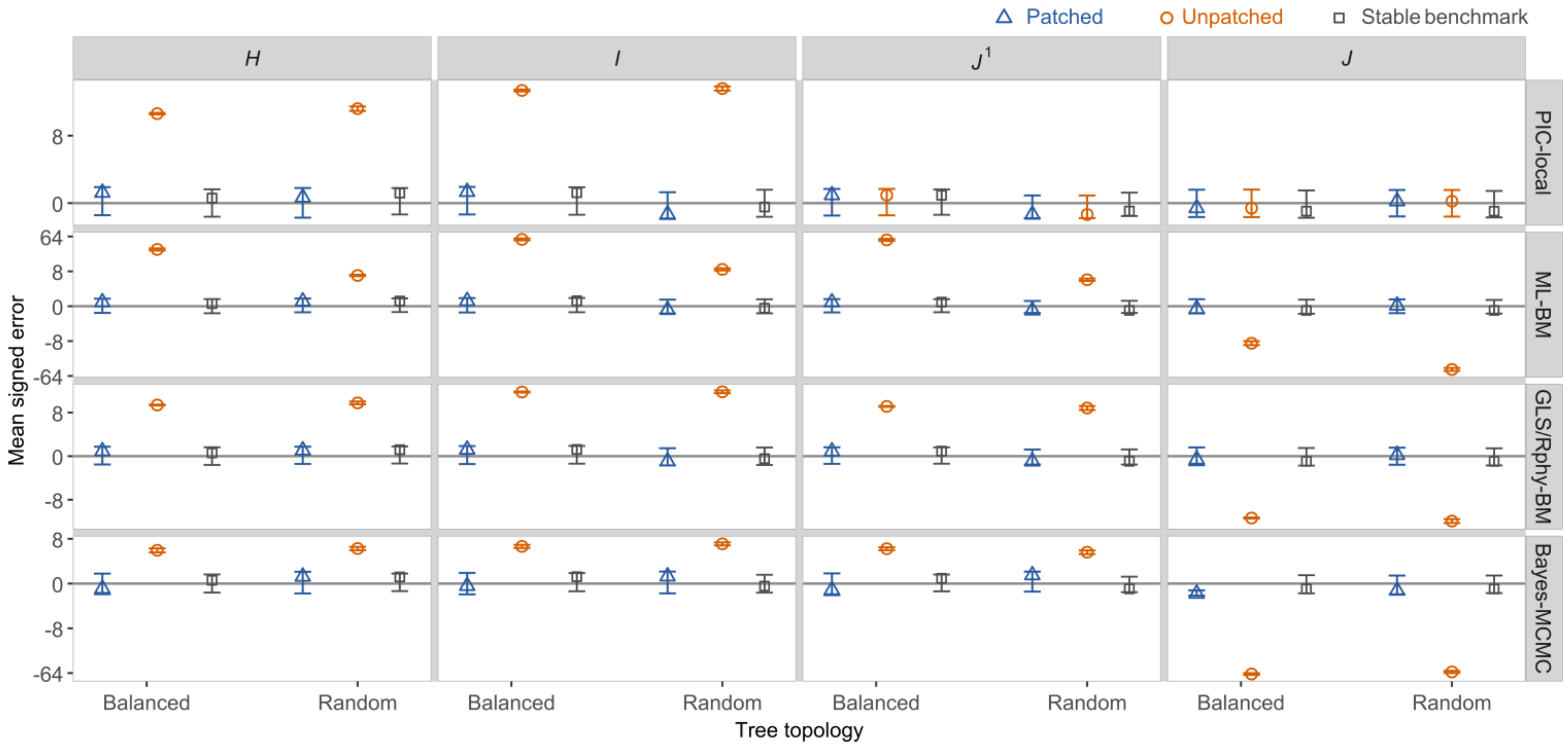

**Figure S6. Effects of random ultrametric tree topology on ASR under an upsizing SBLC.** Mean signed reconstruction errors are shown for the fixed balanced topology used in the primary analysis and for randomly generated ultrametric trees produced under a pure-birth Yule process. All trees contained 128 terminal taxa and were scaled to unit root-to-tip height. The SBLC-bearing branch subtended a descendant subtree containing eight terminal taxa, and traits were simulated under the BM background model with an upsizing SBLC of $k = 16$. Rows represent reconstruction methods, and columns represent the SBLC-centered focal nodes $H$, $I$, $J^1$, and $J$. Circles show unpatched reconstructions, triangles show patched reconstructions, and squares show stable-process reconstructions, which served as an external benchmark and were not patched. Points represent mean signed errors, and error bars indicate bootstrap 95% confidence intervals. To accommodate the large range of positive and negative errors, the vertical axis is displayed using a signed cube-root scale; tick labels denote values on the original signed-error scale.

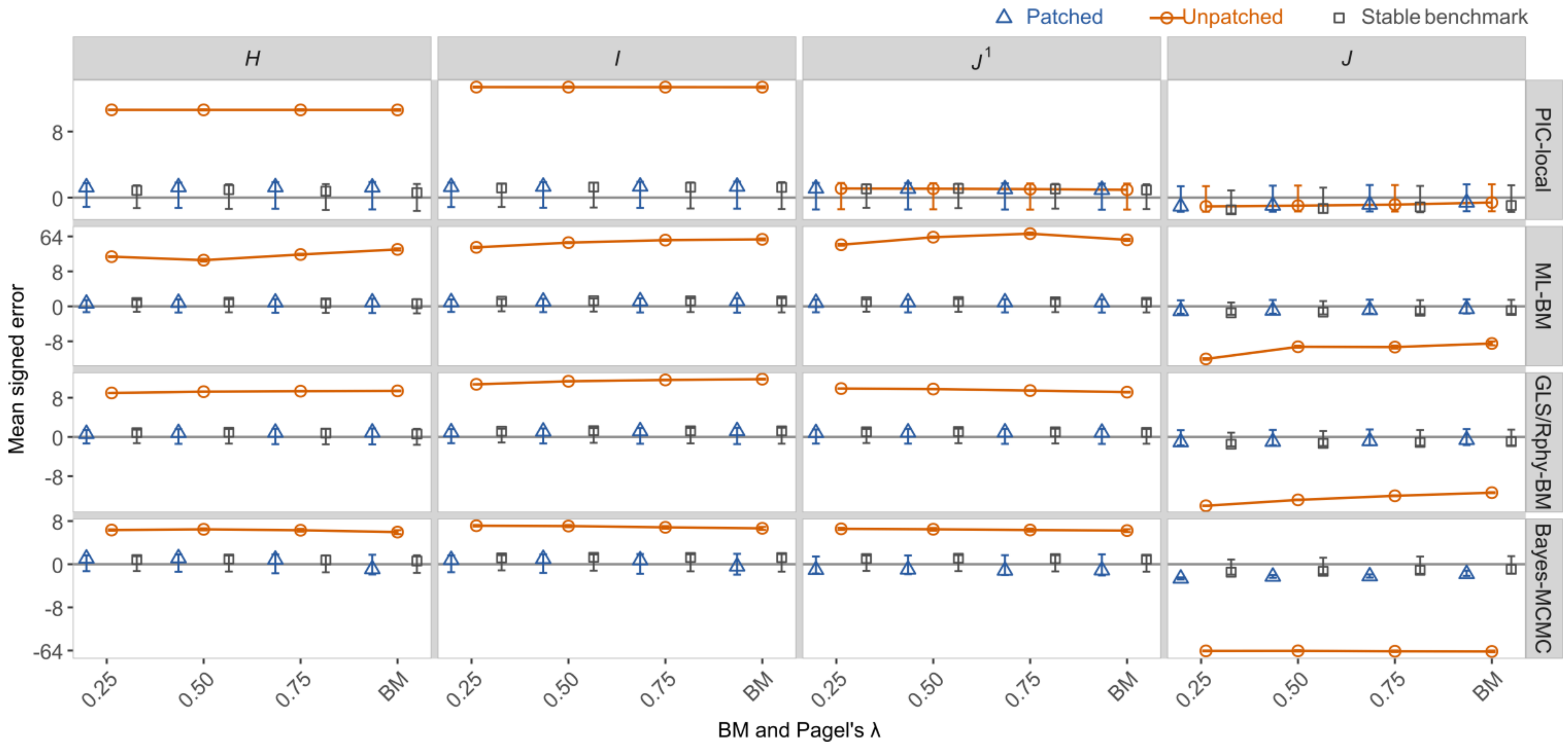

**Figure S7. Sensitivity of ASR error to Pagel's λ under an upsizing SBLC.**
Mean signed reconstruction errors are shown across Pagel's $\lambda$ settings of 0.25, 0.50, and 0.75, with standard BM included as the $\lambda = 1$ reference. The upsizing SBLC used $k = 16$. Circles denote unpatched reconstructions, triangles denote patched reconstructions, and squares denote the stable-process benchmark. Lines connect unpatched estimates across model settings to aid visualization. Points represent mean signed errors across 100 independent simulation replicates, and error bars indicate bootstrap 95% confidence intervals. Positive and negative values indicate overestimation and underestimation, respectively. Because of the large range of errors under the upsizing SBLC, the vertical axis uses the sign-preserving cube-root transformation, with tick labels expressed in the original signed-error units. GLS-BM and Rphy-BM are shown together as GLS/Rphy-BM because their estimates overlap at the plotted resolution. All other settings matched the primary simulation, including the balanced 128-tip ultrametric tree and the eight-tip SBLC-affected descendant subtree ($T_1 = 8$). Focal nodes are defined relative to the SBLC-bearing branch as illustrated in Figure 1 in the main text.

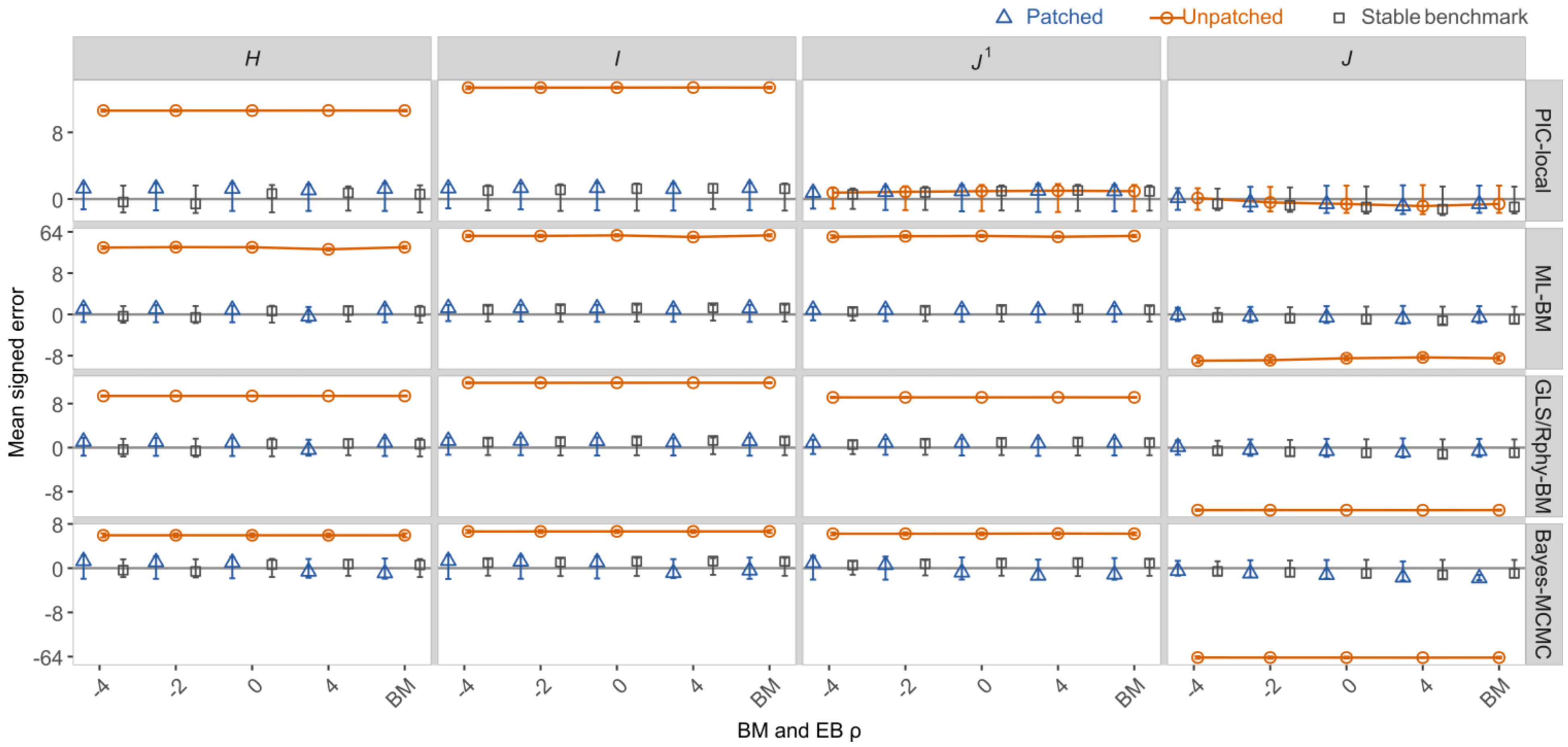

**Figure S8. Sensitivity of ASR error to the early-burst (EB) background model under an upsizing SBLC.** Mean signed reconstruction errors are shown across variance-normalized EB settings of $\rho = -4, -2, 0$, and 4, with the original BM simulation included as a separate reference. Negative $\rho$ values concentrate a greater proportion of the matched total evolutionary variance on deeper branches, whereas positive values concentrate more variance on recent branches; $\rho = 0$ represents the no-rate-heterogeneity EB setting and is theoretically equivalent to BM. The upsizing SBLC used $k = 16$. Circles denote unpatched reconstructions, triangles denote patched reconstructions, and squares denote the stable-process benchmark. Lines connect unpatched estimates across model settings to aid visualization. Points represent mean signed errors across 100 independent simulation replicates, and error bars indicate bootstrap 95% confidence intervals. Positive and negative values indicate overestimation and underestimation, respectively. The vertical axis uses the sign-preserving cube-root transformation, with tick labels expressed in the original signed-error units. GLS-BM and Rphy-BM are shown together as GLS/Rphy-BM because their estimates overlap at the plotted resolution. All other settings matched the primary simulation, including the balanced 128-tip ultrametric tree and $T_1 = 8$. Focal nodes are defined relative to the SBLC-bearing branch as illustrated in Figure 1 in the main text.

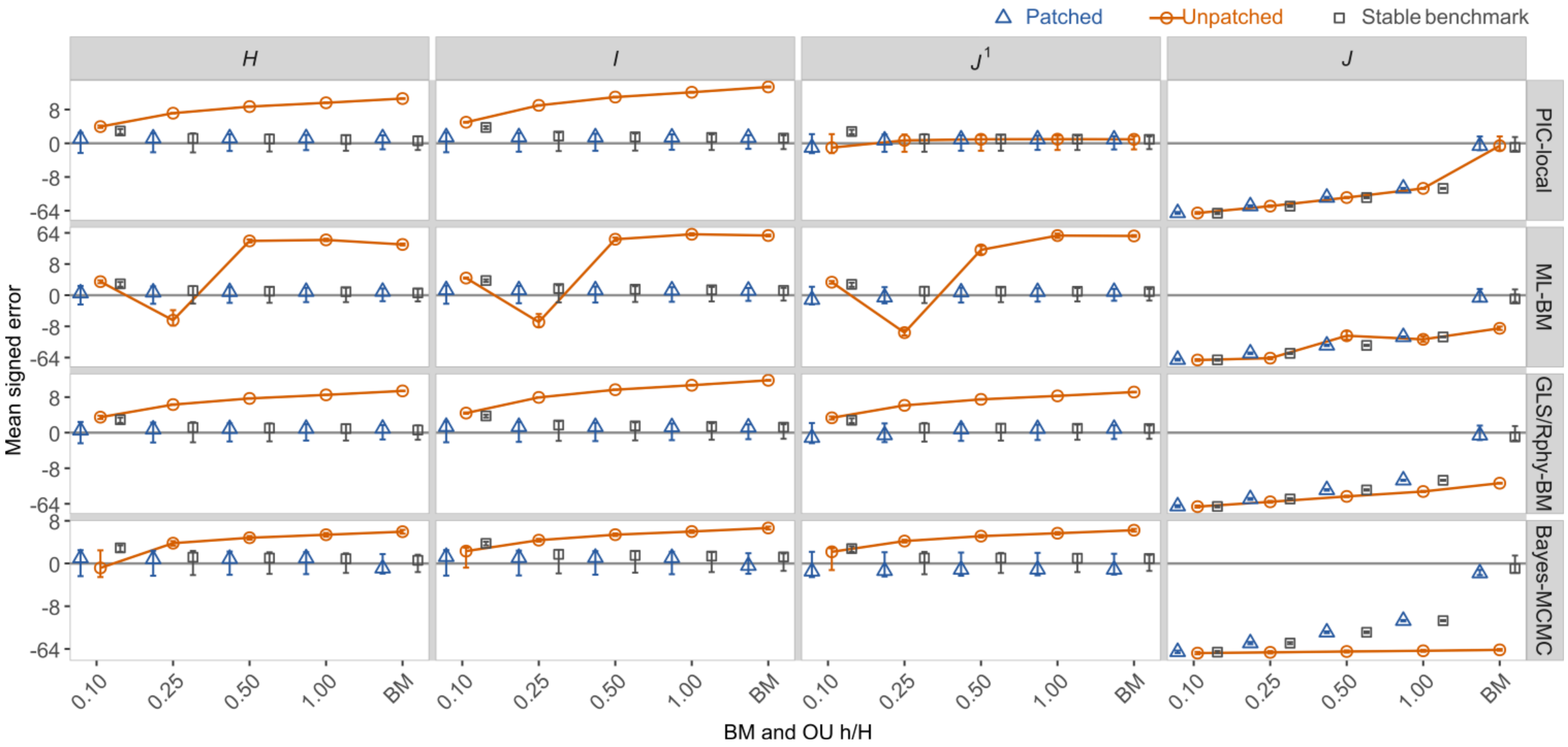

**Figure S9. Sensitivity of ASR error to Ornstein-Uhlenbeck (OU) attraction under an upsizing SBLC.** Mean signed reconstruction errors are shown across variance-normalized OU settings with phylogenetic half-life ratios $h/H$ = 0.10, 0.25, 0.50, and 1.00, with standard BM included as a reference. Smaller $h/H$ values indicate stronger attraction toward the optimum and more rapid loss of phylogenetic memory. The upsizing SBLC used $k$ = 16. Circles denote unpatched reconstructions, triangles denote patched reconstructions, and squares denote the stable-process benchmark. Lines connect unpatched estimates across model settings to aid visualization. Points represent mean signed errors across 100 independent simulation replicates, and error bars indicate bootstrap 95% confidence intervals. Positive and negative values indicate overestimation and underestimation, respectively. The vertical axis uses the sign-preserving cube-root transformation, with tick labels expressed in the original signed-error units. GLS-BM and Rphy-BM are shown together as GLS/Rphy-BM because their estimates overlap at the plotted resolution. All other settings matched the primary simulation, including the balanced 128-tip ultrametric tree and $T_1$ = 8. Focal nodes are defined relative to the SBLC-bearing branch as illustrated in Figure 1 in the main text.

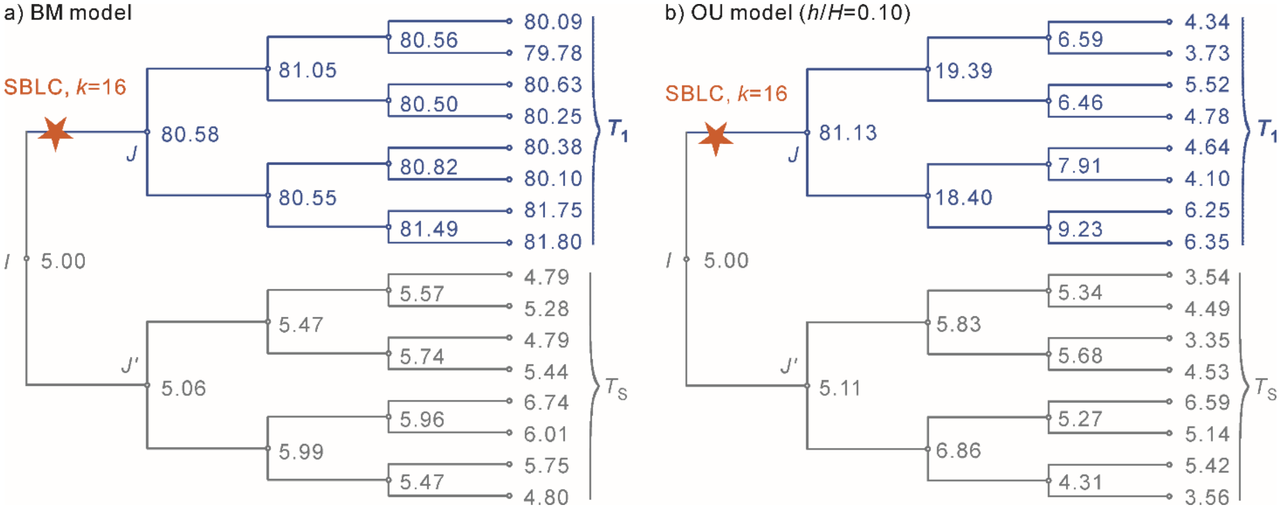

**Figure S10. Illustrative persistence and attenuation of the descendant signal of an upsizing SBLC under BM and strong OU evolution.**
True simulated states from a single prespecified replicate are shown for the eight-tip SBLC-affected descendant subtree $T_1$ and its eight-tip sister reference subtree $T_S$ under (a) Brownian motion (BM) and (b) Ornstein–Uhlenbeck (OU) evolution with $h/H = 0.10$. The upsizing SBLC ($k = 16$) produced comparable immediate post-SBLC states at the descendant-side boundary node $J$ under the two backgrounds ($X_J = 80.58$ under BM and $X_J = 81.13$ under OU). Under BM, the resulting displacement remained evident throughout $T_1$ and at its terminal taxa. Under strong OU attraction, the displacement was progressively attenuated along the descendant branches, and the terminal states returned toward the optimum ($\theta = 5$), thereby retaining little information about the transient post-SBLC state at $J$. For a descendant at distance $d$ from $J$, the expected proportion of the initial displacement retained under OU is $R(d) = \exp(-\alpha d) = 2^{-d/h}$; at the terminal depth represented here, approximately 5.1% of the displacement is expected to remain, compared with 100% under BM. $T_S$ did not descend from the SBLC-bearing branch and is shown as a local phylogenetically matched reference; it forms part of, but is not equivalent to, the remaining tree $T$ defined for the patch procedure. Node labels are true simulated states rather than reconstructed ancestral states and are presented to illustrate the generative mechanism underlying the reconstruction-error patterns in Figure S9 and Repository Appendix A, Figure A6.

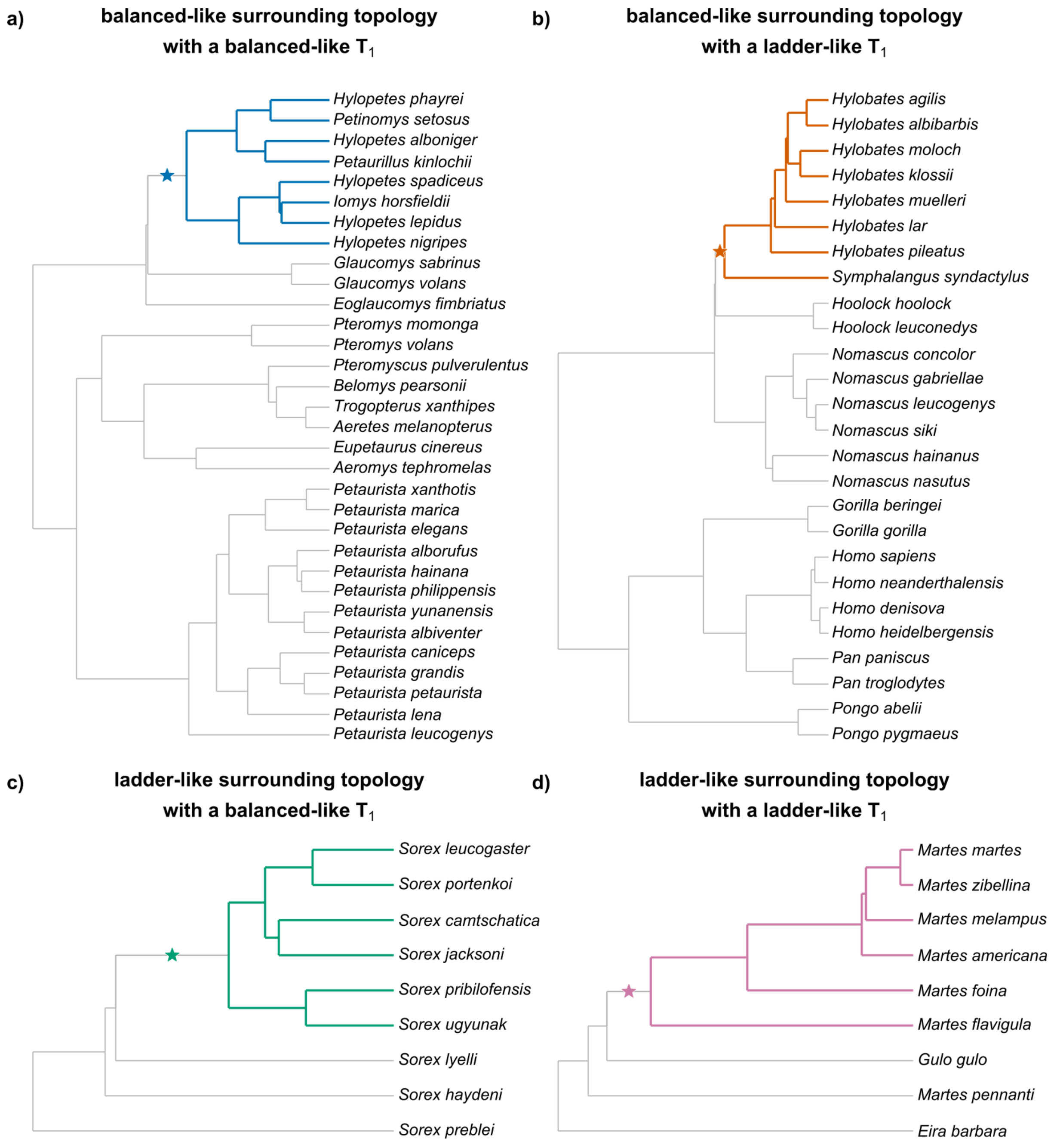

**Figure S11. Representative SBLC placements under contrasting local and surrounding topological configurations on the empirical mammalian phylogeny.**

The full empirical phylogeny contained 4,099 mammalian species; for clarity, only the local phylogenetic region surrounding each selected SBLC-bearing branch is shown. Four configurations were defined by combining the topological balance of the SBLC-descendant subtree $T_1$ with that of the surrounding phylogenetic context: (a) balanced-like surrounding topology with a balanced-like $T_1$ (class $\alpha$); (b) balanced-like surrounding topology with a ladder-like $T_1$ (class $\beta$); (c) ladder-like surrounding topology with a balanced-like $T_1$ (class $\gamma$); and (d) ladder-like surrounding topology with a ladder-like $T_1$ (class $\delta$). The selected $\alpha$ and $\beta$ placements each contained eight descendant species in $T_1$, whereas the selected $\gamma$ and $\delta$ placements each contained six. The SBLC-bearing branch is marked by a star in each panel, and the descendant subtree below this branch corresponds to $T_1$. Topological categories were defined using normalized Colless indices as described in Supplementary Methods Section S3.7.2.

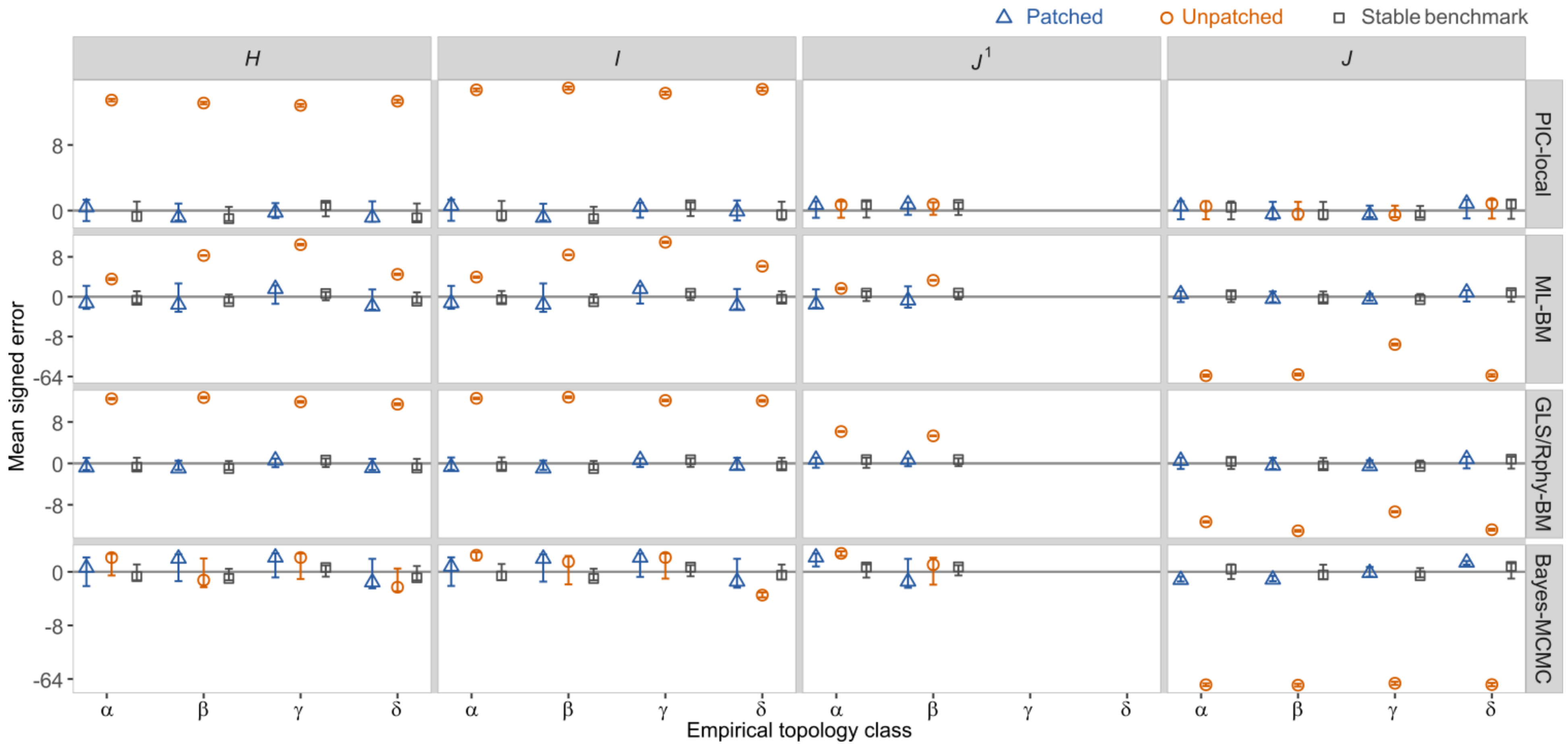

**Figure S12. Effects of empirical local topology on ASR under an upsizing SBLC.**

Mean signed reconstruction errors are shown for four representative SBLC placements on the empirical mammalian phylogeny under an upsizing SBLC of $k$ = 16. The four topology classes combine the balance of the surrounding phylogenetic context with that of the SBLC-descendant subtree $T_1$: $\alpha$, balanced-like surrounding topology with a balanced-like $T_1$; $\beta$, balanced-like surrounding topology with a ladder-like $T_1$; $\gamma$, ladder-like surrounding topology with a balanced-like $T_1$; and $\delta$, ladder-like surrounding topology with a ladder-like $T_1$. Columns correspond to the focal nodes $H$, $I$, $J^1$, and $J$, and rows show PIC-local, ML-BM, GLS/Rphy-BM, and Bayes-MCMC reconstructions. Circles denote unpatched reconstructions, triangles denote patched reconstructions, and squares denote the Stable-process benchmark. Points represent mean signed errors across 100 independent simulation replicates, and error bars indicate bootstrap 95% confidence intervals. Positive and negative values indicate overestimation and underestimation, respectively. $J^1$ is not shown for classes $\gamma$ and $\delta$ because, in those empirical placements, the sister lineage of $T_1$ is a terminal branch and therefore contains no internal node at which ancestral-state reconstruction can be evaluated. GLS-BM and Rphy-BM are shown together as GLS/Rphy-BM because their estimates overlap at the plotted resolution.

**Figure S13. Empirical-phylogeny placements used to evaluate the effect of post-SBLC evolutionary duration under OU evolution.** The full empirical phylogeny contained 4,099 mammalian species; for clarity, only the local phylogenetic regions surrounding the three selected SBLC-bearing branches are shown. Panels (a–c) represent placements with short, intermediate, and long post-SBLC evolutionary durations, respectively. All three placements had $T_1 = 8$ and similar $I \rightarrow J$ branch lengths and $T_1$ topological balance, while differing substantially in the age of the descendant-side boundary node $J$. The SBLC-bearing branch is indicated by a star, and the descendant subtree extending from $J$ corresponds to $T_1$. To facilitate visual comparison of post-SBLC evolutionary duration, node $J$ was aligned at time zero in all three panels, as indicated by the vertical dashed line; branch positions to the left and right therefore represent evolutionary time before and after $J$, respectively. Branch lengths are shown here on the original time-calibrated scale (Ma) to illustrate differences in post-SBLC evolutionary duration among placements; the corresponding trees were rescaled to unit root-to-tip height for trait simulation. Larger surrounding sister clades were collapsed for display only, with the numbers of descendant species indicated in parentheses. The complete empirical topology and branch lengths were retained in all simulations.

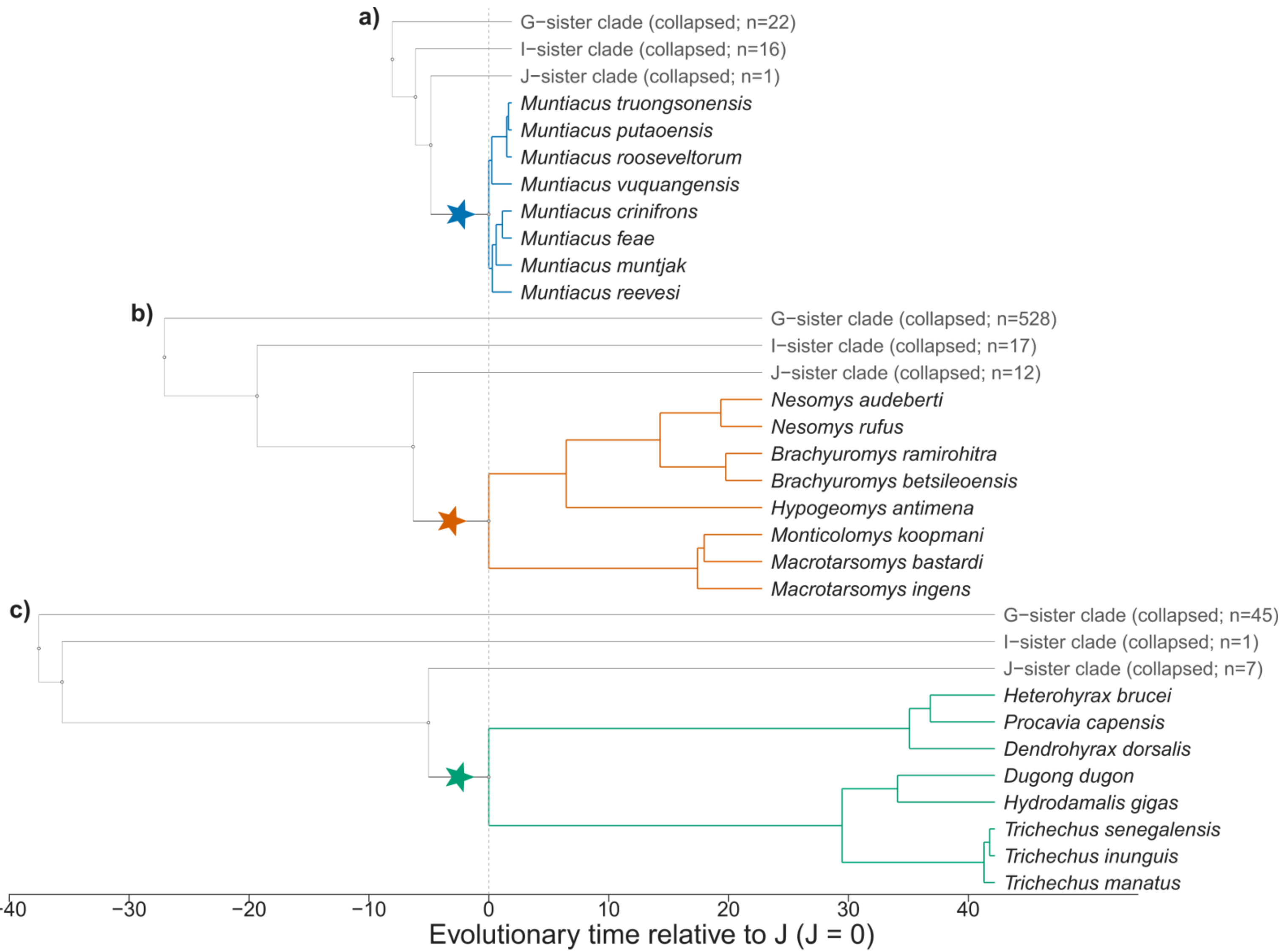

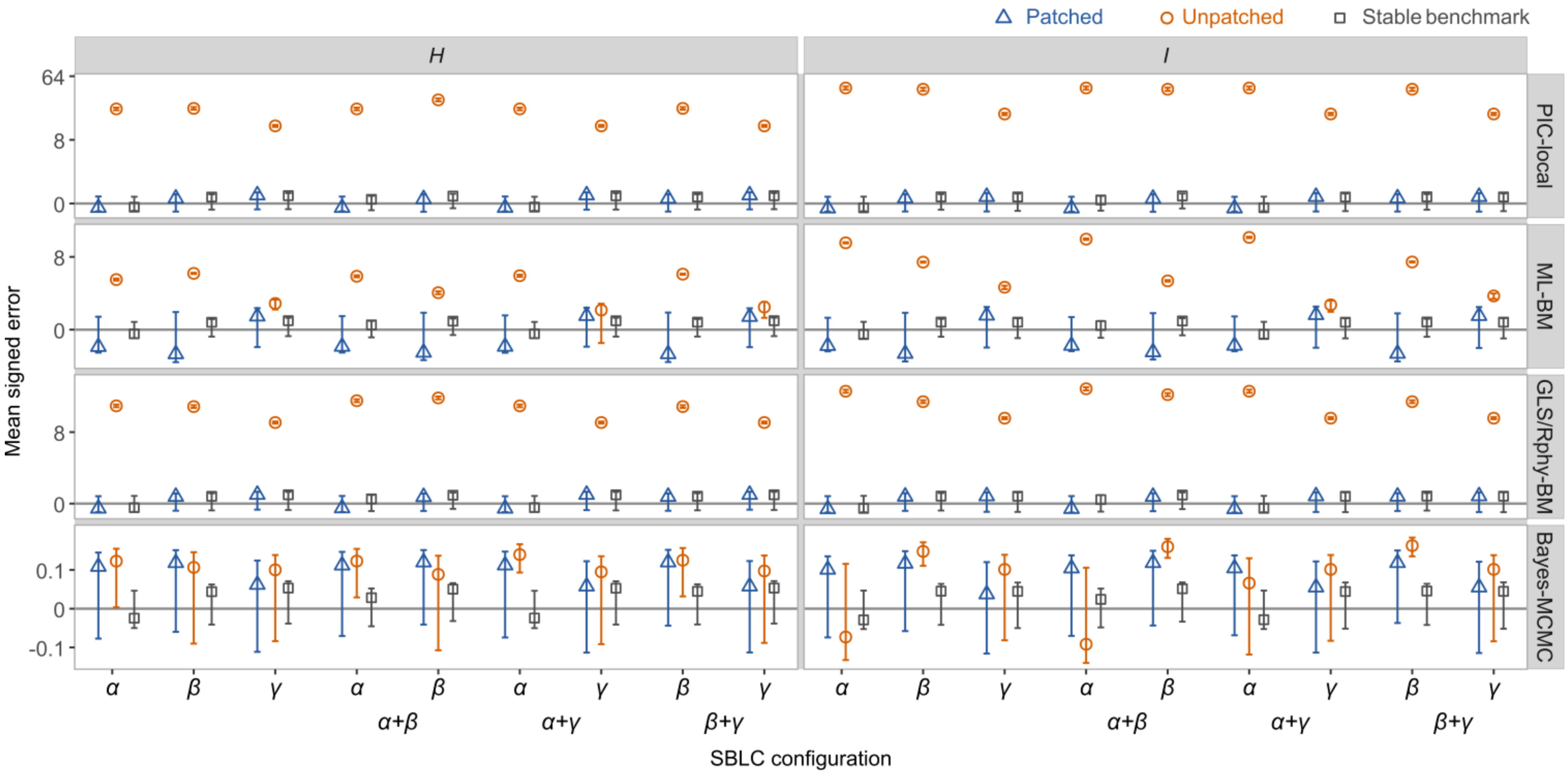

**Figure S14. Effects of single and paired upsizing SBLCs on reconstruction error at the upstream nodes *H* and *I*.**
Mean signed reconstruction errors are shown for an upsizing SBLC ($k$ = 16) introduced separately at placements $\alpha$, $\beta$, or $\gamma$, or jointly at the nearby pair $\alpha + \beta$ and the two distant pairs $\alpha + \gamma$ and $\beta + \gamma$. Within each paired configuration, the lower-axis labels identify the focal SBLC whose associated nodes are evaluated. Columns represent the upstream nodes *H* and *I*, and rows represent PIC-local, ML-BM, GLS/Rphy-BM, and Bayes-MCMC. Circles show conventional reconstructions on the intact phylogeny, triangles show patched reconstructions, and squares show the stable-process benchmark. For a single-SBLC scenario, patching partitioned the phylogeny at the single SBLC-bearing branch; for a paired-SBLC scenario, joint patching partitioned it at both SBLC-bearing branches. Points represent mean signed errors across 100 independently simulated trait realizations, and error bars indicate bootstrap 95% confidence intervals. The horizontal line denotes zero reconstruction error; positive and negative values indicate overestimation and underestimation, respectively. The vertical axis uses a signed cube-root transformation, with tick labels shown on the original error scale; displayed ranges vary among method rows. GLS-BM and Rphy-BM produced identical mean signed errors and are therefore displayed together as GLS/Rphy-BM. The stable-process reconstructions served as an external benchmark and were not patched.

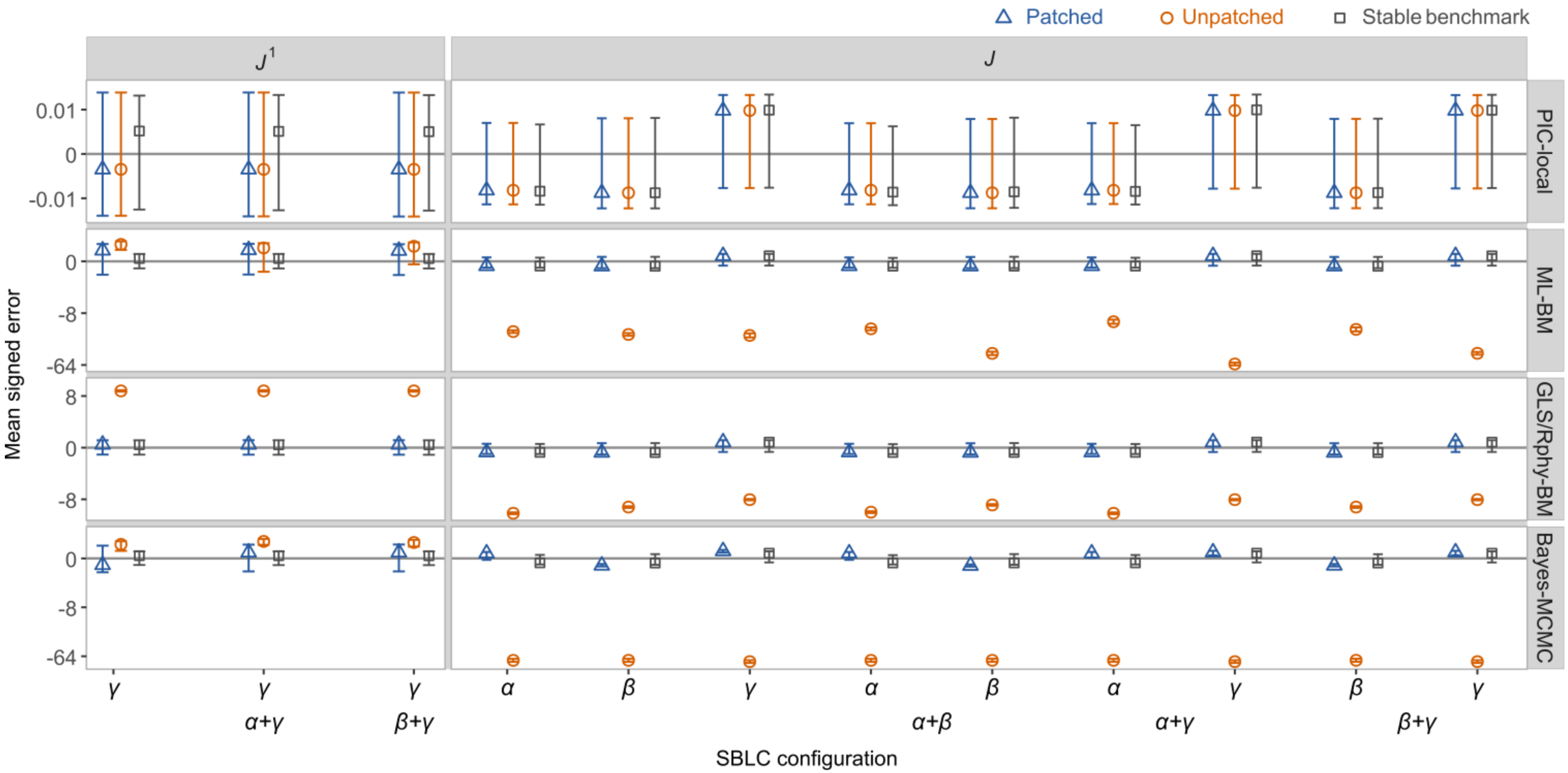

**Figure S15. Effects of single and paired upsizing SBLCs on reconstruction error at the descendant-side and adjacent lateral nodes.** Mean signed reconstruction errors are shown at the descendant-side boundary node $J$ and the adjacent lateral node $J^1$ for an upsizing SBLC ($k$ = 16). SBLCs were introduced separately at $\alpha$, $\beta$, or $\gamma$, or jointly at the nearby pair $\alpha + \beta$ and the two distant pairs $\alpha + \gamma$ and $\beta + \gamma$. Within each paired configuration, the lower-axis labels identify the focal SBLC whose associated node is evaluated. Results at $J$ are shown for all three single placements and for both focal SBLCs within each pair. The $J^1$ panel is restricted to the $\gamma$-associated focal node and therefore shows $\gamma$ alone and $\gamma$ within the $\alpha + \gamma$ and $\beta + \gamma$ configurations. Rows represent PIC-local, ML-BM, GLS/Rphy-BM, and Bayes-MCMC. Circles show conventional reconstructions on the intact phylogeny, triangles show patched reconstructions, and squares show the stable-process benchmark. Patching was performed at the single affected branch in the single-SBLC scenarios and jointly at both affected branches in the paired-SBLC scenarios. Points represent mean signed errors across 100 independently simulated trait realizations, and error bars indicate bootstrap 95% confidence intervals. The horizontal line denotes zero reconstruction error. The vertical axis uses a signed cube-root transformation, with tick labels shown on the original error scale; displayed ranges vary among method rows. GLS-BM and Rphy-BM are displayed together because they produced identical mean signed errors. The stable-process reconstructions served as an external benchmark and were not patched.

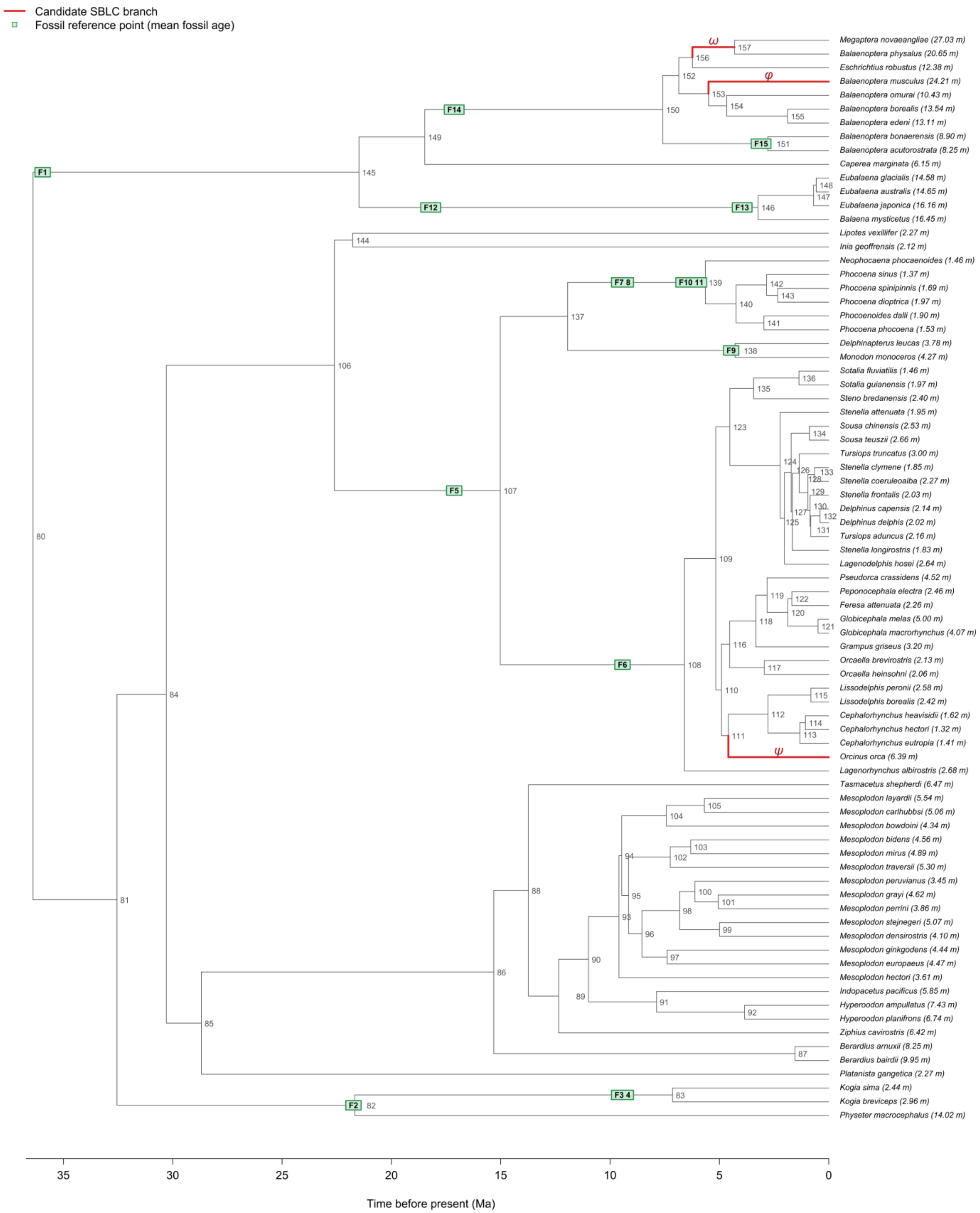

**Figure S16. Empirical cetacean phylogeny showing candidate SBLC branches and fossil reference points.** Time-calibrated phylogeny of the 79 extant cetacean species used for empirical ASR and Lévy-process analyses. Thick branches represent the three candidate SBLCs identified from the Lévy-process analysis and are labeled $\varphi$, $\psi$, and $\omega$ in descending order of absolute signal-to-noise ratio. All three candidate SBLCs represented inferred increases in body length. Symbols placed along branches represent the 15 fossil reference points retained after topological matching and temporal consistency screening; their positions correspond to fossil mean ages projected onto the associated branches and do not represent additional terminal taxa in the molecular phylogeny. Extant species labels include mean total body length (m).

### Supplementary Tables

#### Table S1. Node-wise signed error under representative SBLC scenarios

| Node | PIC-local | ML-BM | GLS-BM | Rphy-BM | Bayes-MCMC |
|---|---|---|---|---|---|
| Upsizing single-branch large change (SBLC), *k* = 16 | | | | | |
| *I* | 37.355 (36.357, 38.345) | 56.203 (53.783, 58.778) | 25.550 (24.900, 26.204) | 25.550 (24.886, 26.204) | 4.597 (4.023, 5.184) |
| *H* | 18.689 (18.155, 19.202) | 34.516 (32.517, 36.529) | 12.944 (12.591, 13.286) | 12.944 (12.591, 13.289) | 3.302 (2.707, 3.913) |
| $J^1$ | **0.013 (-0.048, 0.073)** | 55.000 (52.564, 57.565) | 11.921 (11.593, 12.243) | 11.921 (11.591, 12.241) | 3.788 (3.289, 4.268) |
| *J* | **-0.003 (-0.070, 0.065)** | -9.588 (-11.087, -8.103) | -22.930 (-23.567, -22.301) | -22.930 (-23.563, -22.299) | -66.112 (-68.079, -64.121) |
| Downsizing SBLC, *k* = 1/16 | | | | | |
| *I* | -2.307 (-2.398, -2.213) | -1.570 (-1.658, -1.485) | -1.570 (-1.655, -1.484) | -1.570 (-1.654, -1.483) | -0.451 (-0.595, -0.312) |
| *H* | -1.142 (-1.210, -1.075) | -0.798 (-0.866, -0.731) | -0.798 (-0.867, -0.730) | -0.798 (-0.866, -0.730) | -0.483 (-0.626, -0.339) |
| $J^1$ | **0.013 (-0.049, 0.075)** | -0.735 (-0.793, -0.675) | -0.735 (-0.794, -0.676) | -0.735 (-0.794, -0.675) | -0.261 (-0.388, -0.134) |
| *J* | **-0.003 (-0.071, 0.063)** | 1.433 (1.374, 1.491) | 1.432 (1.374, 1.488) | 1.432 (1.374, 1.489) | 4.063 (3.950, 4.175) |

Values are mean signed errors, $E_{r,n}^{method} = \hat{A}_{r,n}^{method} - A_{r,n}^{true}$, across 100 independent trait replicates, with bootstrap 95% confidence intervals shown in parentheses. Values are rounded to three decimal places. Assessments of whether bootstrap confidence intervals overlapped zero were based on unrounded confidence limits. Confidence intervals entirely above or below zero indicate detectable overestimation or underestimation, respectively; intervals including zero indicate no detectable directional bias. Boldface highlights estimates whose confidence intervals include zero. GLS-BM and Rphy-BM yielded identical mean signed errors. Small differences in their bootstrap confidence intervals may occur at higher numerical precision because of stochastic variation in bootstrap resampling, but the rounded values reported here are identical. The relative positions of the focal nodes are shown in Figure 1 in the main text, and method abbreviations are defined in the Materials and Methods.

**Table S2. Node-wise signed error after patch correction under representative SBLC scenarios**

| Node | PIC-local | ML-BM | GLS-BM | Rphy-BM | Bayes-MCMC | Stable |
|---|---|---|---|---|---|---|
| Upsizing single-branch large change (SBLC), *k* = 16 | | | | | | |
| *I* | **0.037 (-0.039, 0.115)** | **0.028 (-0.045, 0.102)** | **0.028 (-0.046, 0.102)** | **0.028 (-0.046, 0.101)** | **-0.001 (-0.114, 0.112)** | **0.030 (-0.041, 0.102)** |
| *H* | **0.030 (-0.045, 0.105)** | **0.012 (-0.058, 0.079)** | **0.012 (-0.056, 0.079)** | **0.012 (-0.055, 0.081)** | **-0.011 (-0.106, 0.086)** | **0.003 (-0.065, 0.071)** |
| $J^1$ | **0.013 (-0.049, 0.074)** | **0.011 (-0.045, 0.067)** | **0.011 (-0.045, 0.066)** | **0.011 (-0.046, 0.067)** | **-0.022 (-0.140, 0.094)** | **0.013 (-0.042, 0.069)** |
| *J* | **-0.003 (-0.070, 0.065)** | **-0.003 (-0.071, 0.063)** | **-0.003 (-0.073, 0.064)** | **-0.003 (-0.070, 0.065)** | -0.093 (-0.160, -0.025) | **-0.014 (-0.082, 0.054)** |
| Downsizing SBLC, *k* = 1/16 | | | | | | |
| *I* | **0.037 (-0.036, 0.113)** | **0.028 (-0.045, 0.103)** | **0.028 (-0.045, 0.102)** | **0.028 (-0.046, 0.102)** | **-0.001 (-0.110, 0.110)** | -0.076 (-0.149, -0.001) |
| *H* | **0.030 (-0.047, 0.106)** | **0.012 (-0.055, 0.079)** | **0.012 (-0.056, 0.080)** | **0.012 (-0.056, 0.081)** | **-0.011 (-0.105, 0.087)** | **-0.047 (-0.116, 0.019)** |
| $J^1$ | **0.013 (-0.048, 0.074)** | **0.011 (-0.045, 0.068)** | **0.011 (-0.045, 0.068)** | **0.011 (-0.045, 0.068)** | **-0.022 (-0.138, 0.092)** | **-0.034 (-0.088, 0.021)** |
| *J* | **-0.003 (-0.069, 0.064)** | **-0.003 (-0.073, 0.064)** | **-0.003 (-0.071, 0.063)** | **-0.003 (-0.070, 0.064)** | -0.071 (-0.138, -0.005) | 0.084 (0.014, 0.154) |

Values are mean signed errors after patch correction, $E_{r,n}^{method} = \hat{A}_{r,n}^{method} - A_{r,n}^{true}$, across 100 independent trait replicates, with bootstrap 95% confidence intervals shown in parentheses. The first five columns show signed errors after patch correction for the five conventional ASR methods, whereas the rightmost column shows signed errors from stable-process reconstruction. Values are rounded to three decimal places. Values displayed as 0.000 or -0.000 result from rounding to a limited number of decimal places; assessments of whether bootstrap confidence intervals overlapped zero were based on unrounded confidence limits. Confidence intervals entirely above or below zero indicate detectable overestimation or underestimation, respectively; intervals including zero indicate no clearly detectable directional bias. Boldface highlights estimates whose bootstrap confidence intervals include zero. The corresponding unpatched estimates are reported in Supplementary Table S1, and the before- and after-patching comparison is visualized in Figure 3 of the main text. The stable-process reconstruction provides a model-based benchmark for comparison with patched ASR under the same representative SBLC scenarios. The relative positions of the focal nodes are shown in Figure 1 in the main text, and method abbreviations are defined in the Materials and Methods.

**Table S3. Bootstrap analysis of paired differences in signed error before and after patch correction**

| Node | PIC-local | ML-BM | GLS-BM | Rphy-BM | Bayes-MCMC |
|---|---|---|---|---|---|
| Upsizing single-branch large change (SBLC), *k* = 16 | | | | | |
| *I* | -37.317 (-38.338, -36.290) | -56.175 (-58.665, -53.706) | -25.522 (-26.209, -24.819) | -25.522 (-26.224, -24.814) | -4.598 (-5.191, -4.006) |
| *H* | -18.660 (-19.170, -18.130) | -34.504 (-36.554, -32.481) | -12.932 (-13.278, -12.569) | -12.932 (-13.289, -12.583) | -3.313 (-3.922, -2.737) |
| $J^1$ | **0.000 (0.000, 0.000)** | -54.989 (-57.538, -52.542) | -11.910 (-12.232, -11.571) | -11.910 (-12.237, -11.578) | -3.810 (-4.294, -3.321) |
| *J* | **0.000 (0.000, 0.000)** | 9.584 (8.060, 11.062) | 22.926 (22.284, 23.559) | 22.926 (22.291, 23.555) | 66.019 (64.072, 68.025) |
| Downsizing SBLC, *k* = 1/16 | | | | | |
| *I* | 2.344 (2.271, 2.419) | 1.598 (1.552, 1.644) | 1.598 (1.552, 1.644) | 1.598 (1.552, 1.644) | 0.450 (0.361, 0.541) |
| *H* | 1.171 (1.134, 1.208) | 0.810 (0.787, 0.833) | 0.810 (0.786, 0.833) | 0.810 (0.786, 0.833) | 0.472 (0.386, 0.555) |
| $J^1$ | **0.000 (0.000, 0.000)** | 0.746 (0.724, 0.767) | 0.746 (0.724, 0.767) | 0.746 (0.724, 0.767) | 0.238 (0.167, 0.311) |
| *J* | **0.000 (0.000, 0.000)** | -1.436 (-1.478, -1.394) | -1.435 (-1.476, -1.394) | -1.435 (-1.477, -1.395) | -4.134 (-4.265, -4.004) |

Values are mean paired differences in signed reconstruction error, $\Delta E_{r,n}^{patch} = E_{r,n}^{patch} - E_{r,n}^{conventional}$, across 100 independent trait replicates, with bootstrap 95% confidence intervals shown in parentheses. Values are rounded to three decimal places. Assessments of whether bootstrap confidence intervals overlapped zero were based on unrounded confidence limits. Positive values indicate that patching increased signed error relative to the corresponding unpatched reconstruction, whereas negative values indicate that patching decreased signed error. Confidence intervals entirely above or below zero indicate a statistically detectable patch-induced change in signed error; intervals including zero indicate that the paired change was not clearly distinguishable from zero. Boldface highlights paired differences whose bootstrap confidence intervals include zero. Because D measures the signed change from unpatched to patched reconstructions rather than the signed error itself, the direction of correction should be interpreted together with Supplementary Table S1 and Figure 3 of the main text, where values closer to zero indicate reduced directional reconstruction bias. Focal nodes follow the same notation as in Figure 1 in the main text, and method abbreviations are defined in the Materials and Methods.

**Table S4. Node-wise percentage relative signed error after patch correction under representative SBLC scenarios**

| Node | PIC-local | ML-BM | GLS-BM | Rphy-BM | Bayes-MCMC | Stable |
|---|---|---|---|---|---|---|
| Upsizing single-branch large change (SBLC), *k* = 16 | | | | | | |
| *I* | 1.1% (-0.5%, 2.7%) | 1.2% (-0.3%, 2.8%) | 1.2% (-0.3%, 2.8%) | 1.2% (-0.3%, 2.8%) | 1.4% (-1.0%, 3.8%) | 1.2% (-0.3%, 2.7%) |
| *H* | 0.7% (-0.9%, 2.2%) | 0.6% (-0.7%, 2.0%) | 0.6% (-0.7%, 2.0%) | 0.6% (-0.7%, 2.0%) | 0.6% (-1.4%, 2.6%) | 0.4% (-0.9%, 1.8%) |
| $J^1$ | 0.2% (-1.1%, 1.5%) | 0.5% (-0.7%, 1.6%) | 0.5% (-0.7%, 1.6%) | 0.5% (-0.7%, 1.6%) | 0.9% (-1.5%, 3.3%) | 0.5% (-0.7%, 1.6%) |
| *J* | -0.0% (-0.1%, 0.1%) | -0.0% (-0.1%, 0.1%) | -0.0% (-0.1%, 0.1%) | -0.0% (-0.1%, 0.1%) | -0.1% (-0.2%, -0.0%) | -0.0% (-0.1%, 0.1%) |
| Downsizing SBLC, *k* = 1/16 | | | | | | |
| *I* | 1.1% (-0.5%, 2.7%) | 1.2% (-0.3%, 2.8%) | 1.2% (-0.3%, 2.8%) | 1.2% (-0.3%, 2.8%) | 1.4% (-1.0%, 3.8%) | -0.9% (-2.5%, 0.6%) |
| *H* | 0.7% (-0.9%, 2.2%) | 0.6% (-0.7%, 2.0%) | 0.6% (-0.7%, 2.0%) | 0.6% (-0.7%, 2.0%) | 0.6% (-1.4%, 2.6%) | -0.6% (-1.9%, 0.8%) |
| $J^1$ | 0.2% (-1.1%, 1.5%) | 0.5% (-0.7%, 1.6%) | 0.5% (-0.7%, 1.6%) | 0.5% (-0.7%, 1.6%) | 0.9% (-1.5%, 3.3%) | -0.5% (-1.6%, 0.7%) |
| *J* | 199.8% (-142.9%, 786.1%) | 199.8% (-142.9%, 786.1%) | 199.8% (-142.9%, 786.1%) | 199.8% (-142.9%, 786.1%) | 204.5% (-176.3%, 856.7%) | 216.0% (-104.4%, 768.1%) |

Values are mean relative signed errors, $R_v = E_v / x_v$, expressed as percentages across 100 independent trait replicates, with bootstrap 95% confidence intervals in parentheses, where $E_v$ is signed reconstruction error and $x_v$ is the true ancestral state at node *v*. Positive and negative values indicate overestimation and underestimation, respectively. The first five columns show results after patch correction, and the Stable column shows stable-process reconstruction as an external benchmark. Values are rounded to one decimal percentage point; 0.0% and −0.0% therefore represent small positive and negative values, respectively. Asterisks indicate bootstrap confidence intervals that did not include zero based on unrounded limits. Corresponding unscaled signed errors are reported in Table S2. The large relative errors at node *J* under the downsizing SBLC (*k* = 1/16) are primarily a denominator effect: approximately 17% of replicates had ($|x_J| < 0.1$), causing percentage errors and their confidence intervals to be strongly amplified. These values should therefore not be interpreted as evidence of large reconstruction errors; the unscaled signed errors in Table S2 provide the more appropriate assessment for this case. Focal-node positions are shown in Figure 1 in the main text, and method abbreviations are defined in the Materials and Methods.

#### Table S5. Propagation of SBLC-induced reconstruction errors across phylogenetic distances

| Node | PIC-local unpatched | PIC-local patched | ML-BM unpatched | ML-BM patched | GLS/Rphy-BM unpatched | GLS/Rphy-BM patched | Bayes-MCMC unpatched | Bayes-MCMC patched | Stable |
|---|---|---|---|---|---|---|---|---|---|
| Upsizing SBLC, *k* = 16 | | | | | | | | | |
| *I* | 37.526(26.60,47.38) | -0.016(-0.66,0.593) | 214.361(83.627,325.561) | -0.023(-1.293,1.220) | 25.072(18.034,31.566) | -0.013(-0.648,0.614) | 1.152(-0.037,2.192) | -0.019(-1.218,1.296) | -0.012(-0.648,0.620) |
| *H* | 18.711(12.844,23.794) | -0.054(-0.632,0.494) | 44.689(8.454,127.540) | -0.033(-1.026,0.941) | 12.511(8.712,15.776) | -0.035(-0.595,0.412) | 1.153(0.192,1.923) | -0.032(-0.981,0.918) | -0.036(-0.596,0.408) |
| *G* | 9.360(6.101,12.345) | -0.021(-0.602,0.644) | 3.580(0.482,12.042) | -0.034(-0.858,0.804) | 6.286(4.253,8.224) | -0.021(-0.538,0.400) | 1.130(0.277,1.862) | -0.048(-0.941,0.780) | -0.023(-0.539,0.404) |
| *F* | 4.693(3.091,6.537) | 0.002(-0.602,0.585) | 1.205(0.526,1.919) | -0.014(-0.585,0.639) | 3.225(2.124,4.323) | -0.001(-0.449,0.368) | 1.184(0.392,1.948) | -0.032(-0.648,0.601) | -0.005(-0.477,0.376) |
| *E* | 2.355(1.205,3.239) | 0.010(-0.536,0.553) | 1.162(0.562,1.693) | -0.011(-0.534,0.518) | 1.758(0.959,2.467) | -0.001(-0.442,0.485) | 1.144(0.310,1.914) | -0.012(-0.596,0.548) | 0.006(-0.443,0.512) |
| *D* | 1.153(0.456,1.933) | -0.020(-0.447,0.501) | 1.153(0.454,1.933) | -0.020(-0.453,0.499) | 1.153(0.456,1.933) | -0.020(-0.447,0.501) | 1.091(0.354,1.957) | -0.020(-0.547,0.561) | -0.016(-0.441,0.502) |
| *J* | -0.046(-0.626,0.545) | -0.046(-0.626,0.545) | 195.418(41.660,330.910) | -0.046(-0.626,0.545) | -24.832(-31.425,-16.880) | -0.046(-0.626,0.545) | -73.907(-94.136,-50.187) | -0.086(-0.874,0.774) | -0.047(-0.626,0.544) |
| *K* | -0.039(-0.364,0.322) | -0.039(-0.364,0.322) | 156.716(30.888,267.994) | -0.039(-0.364,0.322) | -12.236(-15.502,-8.305) | -0.039(-0.364,0.322) | -73.901(-94.355,-50.372) | -0.073(-0.562,0.407) | -0.039(-0.357,0.327) |
| *L* | -0.025(-0.247,0.213) | -0.025(-0.247,0.213) | 101.216(9.593,175.988) | -0.025(-0.247,0.213) | -5.926(-7.661,-4.031) | -0.025(-0.247,0.213) | -73.911(-94.754,-50.348) | -0.044(-0.299,0.311) | -0.023(-0.246,0.209) |
| *M* | -0.012(-0.196,0.163) | -0.012(-0.196,0.163) | 58.261(4.686,102.311) | -0.012(-0.196,0.163) | -2.766(-3.538,-1.942) | -0.012(-0.196,0.163) | -73.888(-94.468,-50.499) | -0.022(-0.237,0.215) | -0.011(-0.200,0.162) |
| *N* | -0.004(-0.111,0.100) | -0.004(-0.111,0.100) | 28.557(2.913,52.073) | -0.004(-0.111,0.100) | -1.185(-1.525,-0.771) | -0.004(-0.111,0.100) | -73.880(-94.522,-50.361) | -0.009(-0.149,0.134) | -0.004(-0.112,0.105) |
| *O* | -0.007(-0.070,0.049) | -0.007(-0.070,0.049) | 10.419(-1.194,20.634) | -0.007(-0.070,0.049) | -0.400(-0.531,-0.271) | -0.007(-0.070,0.049) | -73.701(-94.299,-50.208) | -0.010(-0.086,0.058) | -0.007(-0.070,0.048) |
| $J^1$ | -0.020(-0.609,0.552) | -0.020(-0.609,0.552) | 43.624(4.300,130.403) | -0.054(-1.509,1.110) | 12.411(8.753,15.448) | -0.033(-0.568,0.448) | 1.101(0.057,2.193) | -0.049(-1.557,1.152) | -0.033(-0.573,0.445) |
| $J^2$ | 0.009(-0.298,0.339) | 0.009(-0.298,0.339) | -3.837(-7.512,0.572) | 0.065(-1.107,1.232) | 3.124(2.046,4.131) | 0.024(-0.276,0.386) | 1.229(0.089,2.127) | 0.066(-1.121,1.227) | 0.023(-0.277,0.390) |
| $J^3$ | 0.031(-0.224,0.259) | 0.031(-0.224,0.259) | -3.609(-7.861,-0.439) | -0.013(-1.082,1.316) | 0.805(0.396,1.139) | 0.027(-0.229,0.261) | 1.154(0.255,2.208) | -0.024(-1.087,1.284) | 0.027(-0.232,0.261) |
| $J^4$ | 0.016(-0.200,0.248) | 0.016(-0.200,0.248) | -3.684(-6.802,-1.079) | -0.000(-0.816,0.869) | 0.215(0.005,0.454) | 0.016(-0.176,0.240) | 1.151(0.438,2.080) | -0.005(-0.815,0.902) | 0.016(-0.189,0.236) |
| $J^5$ | 0.018(-0.106,0.189) | 0.018(-0.106,0.189) | -3.763(-6.973,-1.141) | -0.006(-0.838,0.751) | 0.072(-0.052,0.229) | 0.018(-0.102,0.184) | 1.165(0.292,1.948) | -0.010(-0.839,0.765) | 0.018(-0.107,0.185) |
| $J^6$ | 0.002(-0.097,0.099) | 0.002(-0.097,0.099) | -3.960(-7.003,-0.990) | 0.021(-0.496,0.589) | 0.020(-0.081,0.120) | 0.002(-0.097,0.100) | 1.195(0.373,1.905) | 0.024(-0.509,0.604) | 0.003(-0.096,0.100) |
| $J^1$ | -0.020(-0.609,0.552) | -0.020(-0.609,0.552) | 43.624(4.300,130.403) | -0.054(-1.509,1.110) | 12.411(8.753,15.448) | -0.033(-0.568,0.448) | 1.101(0.057,2.193) | -0.049(-1.557,1.152) | -0.033(-0.573,0.445) |
| $K^1$ | -0.002(-0.364,0.357) | -0.002(-0.364,0.357) | 0.783(-5.628,12.945) | -0.034(-1.535,1.215) | 6.114(4.320,7.670) | -0.009(-0.332,0.344) | 1.126(-0.214,2.150) | -0.041(-1.534,1.234) | -0.008(-0.328,0.341) |
| $L^1$ | -0.017(-0.212,0.214) | -0.017(-0.212,0.214) | -1.294(-4.061,1.027) | -0.045(-1.639,1.249) | 2.942(2.116,3.691) | -0.020(-0.207,0.187) | 1.106(-0.105,2.197) | -0.060(-1.651,1.258) | -0.020(-0.209,0.187) |
| $M^1$ | -0.004(-0.189,0.184) | -0.004(-0.189,0.184) | -0.799(-2.327,0.177) | -0.026(-1.655,1.304) | 1.377(0.946,1.733) | -0.006(-0.186,0.186) | 1.133(-0.108,2.339) | -0.039(-1.642,1.311) | -0.005(-0.184,0.186) |
| $N^1$ | -0.007(-0.136,0.112) | -0.007(-0.136,0.112) | -0.401(-1.091,0.066) | -0.021(-1.610,1.218) | 0.585(0.383,0.767) | -0.008(-0.131,0.111) | 1.115(-0.217,2.234) | -0.047(-1.681,1.246) | -0.008(-0.129,0.118) |
| $O^1$ | -0.002(-0.057,0.054) | -0.002(-0.057,0.054) | -0.143(-0.468,0.052) | 0.013(-1.042,0.973) | 0.195(0.127,0.271) | -0.003(-0.060,0.053) | 1.021(-0.167,2.088) | -0.038(-1.572,1.258) | -0.002(-0.060,0.054) |
| Downsizing SBLC, *k* = 1/16 | | | | | | | | | |
| *I* | -2.354(-3.309, -1.337) | -0.016(-0.646, 0.593) | -0.098(-1.357, 1.214) | -0.023(-1.293, 1.220) | -1.587(-2.328, -0.672) | -0.013(-0.648, 0.614) | -0.096(-1.473, 1.207) | -0.019(-1.218, 1.296) | -0.091(-0.772, 0.560) |
| *H* | -1.228(-1.778, -0.650) | -0.054(-0.632, 0.494) | -0.106(-1.102, 0.862) | -0.033(-1.026, 0.941) | -0.822(-1.295, -0.300) | -0.035(-0.595, 0.412) | -0.101(-1.147, 0.808) | -0.032(-0.981, 0.918) | -0.075(-0.686, 0.379) |
| *G* | -0.610(-1.150, -0.009) | -0.021(-0.602, 0.644) | -0.108(-0.950, 0.740) | -0.034(-0.858, 0.804) | -0.417(-0.860, -0.002) | -0.021(-0.538, 0.400) | -0.111(-0.970, 0.720) | -0.048(-0.941, 0.780) | -0.041(-0.551, 0.383) |
| *F* | -0.292(-0.834, 0.263) | 0.002(-0.602, 0.585) | -0.087(-0.649, 0.567) | -0.014(-0.585, 0.639) | -0.203(-0.669, 0.176) | -0.001(-0.449, 0.368) | -0.077(-0.733, 0.589) | -0.032(-0.648, 0.601) | -0.011(-0.484, 0.371) |
| *E* | -0.138(-0.685, 0.380) | 0.010(-0.536, 0.553) | -0.085(-0.610, 0.441) | -0.011(-0.534, 0.518) | -0.111(-0.537, 0.375) | -0.001(-0.442, 0.485) | -0.096(-0.620, 0.415) | -0.012(-0.596, 0.548) | -0.006(-0.453, 0.480) |
| *D* | -0.093(-0.522, 0.419) | -0.020(-0.447, 0.501) | -0.093(-0.522, 0.419) | -0.020(-0.453, 0.499) | -0.093(-0.522, 0.419) | -0.020(-0.447, 0.501) | -0.115(-0.608, 0.446) | -0.020(-0.547, 0.561) | -0.023(-0.458, 0.504) |

| Node | PIC-local unpatched | PIC-local patched | ML-BM unpatched | ML-BM patched | GLS/Rphy-BM unpatched | GLS/Rphy-BM patched | Bayes-MCMC unpatched | Bayes-MCMC patched | Stable |
|---|---|---|---|---|---|---|---|---|---|
| $J$ | -0.046(-0.626, 0.545) | -0.046(-0.626, 0.545) | 4.578(3.904, 5.229) | -0.046(-0.626, 0.545) | 1.509(0.990, 2.016) | -0.046(-0.626, 0.545) | 4.591(3.806, 5.403) | -0.066(-0.879, 0.772) | 0.034(-0.568, 0.623) |
| $K$ | -0.039(-0.364, 0.322) | -0.039(-0.364, 0.322) | 4.585(3.745, 5.413) | -0.039(-0.364, 0.322) | 0.726(0.380, 1.065) | -0.039(-0.364, 0.322) | 4.595(3.705, 5.369) | -0.056(-0.521, 0.403) | 0.001(-0.332, 0.355) |
| $L$ | -0.025(-0.247, 0.213) | -0.025(-0.247, 0.213) | 4.599(3.825, 5.488) | -0.025(-0.247, 0.213) | 0.345(0.130, 0.564) | -0.025(-0.247, 0.213) | 4.601(3.810, 5.532) | -0.030(-0.302, 0.337) | -0.004(-0.234, 0.264) |
| $M$ | -0.012(-0.196, 0.163) | -0.012(-0.196, 0.163) | 4.602(3.836, 5.499) | -0.012(-0.196, 0.163) | 0.161(-0.041, 0.323) | -0.012(-0.196, 0.163) | 4.617(3.853, 5.522) | -0.010(-0.224, 0.227) | -0.001(-0.190, 0.194) |
| $N$ | -0.004(-0.111, 0.100) | -0.004(-0.111, 0.100) | 4.386(3.592, 5.224) | -0.004(-0.111, 0.100) | 0.070(-0.037, 0.170) | -0.004(-0.111, 0.100) | 4.616(3.756, 5.499) | 0.002(-0.132, 0.143) | 0.000(-0.108, 0.107) |
| $O$ | -0.007(-0.070, 0.049) | -0.007(-0.070, 0.049) | 1.742(0.693, 3.202) | -0.007(-0.070, 0.049) | 0.018(-0.041, 0.074) | -0.007(-0.070, 0.049) | 4.537(3.633, 5.458) | -0.004(-0.080, 0.066) | -0.006(-0.069, 0.050) |
| $J^1$ | -0.020(-0.609, 0.552) | -0.020(-0.609, 0.552) | -0.131(-1.574, 1.042) | -0.054(-1.509, 1.110) | -0.814(-1.392, -0.309) | -0.033(-0.568, 0.448) | -0.137(-1.460, 1.075) | -0.049(-1.557, 1.152) | -0.072(-0.612, 0.423) |
| $J^2$ | 0.009(-0.298, 0.339) | 0.009(-0.298, 0.339) | -0.002(-1.178, 1.157) | 0.065(-1.107, 1.232) | -0.171(-0.481, 0.190) | 0.024(-0.276, 0.386) | -0.008(-1.223, 1.164) | 0.066(-1.121, 1.227) | 0.013(-0.299, 0.382) |
| $J^3$ | 0.031(-0.224, 0.259) | 0.031(-0.224, 0.259) | -0.087(-1.160, 1.251) | -0.013(-1.082, 1.316) | -0.021(-0.276, 0.224) | 0.027(-0.229, 0.261) | -0.088(-1.153, 1.198) | -0.024(-1.087, 1.284) | 0.025(-0.232, 0.258) |
| $J^4$ | 0.016(-0.200, 0.248) | 0.016(-0.200, 0.248) | -0.076(-0.886, 0.803) | -0.000(-0.816, 0.869) | 0.003(-0.190, 0.227) | 0.016(-0.176, 0.240) | -0.083(-0.912, 0.828) | -0.005(-0.815, 0.902) | 0.015(-0.182, 0.237) |
| $J^5$ | 0.018(-0.106, 0.189) | 0.018(-0.106, 0.189) | -0.081(-0.911, 0.680) | -0.006(-0.838, 0.751) | 0.014(-0.105, 0.181) | 0.018(-0.102, 0.184) | -0.081(-0.909, 0.676) | -0.010(-0.839, 0.765) | 0.017(-0.108, 0.183) |
| $J^6$ | 0.002(-0.097, 0.099) | 0.002(-0.097, 0.099) | -0.051(-0.569, 0.528) | 0.021(-0.496, 0.589) | 0.001(-0.099, 0.098) | 0.002(-0.097, 0.100) | -0.051(-0.562, 0.540) | 0.024(-0.509, 0.604) | 0.002(-0.096, 0.099) |
| $J^1$ | -0.020(-0.609, 0.552) | -0.020(-0.609, 0.552) | -0.131(-1.574, 1.042) | -0.054(-1.509, 1.110) | -0.814(-1.392, -0.309) | -0.033(-0.568, 0.448) | -0.137(-1.460, 1.075) | -0.049(-1.557, 1.152) | -0.072(-0.612, 0.423) |
| $K^1$ | -0.002(-0.364, 0.357) | -0.002(-0.364, 0.357) | -0.114(-1.600, 1.145) | -0.034(-1.535, 1.215) | -0.393(-0.716, -0.058) | -0.009(-0.332, 0.344) | -0.116(-1.591, 1.137) | -0.041(-1.534, 1.234) | -0.028(-0.346, 0.323) |
| $L^1$ | -0.017(-0.212, 0.214) | -0.017(-0.212, 0.214) | -0.129(-1.704, 1.185) | -0.045(-1.639, 1.249) | -0.206(-0.408, 0.007) | -0.020(-0.207, 0.187) | -0.133(-1.727, 1.191) | -0.060(-1.651, 1.258) | -0.029(-0.251, 0.179) |
| $M^1$ | -0.004(-0.189, 0.184) | -0.004(-0.189, 0.184) | -0.116(-1.718, 1.226) | -0.026(-1.655, 1.304) | -0.092(-0.282, 0.089) | -0.006(-0.186, 0.186) | -0.115(-1.697, 1.248) | -0.039(-1.642, 1.311) | -0.010(-0.189, 0.182) |
| $N^1$ | -0.007(-0.136, 0.112) | -0.007(-0.136, 0.112) | -0.115(-1.670, 1.132) | -0.021(-1.610, 1.218) | -0.045(-0.158, 0.078) | -0.008(-0.131, 0.111) | -0.121(-1.732, 1.186) | -0.047(-1.681, 1.246) | -0.010(-0.130, 0.115) |
| $O^1$ | -0.002(-0.057, 0.054) | -0.002(-0.057, 0.054) | -0.062(-0.832, 0.495) | 0.013(-1.042, 0.973) | -0.015(-0.073, 0.040) | -0.003(-0.060, 0.053) | -0.109(-1.645, 1.199) | -0.038(-1.572, 1.258) | -0.003(-0.060, 0.053) |

Values are mean signed errors, $E_{r,n}^{method} = \hat{A}_{r,n}^{method} - A_{r,n}^{true}$, across 100 independent trait replicates, with bootstrap 95% confidence intervals shown in parentheses. Confidence intervals entirely above or below zero indicate detectable overestimation or underestimation, respectively; intervals including zero indicate no detectable directional bias. GLS-BM and Rphy-BM produced identical mean signed errors and are therefore reported together. Their bootstrap confidence limits occasionally differed slightly because resampling indices were generated separately for the two methods; the combined GLS/Rphy-BM columns report the GLS-BM confidence limits. The relative positions of the focal nodes are shown in Figure 5 of the main text and Figure S1, and method abbreviations are defined in the Materials and Methods. Reconstruction errors were examined across six successive node positions in four topological directions relative to the SBLC-bearing *I*–*J* branch: (a) upstream propagation from *I* toward the root (*I*–*D*); (b) downstream propagation from *J* within $T_1$ (*J*–*O*); (c) a depth-controlled lateral series ($J^1$–$J^6$); and (d) progressive penetration into the sister lineage ($J^1$–$O^1$). Node nomenclature and topological relationships follow the scheme illustrated in Figure 1 in the main text.

**Table S6. Node-wise signed ASR errors across post-SBLC evolutionary durations under OU evolution with an upsizing SBLC ($k$ = 16)**

| Node | PIC-local unpatched | PIC-local patched | ML-BM unpatched | ML-BM patched | GLS/Rphy-BM unpatched | GLS/Rphy-BM patched | Bayes-MCMC unpatched | Bayes-MCMC patched | Stable |
|---|---|---|---|---|---|---|---|---|---|
| Long post-SBLC durations in $T_1$, $k$ = 16 | | | | | | | | | |
| $H$ | 6.00(5.74,6.25) | 0.27(0.06,0.47) | 0.17(-0.01,0.35) | 0.15(-0.03,0.32) | 1.66(1.49,1.81) | 0.14(-0.04,0.32) | 0.54(0.36,0.72) | 0.08(-0.11,0.26) | 0.29(0.10,0.48) |
| $I$ | 9.18(8.97,9.41) | 0.14(-0.08,0.35) | 0.07(-0.12,0.27) | 0.03(-0.16,0.23) | 6.72(6.61,6.84) | 0.07(-0.13,0.27) | 0.48(0.29,0.67) | 0.03(-0.18,0.23) | 0.69(0.34,1.06) |
| $J$ | -59.93(-62.54,-57.32) | -59.93(-62.55,-57.29) | -74.12(-77.43,-70.90) | -59.93(-62.55,-57.36) | -66.02(-68.95,-63.25) | -59.93(-62.60,-57.35) | -74.11(-77.31,-70.89) | -60.15(-62.81,-57.54) | -60.49(-63.07,-58.01) |
| $J^1$ | 0.05(-0.10,0.18) | 0.05(-0.09,0.19) | -0.01(-0.18,0.14) | -0.04(-0.20,0.12) | 1.05(0.91,1.18) | 0.04(-0.10,0.17) | 0.36(0.18,0.53) | 0.02(-0.16,0.20) | 0.14(-0.01,0.29) |
| $K^1$ | -9.27(-9.69,-8.87) | -9.27(-9.69,-8.86) | -19.33(-20.30,-18.37) | -9.26(-9.68,-8.85) | -10.16(-10.61,-9.71) | -9.26(-9.68,-8.86) | -23.44(-24.52,-22.43) | -9.27(-9.69,-8.85) | -9.34(-9.75,-8.95) |
| $K^2$ | -4.63(-4.86,-4.40) | -4.63(-4.86,-4.40) | -11.07(-11.77,-10.34) | -4.63(-4.86,-4.40) | -5.07(-5.32,-4.83) | -4.63(-4.87,-4.41) | -18.73(-19.60,-17.92) | -4.65(-4.88,-4.43) | -4.67(-4.89,-4.45) |
| $L^1$ | -0.52(-0.57,-0.48) | -0.52(-0.56,-0.48) | -6.74(-7.30,-6.19) | -0.52(-0.56,-0.47) | -0.54(-0.59,-0.50) | -0.52(-0.56,-0.47) | -14.41(-15.10,-13.72) | -0.53(-0.57,-0.48) | -0.52(-0.56,-0.48) |
| $L^2$ | -5.32(-5.56,-5.08) | -5.32(-5.56,-5.08) | -12.28(-12.94,-11.61) | -5.37(-5.60,-5.13) | -5.78(-6.04,-5.53) | -5.37(-5.60,-5.13) | -19.34(-20.20,-18.49) | -5.30(-5.54,-5.06) | -5.40(-5.64,-5.18) |
| $L^3$ | -3.40(-3.57,-3.23) | -3.40(-3.58,-3.23) | -8.50(-9.04,-7.91) | -3.38(-3.56,-3.21) | -3.65(-3.84,-3.46) | -3.38(-3.55,-3.21) | -17.58(-18.36,-16.82) | -3.44(-3.61,-3.27) | -3.41(-3.58,-3.24) |
| $M$ | -0.27(-0.29,-0.24) | -0.27(-0.29,-0.24) | -1.56(-1.81,-1.31) | -0.27(-0.29,-0.24) | -0.28(-0.30,-0.25) | -0.27(-0.29,-0.24) | -14.04(-14.76,-13.36) | -0.27(-0.29,-0.24) | -0.27(-0.30,-0.24) |
| Intermediate post-SBLC durations in $T_1$, $k$ = 16 | | | | | | | | | |
| $H$ | 5.28(5.10,5.45) | -0.01(-0.19,0.18) | 0.07(-0.13,0.28) | 0.01(-0.20,0.21) | 3.41(3.27,3.55) | -0.00(-0.19,0.19) | 0.13(-0.08,0.34) | 0.22(0.01,0.44) | 0.01(-0.17,0.20) |
| $I$ | 14.47(14.04,14.89) | 0.04(-0.12,0.19) | 0.24(0.06,0.43) | 0.07(-0.12,0.25) | 10.75(10.47,11.01) | 0.04(-0.11,0.20) | 0.21(0.02,0.41) | 0.21(0.02,0.41) | 0.09(-0.07,0.25) |
| $J$ | -43.39(-45.16,-41.63) | -43.39(-45.10,-41.62) | -71.98(-74.90,-69.01) | -43.39(-45.12,-41.58) | -54.43(-56.57,-52.24) | -43.39(-45.12,-41.65) | -73.80(-76.74,-70.81) | -43.47(-45.21,-41.68) | -43.43(-45.25,-41.66) |
| $J^1$ | 0.05(-0.10,0.19) | 0.05(-0.10,0.19) | 0.13(-0.05,0.31) | 0.08(-0.10,0.26) | 6.04(5.87,6.21) | 0.05(-0.09,0.20) | -0.21(-0.40,-0.02) | 0.20(0.01,0.40) | 0.08(-0.07,0.23) |
| $K^1$ | -7.07(-7.36,-6.76) | -7.07(-7.36,-6.77) | -15.48(-16.49,-14.43) | -7.06(-7.35,-6.77) | -8.15(-8.49,-7.82) | -7.06(-7.35,-6.76) | -37.68(-39.18,-36.21) | -7.11(-7.40,-6.81) | -7.07(-7.36,-6.77) |
| $K^2$ | -27.06(-28.13,-25.99) | -27.06(-28.15,-25.95) | -52.17(-54.38,-49.93) | -27.07(-28.15,-25.97) | -32.68(-34.01,-31.40) | -27.07(-28.16,-25.98) | -57.69(-60.02,-55.38) | -27.03(-28.12,-25.92) | -27.10(-28.17,-26.00) |
| $L^1$ | -6.26(-6.53,-5.98) | -6.26(-6.53,-5.98) | -14.88(-15.83,-13.88) | -6.28(-6.55,-6.01) | -7.17(-7.48,-6.87) | -6.28(-6.54,-6.01) | -36.87(-38.34,-35.34) | -6.33(-6.59,-6.06) | -6.28(-6.54,-6.01) |
| $L^2$ | -11.98(-12.47,-11.47) | -11.98(-12.47,-11.48) | -28.05(-29.71,-26.41) | -11.98(-12.48,-11.48) | -13.69(-14.26,-13.14) | -11.98(-12.47,-11.48) | -42.64(-44.36,-40.91) | -11.90(-12.40,-11.39) | -11.99(-12.47,-11.49) |
| $L^3$ | -3.84(-4.01,-3.67) | -3.84(-4.01,-3.68) | -5.83(-6.33,-5.16) | -3.85(-4.03,-3.68) | -4.23(-4.41,-4.04) | -3.85(-4.02,-3.68) | -34.35(-35.77,-32.91) | -3.83(-4.00,-3.66) | -3.85(-4.02,-3.68) |
| $M$ | -4.40(-4.59,-4.20) | -4.40(-4.60,-4.19) | -7.64(-8.22,-6.94) | -4.39(-4.58,-4.19) | -4.82(-5.03,-4.61) | -4.39(-4.58,-4.20) | -34.53(-35.94,-33.12) | -4.44(-4.63,-4.25) | -4.39(-4.59,-4.20) |
| Short post-SBLC durations in $T_1$, $k$ = 16 | | | | | | | | | |
| $H$ | 17.69(17.02,18.38) | 0.05(-0.05,0.14) | 0.46(0.28,0.64) | 0.07(-0.09,0.22) | 12.06(11.63,12.49) | 0.06(-0.03,0.15) | 0.46(0.27,0.65) | 0.16(-0.04,0.34) | 0.07(-0.02,0.15) |
| $I$ | 38.56(37.03,40.09) | 0.01(-0.09,0.10) | 2.92(2.81,3.04) | 0.02(-0.12,0.16) | 20.00(19.26,20.77) | 0.02(-0.07,0.11) | 0.08(-0.13,0.28) | 0.10(-0.08,0.27) | 0.03(-0.07,0.12) |
| $J$ | -5.30(-5.53,-5.07) | -5.30(-5.53,-5.08) | -19.89(-20.97,-18.82) | -5.30(-5.53,-5.07) | -9.57(-9.99,-9.17) | -5.30(-5.53,-5.07) | -74.06(-77.26,-70.89) | -5.31(-5.55,-5.07) | -5.30(-5.53,-5.08) |
| $J^1$ | — | — | — | — | — | — | — | — | — |
| $K^1$ | -4.45(-4.65,-4.26) | -4.45(-4.65,-4.26) | -13.27(-14.14,-12.42) | -4.47(-4.67,-4.28) | -7.29(-7.60,-6.98) | -4.47(-4.67,-4.27) | -72.86(-76.04,-69.75) | -4.49(-4.68,-4.29) | -4.47(-4.67,-4.28) |
| $K^2$ | -4.60(-4.80,-4.40) | -4.60(-4.80,-4.40) | -17.12(-17.96,-16.30) | -4.58(-4.78,-4.37) | -7.79(-8.12,-7.46) | -4.57(-4.78,-4.37) | -72.39(-75.49,-69.30) | -4.56(-4.75,-4.35) | -4.58(-4.78,-4.38) |
| $L^1$ | -3.54(-3.70,-3.39) | -3.54(-3.70,-3.38) | -7.69(-8.44,-6.93) | -3.56(-3.72,-3.40) | -5.32(-5.56,-5.09) | -3.56(-3.72,-3.40) | -71.73(-74.80,-68.74) | -3.58(-3.74,-3.42) | -3.56(-3.72,-3.40) |
| $L^2$ | -1.12(-1.17,-1.07) | -1.12(-1.17,-1.07) | -3.14(-3.92,-2.38) | -1.12(-1.17,-1.07) | -1.50(-1.56,-1.43) | -1.12(-1.17,-1.07) | -69.54(-72.55,-66.52) | -1.13(-1.18,-1.08) | -1.12(-1.17,-1.07) |

| Node | PIC-local unpatched | PIC-local patched | ML-BM unpatched | ML-BM patched | GLS/Rphy-BM unpatched | GLS/Rphy-BM patched | Bayes-MCMC unpatched | Bayes-MCMC patched | Stable |
|---|---|---|---|---|---|---|---|---|---|
| $L^3$ | -2.06(-2.16,-1.97) | -2.06(-2.16,-1.98) | -0.06(-0.76,0.62) | -2.09(-2.18,-1.99) | -2.84(-2.96,-2.71) | -2.09(-2.18,-1.99) | -70.49(-73.66,-67.52) | -2.08(-2.17,-1.99) | -2.09(-2.18,-1.99) |
| *M* | -0.76(-0.80,-0.71) | -0.76(-0.80,-0.71) | -1.97(-2.65,-1.31) | -0.76(-0.80,-0.72) | -0.95(-1.00,-0.90) | -0.76(-0.80,-0.72) | -69.19(-72.23,-66.20) | -0.78(-0.82,-0.74) | -0.76(-0.80,-0.72) |

Values are mean signed errors, calculated as the reconstructed ancestral state minus the true simulated ancestral state, across 100 independent strong-OU trait replicates; bootstrap 95% confidence intervals based on 10,000 resamples are shown in parentheses. Positive and negative values indicate overestimation and underestimation, respectively. Confidence intervals entirely above or below zero indicate detectable directional bias, whereas intervals including zero indicate no detectable directional bias. Results are shown for three fixed *I*–*J* placements on the 4,099-tip empirical mammalian phylogeny, selected to represent long, intermediate, and short post-SBLC durations in $T_1$. *H*, *I*, *J*, and $J^1$ are the focal nodes around the SBLC-bearing *I*–*J* branch. The six internal descendants of *J* are labeled by generation as *K*, *L*, and *M*, with superscripts distinguishing multiple nodes in the same generation. The table lists nodes in the first column, followed by adjacent unpatched and patched results for PIC-local, ML-BM, GLS/Rphy-BM, and Bayes-MCMC; the Stable column provides a stable-process benchmark. GLS-BM and Rphy-BM produced identical mean signed errors and are therefore reported together. Their bootstrap confidence limits occasionally differed slightly because resampling indices were generated separately for the two methods; the combined GLS/Rphy-BM columns report the GLS-BM confidence limits. Values are rounded to two decimal places; 0.00 and -0.00 therefore denote small positive and negative errors, respectively. The corresponding $k$ = 1/16 analyses used paired replicates sharing the same background OU history, SBLC innovation, and downstream random innovations; their results are reported in Repository Appendix A, Figure A8 and Table A1. Node nomenclature follows the selected-tree topology, and method abbreviations are defined in the Materials and Methods.

**Table S7. ASR estimates at candidate SBLC-associated nodes and fossil reference points in cetaceans**

| Method | Treatment | Cut position(s) | Node ID 108 | 110 | 111 | 112 | 149 | 151 | 152 | 153 | 154 | 156 | 157 |
|---|---|---|---|---|---|---|---|---|---|---|---|---|---|
| PIC-local | Unpatched | | 2.80 | 3.24 | 3.65 | 1.93 | 11.59 | 8.57 | 17.82 | 16.25 | 12.04 | 19.31 | 23.84 |
| | Patched | $\varphi$ | 2.80 | 3.24 | 3.65 | 1.93 | **10.82** | **8.57** | **16.25** | **13.64** | **12.04** | 19.31 | 23.84 |
| | | $\psi$ | **2.48** | **2.58** | **2.48** | **1.93** | 11.59 | 8.57 | 17.82 | 16.25 | 12.04 | 19.31 | 23.84 |
| | | $\omega$ | 2.80 | 3.24 | 3.65 | 1.93 | **10.15** | **8.57** | **15.00** | 16.25 | 12.04 | **14.76** | **23.84** |
| | | $\varphi + \omega$ | 2.80 | 3.24 | 3.65 | 1.93 | **8.94** | **8.57** | **12.17** | **12.09** | **12.04** | **12.19** | **23.84** |
| | | $\varphi + \psi + \omega$ | **2.48** | **2.58** | **2.48** | **1.93** | **8.94** | **8.57** | **12.17** | **12.09** | **12.04** | **12.19** | **23.84** |
| ML/GLS/Rphy-BM | Unpatched | | 2.86 | 2.93 | 3.04 | 2.34 | 11.38 | 9.97 | 15.74 | 15.95 | 14.83 | 16.46 | 19.94 |
| | Patched | $\varphi$ | 2.86 | 2.93 | 3.04 | 2.34 | **10.88** | **9.70** | **14.36** | **13.62** | **13.17** | 15.36 | 19.36 |
| | | $\psi$ | **2.57** | **2.49** | **2.43** | **2.11** | 11.38 | 9.97 | 15.74 | 15.95 | 14.83 | 16.46 | 19.94 |
| | | $\omega$ | 2.85 | 2.93 | 3.04 | 2.34 | **10.44** | **9.47** | **13.16** | 14.45 | 13.76 | **13.09** | **23.84** |
| | | $\varphi + \omega$ | 2.85 | 2.93 | 3.04 | 2.34 | **9.66** | **9.05** | **11.02** | **11.35** | **11.54** | **11.14** | **23.84** |
| | | $\varphi + \psi + \omega$ | **2.57** | **2.49** | **2.43** | **2.11** | **9.66** | **9.05** | **11.02** | **11.34** | **11.54** | **11.14** | **23.84** |
| Bayes-MCMC | Unpatched | | 5.85 | 4.55 | 4.71 | 4.46 | 8.13 | 8.63 | 10.22 | 11.55 | 10.85 | 10.57 | 16.23 |
| | Patched | $\varphi$ | 5.44 | 6.58 | 6.08 | 3.45 | **6.35** | **9.03** | **7.05** | **7.71** | **8.11** | 6.63 | 9.46 |
| | | $\psi$ | **6.25** | **4.38** | **4.13** | **2.70** | 7.41 | 8.34 | 10.64 | 10.40 | 10.86 | 11.70 | 15.09 |
| | | $\omega$ | 4.14 | 5.68 | 5.92 | 4.63 | **7.82** | **9.33** | **9.07** | 9.43 | 9.61 | **9.37** | **23.84** |
| | | $\varphi + \omega$ | 6.23 | 5.88 | 5.90 | 4.54 | **8.23** | **7.49** | **9.33** | **9.29** | **9.26** | **9.61** | **23.84** |
| | | $\varphi + \psi + \omega$ | **7.71** | **5.63** | **5.54** | **4.98** | **4.77** | **8.33** | **7.66** | **7.67** | **7.68** | **8.09** | **23.84** |
| Stable | | | 2.40 | 2.34 | 2.33 | 2.16 | 8.97 | 8.69 | 11.93 | 12.04 | 12.09 | 12.06 | 20.68 |
| | Fossil reference(length, m) | | FR6(2.23) | — | — | — | FR14(7.41) | FR15(6.40) | — | — | — | — | — |

Node IDs refer to internal nodes of the 79-tip cetacean molecular phylogeny shown in Figure S16. The displayed nodes were selected because of their proximity to one or more of the three candidate SBLCs identified in Figure S16 or to mapped fossil reference points. Cut positions $\varphi$, $\psi$, and $\omega$ correspond to the branches bearing these three candidate SBLCs and indicate where the phylogeny was divided for patch reconstruction. Combinations indicate simultaneous patching at multiple cut positions; for example, $\varphi + \omega$ denotes joint patching at positions $\varphi$ and $\omega$, whereas $\varphi + \psi + \omega$ denotes joint patching at all three positions. Values in bold type indicate focal nodes associated with the corresponding cut position(s), including the nearest upstream, downstream, and closely adjacent lateral nodes. Values in regular type denote other reconstructed nodes included for comparison, whereas gray shading identifies reference values, including unpatched estimates, stable-process estimates, and fossil-based external references. Fossil reference points are defined in Figure S16 and Supplementary Methods Section S4. FR6, FR14, and FR15 correspond to *Eodelphinus kabatensis*, *Titanocetus sammarinensis*, and *Balaenoptera bertae*, respectively, and provide external references for the focal nodes shown here. Fossil reference values were used only for comparison and were not incorporated as ancestral-state observations in any reconstruction. ASR estimates are shown for unpatched, patched, and stable-process reconstructions. ML-BM, GLS-BM, and Rphy-BM are reported together because their ASR estimates were identical after rounding to two decimal places; differences among the methods appeared only at the third decimal place.
